## Supplementary material for "Polygenic adaptation of rosette growth in *Arabidopsis thaliana*": R Markdown Phenotypic analysis: SupplementaryFile1.html

Polygenic adaptation of rosette growth variation in European Arabidopsis thaliana populations


### Polygenic adaptation of rosette growth variation in European Arabidopsis thaliana populations

###### Benedict Wieters

#### 2020

#### R Markdown

This is an R Markdown document. Markdown is a simple formatting syntax for authoring HTML, PDF, and MS Word documents. For more details on using R Markdown see http://rmarkdown.rstudio.com.

### PART1

##### Loading raw data

##### Growth curve modeling

##### Estimating genotypic means

##### Data overview

Analyses are run separately per treatment, for demonstration I will only run HL completely

##### Separate the experiments

```
HL <- measurements[which(measurements$treatment=="HL"),]
LL <- measurements[which(measurements$treatment=="LL"),]
summary(HL)
```

```
##              Region       Country                  Name            ID     
##  China          : 45   China  : 45   Agu-1           :  3   1002    :  3  
##  Northern_Europe:245   France :145   Ale-Stenar-41-1 :  3   1061    :  3  
##  Spain          :340   Germany: 21   Ale-Stenar-44-4 :  3   1062    :  3  
##  Western_Europe :148   Norway :  3   Ale-Stenar-56-14:  3   1063    :  3  
##                        Spain  :340   Ale-Stenar-64-24:  3   1066    :  3  
##                        Sweden :224   All1-4          :  3   2014-242:  3  
##                                      (Other)         :760   (Other) :760  
##     Latitude       Longitude       treatment   Replicate          tray      
##  Min.   :26.75   Min.   : -7.800   HL:778    Min.   :1.000   Min.   : 1.00  
##  1st Qu.:41.50   1st Qu.: -3.680   LL:  0    1st Qu.:1.000   1st Qu.: 6.00  
##  Median :44.65   Median :  1.743             Median :2.000   Median :12.00  
##  Mean   :46.57   Mean   :  8.551             Mean   :2.001   Mean   :12.48  
##  3rd Qu.:55.69   3rd Qu.: 13.740             3rd Qu.:3.000   3rd Qu.:19.00  
##  Max.   :60.39   Max.   :119.924             Max.   :3.000   Max.   :24.00  
##  NA's   :9       NA's   :9                                                  
##       row            column        Final.Size          t50       
##  Min.   :1.000   Min.   :1.000   Min.   : 4.688   Min.   :13.19  
##  1st Qu.:2.000   1st Qu.:2.000   1st Qu.: 7.959   1st Qu.:17.38  
##  Median :3.000   Median :4.000   Median : 8.652   Median :18.86  
##  Mean   :2.979   Mean   :3.992   Mean   : 8.721   Mean   :19.39  
##  3rd Qu.:4.000   3rd Qu.:6.000   3rd Qu.: 9.475   3rd Qu.:20.88  
##  Max.   :5.000   Max.   :7.000   Max.   :12.304   Max.   :34.81  
##                                                                  
##      Slope            diam04           diam11          diam15     
##  Min.   : 1.988   Min.   :0.2440   Min.   :0.166   Min.   :1.102  
##  1st Qu.: 4.347   1st Qu.:0.3310   1st Qu.:1.045   1st Qu.:1.611  
##  Median : 4.967   Median :0.3440   Median :1.276   Median :1.782  
##  Mean   : 5.199   Mean   :0.3626   Mean   :1.266   Mean   :1.837  
##  3rd Qu.: 5.948   3rd Qu.:0.4200   3rd Qu.:1.500   3rd Qu.:2.317  
##  Max.   :11.069   Max.   :0.4740   Max.   :2.377   Max.   :2.372  
##                   NA's   :773                      NA's   :773    
##      diam18          diam21          diam24        diam25           diam27   
##  Min.   :0.362   Min.   :0.450   Min.   : NA   Min.   : 0.670   Min.   : NA  
##  1st Qu.:2.611   1st Qu.:3.896   1st Qu.: NA   1st Qu.: 6.021   1st Qu.: NA  
##  Median :3.624   Median :5.133   Median : NA   Median : 7.173   Median : NA  
##  Mean   :3.572   Mean   :5.036   Mean   :NaN   Mean   : 7.052   Mean   :NaN  
##  3rd Qu.:4.505   3rd Qu.:6.199   3rd Qu.: NA   3rd Qu.: 8.247   3rd Qu.: NA  
##  Max.   :7.230   Max.   :9.277   Max.   : NA   Max.   :11.842   Max.   : NA  
##  NA's   :2       NA's   :5       NA's   :778   NA's   :5        NA's   :778  
##      diam31        diam32           diam34        diam38        diam39      
##  Min.   : NA   Min.   : 2.468   Min.   : NA   Min.   : NA   Min.   : 4.073  
##  1st Qu.: NA   1st Qu.: 7.573   1st Qu.: NA   1st Qu.: NA   1st Qu.: 7.518  
##  Median : NA   Median : 8.312   Median : NA   Median : NA   Median : 8.220  
##  Mean   :NaN   Mean   : 8.343   Mean   :NaN   Mean   :NaN   Mean   : 8.293  
##  3rd Qu.: NA   3rd Qu.: 9.269   3rd Qu.: NA   3rd Qu.: NA   3rd Qu.: 9.073  
##  Max.   : NA   Max.   :12.301   Max.   : NA   Max.   : NA   Max.   :12.320  
##  NA's   :778                    NA's   :778   NA's   :778   NA's   :2       
##      diam42        diam46           diam47        diam53        diam60   
##  Min.   : NA   Min.   : 4.446   Min.   : NA   Min.   : NA   Min.   : NA  
##  1st Qu.: NA   1st Qu.: 7.351   1st Qu.: NA   1st Qu.: NA   1st Qu.: NA  
##  Median : NA   Median : 8.047   Median : NA   Median : NA   Median : NA  
##  Mean   :NaN   Mean   : 8.140   Mean   :NaN   Mean   :NaN   Mean   :NaN  
##  3rd Qu.: NA   3rd Qu.: 8.926   3rd Qu.: NA   3rd Qu.: NA   3rd Qu.: NA  
##  Max.   : NA   Max.   :11.569   Max.   : NA   Max.   : NA   Max.   : NA  
##  NA's   :778   NA's   :33       NA's   :778   NA's   :778   NA's   :778  
##      diam68        diam74        diam81        diam89       bolting     
##  Min.   : NA   Min.   : NA   Min.   : NA   Min.   : NA   Min.   :25.00  
##  1st Qu.: NA   1st Qu.: NA   1st Qu.: NA   1st Qu.: NA   1st Qu.:38.00  
##  Median : NA   Median : NA   Median : NA   Median : NA   Median :45.00  
##  Mean   :NaN   Mean   :NaN   Mean   :NaN   Mean   :NaN   Mean   :43.17  
##  3rd Qu.: NA   3rd Qu.: NA   3rd Qu.: NA   3rd Qu.: NA   3rd Qu.:49.00  
##  Max.   : NA   Max.   : NA   Max.   : NA   Max.   : NA   Max.   :56.00  
##  NA's   :778   NA's   :778   NA's   :778   NA's   :778   NA's   :442    
##    flowering    
##  Min.   :28.00  
##  1st Qu.:40.00  
##  Median :46.00  
##  Mean   :45.37  
##  3rd Qu.:52.00  
##  Max.   :59.00  
##  NA's   :482
```

```
summary(LL)
```

```
##              Region       Country                 Name           ID     
##  China          : 63   China  : 63   Ctl-62         :  4   Ctl-62 :  4  
##  Northern_Europe:251   France :155   Scg-77         :  4   Scg-77 :  4  
##  Spain          :339   Germany: 18   Agu-1          :  3   1002   :  3  
##  Western_Europe :158   Norway :  3   Ahthx-23       :  3   1061   :  3  
##                        Spain  :339   Ale-Stenar-41-1:  3   1062   :  3  
##                        Sweden :233   Ale-Stenar-44-4:  3   1063   :  3  
##                                      (Other)        :791   (Other):791  
##     Latitude       Longitude       treatment   Replicate          tray     
##  Min.   :26.75   Min.   : -7.800   HL:  0    Min.   :1.000   Min.   : 1.0  
##  1st Qu.:41.32   1st Qu.: -3.575   LL:811    1st Qu.:1.000   1st Qu.: 7.0  
##  Median :44.65   Median :  2.170             Median :2.000   Median :12.0  
##  Mean   :46.18   Mean   : 11.920             Mean   :2.001   Mean   :12.5  
##  3rd Qu.:55.67   3rd Qu.: 13.848             3rd Qu.:3.000   3rd Qu.:18.5  
##  Max.   :60.39   Max.   :119.924             Max.   :3.000   Max.   :24.0  
##                                                                            
##       row            column        Final.Size         t50       
##  Min.   :1.000   Min.   :1.000   Min.   : 8.08   Min.   :25.75  
##  1st Qu.:2.000   1st Qu.:2.000   1st Qu.:13.42   1st Qu.:31.01  
##  Median :3.000   Median :4.000   Median :14.54   Median :33.19  
##  Mean   :2.988   Mean   :4.011   Mean   :14.54   Mean   :33.89  
##  3rd Qu.:4.000   3rd Qu.:6.000   3rd Qu.:15.65   3rd Qu.:36.41  
##  Max.   :5.000   Max.   :7.000   Max.   :22.42   Max.   :50.27  
##                                                                 
##      Slope           diam04        diam11        diam15        diam18   
##  Min.   :2.156   Min.   : NA   Min.   : NA   Min.   : NA   Min.   : NA  
##  1st Qu.:4.957   1st Qu.: NA   1st Qu.: NA   1st Qu.: NA   1st Qu.: NA  
##  Median :5.439   Median : NA   Median : NA   Median : NA   Median : NA  
##  Mean   :5.472   Mean   :NaN   Mean   :NaN   Mean   :NaN   Mean   :NaN  
##  3rd Qu.:5.972   3rd Qu.: NA   3rd Qu.: NA   3rd Qu.: NA   3rd Qu.: NA  
##  Max.   :9.073   Max.   : NA   Max.   : NA   Max.   : NA   Max.   : NA  
##                  NA's   :811   NA's   :811   NA's   :811   NA's   :811  
##      diam21        diam24          diam25        diam27          diam31      
##  Min.   : NA   Min.   :0.556   Min.   : NA   Min.   :0.813   Min.   : 1.518  
##  1st Qu.: NA   1st Qu.:2.198   1st Qu.: NA   1st Qu.:3.062   1st Qu.: 4.434  
##  Median : NA   Median :2.739   Median : NA   Median :3.647   Median : 5.568  
##  Mean   :NaN   Mean   :2.779   Mean   :NaN   Mean   :3.767   Mean   : 5.664  
##  3rd Qu.: NA   3rd Qu.:3.313   3rd Qu.: NA   3rd Qu.:4.449   3rd Qu.: 6.843  
##  Max.   : NA   Max.   :6.151   Max.   : NA   Max.   :7.970   Max.   :10.914  
##  NA's   :811                   NA's   :811                                   
##      diam32        diam34           diam38           diam39        diam42      
##  Min.   : NA   Min.   : 1.985   Min.   : 2.640   Min.   : NA   Min.   : 3.649  
##  1st Qu.: NA   1st Qu.: 5.942   1st Qu.: 7.814   1st Qu.: NA   1st Qu.: 9.661  
##  Median : NA   Median : 7.307   Median : 9.237   Median : NA   Median :11.052  
##  Mean   :NaN   Mean   : 7.256   Mean   : 9.126   Mean   :NaN   Mean   :10.856  
##  3rd Qu.: NA   3rd Qu.: 8.624   3rd Qu.:10.601   3rd Qu.: NA   3rd Qu.:12.217  
##  Max.   : NA   Max.   :12.276   Max.   :14.098   Max.   : NA   Max.   :16.508  
##  NA's   :811                                     NA's   :811                   
##      diam46        diam47           diam53           diam60     
##  Min.   : NA   Min.   : 4.965   Min.   : 7.068   Min.   : 7.48  
##  1st Qu.: NA   1st Qu.:11.233   1st Qu.:12.753   1st Qu.:12.79  
##  Median : NA   Median :12.616   Median :13.944   Median :14.02  
##  Mean   :NaN   Mean   :12.391   Mean   :13.830   Mean   :13.95  
##  3rd Qu.: NA   3rd Qu.:13.685   3rd Qu.:15.046   3rd Qu.:15.16  
##  Max.   : NA   Max.   :17.452   Max.   :18.566   Max.   :20.27  
##  NA's   :811                                     NA's   :58     
##      diam68           diam74          diam81           diam89     
##  Min.   : 7.494   Min.   : 8.00   Min.   : 7.964   Min.   : 7.74  
##  1st Qu.:13.093   1st Qu.:12.98   1st Qu.:12.859   1st Qu.:12.68  
##  Median :14.175   Median :14.21   Median :14.086   Median :13.91  
##  Mean   :14.187   Mean   :14.14   Mean   :14.012   Mean   :13.89  
##  3rd Qu.:15.277   3rd Qu.:15.20   3rd Qu.:15.183   3rd Qu.:15.16  
##  Max.   :19.327   Max.   :19.89   Max.   :19.632   Max.   :20.19  
##  NA's   :33       NA's   :62      NA's   :87       NA's   :184    
##     bolting        flowering    
##  Min.   :31.00   Min.   :38.00  
##  1st Qu.:59.00   1st Qu.:63.00  
##  Median :66.00   Median :73.00  
##  Mean   :67.68   Mean   :70.93  
##  3rd Qu.:81.00   3rd Qu.:81.00  
##  Max.   :90.00   Max.   :90.00  
##  NA's   :447     NA's   :499
```

##### Diameter analysis for growth curves --> re-roganise the data

```
library(reshape2)
library(tidyr)
```

```
## 
## Attaching package: 'tidyr'
```

```
## The following object is masked from 'package:reshape2':
## 
##     smiths
```

```
#subset the needed columns
HLdiam <- HL[,c("Country","Region", "Name", "ID","Latitude", "Longitude","treatment", "Replicate", "tray", "row", "column", "diam04", "diam11", "diam15", "diam18", "diam21", "diam25", "diam32", "diam39","diam46", "bolting", "flowering")]
HLdiam$Plant <- paste(HLdiam$ID,HLdiam$Replicate,sep="_")#create Plant identifier
head(HLdiam)
```

```
##   Country          Region            Name  ID Latitude Longitude treatment
## 1  Sweden Northern_Europe Ale-Stenar-41-1 991  55.3833     14.05        HL
## 2  Sweden Northern_Europe Ale-Stenar-41-1 991  55.3833     14.05        HL
## 3  Sweden Northern_Europe Ale-Stenar-41-1 991  55.3833     14.05        HL
## 4  Sweden Northern_Europe Ale-Stenar-44-4 992  55.3833     14.05        HL
## 5  Sweden Northern_Europe Ale-Stenar-44-4 992  55.3833     14.05        HL
## 6  Sweden Northern_Europe Ale-Stenar-44-4 992  55.3833     14.05        HL
##   Replicate tray row column diam04 diam11 diam15 diam18 diam21 diam25 diam32
## 1         1    2   5      7     NA  1.312     NA  5.051  7.020  9.777  9.933
## 2         2   13   4      1     NA  1.762     NA  5.054  7.893 10.282 10.290
## 3         3   23   1      3     NA  1.657     NA  5.188  7.766  9.577  9.714
## 4         1    2   5      2     NA  1.625     NA  4.519  6.179  7.535  7.801
## 5         2   13   2      2     NA  1.573     NA  4.108  4.340  7.614  8.407
## 6         3   19   5      3     NA  1.009     NA  3.517  5.078  7.340  7.973
##   diam39 diam46 bolting flowering Plant
## 1  8.189  9.829      NA        NA 991_1
## 2  9.765  9.261      NA        NA 991_2
## 3  9.337  9.431      NA        NA 991_3
## 4  8.104  8.084      NA        NA 992_1
## 5  7.893  7.204      NA        NA 992_2
## 6  8.082  7.646      NA        NA 992_3
```

```
LLdiam <- LL[,c("Country","Region", "Name", "ID","Latitude", "Longitude","treatment", "Replicate", "tray", "row", "column", "diam24", "diam27", "diam31", "diam34", "diam38", "diam42", "diam47", "diam53","diam60","diam68","diam74","diam81","diam89", "bolting", "flowering")]
LLdiam$Plant <- paste(LLdiam$ID,LLdiam$Replicate,sep="_")
head(LLdiam)##only TPs with measured diameters are included per treatment
```

```
##     Country          Region            Name  ID Latitude Longitude treatment
## 779  Sweden Northern_Europe Ale-Stenar-41-1 991  55.3833     14.05        LL
## 780  Sweden Northern_Europe Ale-Stenar-41-1 991  55.3833     14.05        LL
## 781  Sweden Northern_Europe Ale-Stenar-41-1 991  55.3833     14.05        LL
## 782  Sweden Northern_Europe Ale-Stenar-44-4 992  55.3833     14.05        LL
## 783  Sweden Northern_Europe Ale-Stenar-44-4 992  55.3833     14.05        LL
## 784  Sweden Northern_Europe Ale-Stenar-44-4 992  55.3833     14.05        LL
##     Replicate tray row column diam24 diam27 diam31 diam34 diam38 diam42 diam47
## 779         1    7   2      1  4.253  6.351  7.621 10.453 11.423 12.632 13.140
## 780         2   13   4      1  3.514  4.867  5.037  6.769 10.368 12.336 12.972
## 781         3   23   1      3  3.453  4.437  6.372  9.153 10.281 13.099 14.098
## 782         1    5   5      4  2.557  2.918  4.941  5.626  8.512  9.533 11.260
## 783         2   13   2      2  3.818  4.690  6.990  8.430 11.600 12.820 13.421
## 784         3   19   5      3  3.141  3.192  5.696  7.529  8.219 11.398 11.666
##     diam53 diam60 diam68 diam74 diam81 diam89 bolting flowering Plant
## 779 13.684 13.219 13.396 12.965 12.772 12.884      NA        NA 991_1
## 780 14.748 13.574 15.273 15.162 15.048 14.355      NA        NA 991_2
## 781 16.135 15.439 15.544 15.609 13.810 16.039      NA        NA 991_3
## 782 15.484 15.817 14.840 15.346 14.799 13.953      NA        NA 992_1
## 783 13.586 13.664 14.399 14.641 13.956 14.376      NA        NA 992_2
## 784 13.766 13.773 14.262 14.161 15.083 13.021      NA        NA 992_3
```

```
###melt the data -> each datapoint seperated and connected to plant
HLdiam.melted <- melt(HLdiam, id=c("Plant","Country","Region", "Name", "ID", "Latitude", "Longitude","treatment", "Replicate", "tray", "row", "column", "bolting", "flowering"), na.rm=T)
head(HLdiam.melted)
```

```
##        Plant Country          Region            Name     ID Latitude Longitude
## 762 Jnf-27_1   China           China          Jnf-27 Jnf-27  26.9860   116.242
## 763 Jnf-27_2   China           China          Jnf-27 Jnf-27  26.9860   116.242
## 764 Jnf-27_3   China           China          Jnf-27 Jnf-27  26.9860   116.242
## 777 Zla-62_2   China           China          Zla-62 Zla-62  30.0540   119.924
## 778 Zla-62_3   China           China          Zla-62 Zla-62  30.0540   119.924
## 779    991_1  Sweden Northern_Europe Ale-Stenar-41-1    991  55.3833    14.050
##     treatment Replicate tray row column bolting flowering variable value
## 762        HL         1    4   1      1      35        38   diam04 0.420
## 763        HL         2    9   2      3      38        40   diam04 0.244
## 764        HL         3   21   1      6      35        40   diam04 0.344
## 777        HL         2   11   3      5      42        47   diam04 0.331
## 778        HL         3   23   5      2      42        49   diam04 0.474
## 779        HL         1    2   5      7      NA        NA   diam11 1.312
```

```
str(HLdiam.melted)
```

```
## 'data.frame':    5409 obs. of  16 variables:
##  $ Plant    : chr  "Jnf-27_1" "Jnf-27_2" "Jnf-27_3" "Zla-62_2" ...
##  $ Country  : Factor w/ 6 levels "China","France",..: 1 1 1 1 1 6 6 6 6 6 ...
##  $ Region   : Factor w/ 4 levels "China","Northern_Europe",..: 1 1 1 1 1 2 2 2 2 2 ...
##  $ Name     : Factor w/ 284 levels "Agu-1","Ahthx-23",..: 160 160 160 284 284 3 3 3 4 4 ...
##  $ ID       : Factor w/ 280 levels "1002","1061",..: 272 272 272 280 280 249 249 249 250 250 ...
##  $ Latitude : num  27 27 27 30.1 30.1 ...
##  $ Longitude: num  116 116 116 120 120 ...
##  $ treatment: Factor w/ 2 levels "HL","LL": 1 1 1 1 1 1 1 1 1 1 ...
##  $ Replicate: int  1 2 3 2 3 1 2 3 1 2 ...
##  $ tray     : int  4 9 21 11 23 2 13 23 2 13 ...
##  $ row      : int  1 2 1 3 5 5 4 1 5 2 ...
##  $ column   : int  1 3 6 5 2 7 1 3 2 2 ...
##  $ bolting  : int  35 38 35 42 42 NA NA NA NA NA ...
##  $ flowering: int  38 40 40 47 49 NA NA NA NA NA ...
##  $ variable : Factor w/ 9 levels "diam04","diam11",..: 1 1 1 1 1 2 2 2 2 2 ...
##  $ value    : num  0.42 0.244 0.344 0.331 0.474 ...
```

```
#separate phenotype and timepoint by the m in diam -> time axis
HLdiam.melted <- separate(HLdiam.melted, variable, c("phenotype", "time"), sep = "m")
HLdiam.melted$time <- as.numeric(HLdiam.melted$time)
HLdiam.melted$phenotype <- NULL 
head(HLdiam.melted)
```

```
##        Plant Country          Region            Name     ID Latitude Longitude
## 762 Jnf-27_1   China           China          Jnf-27 Jnf-27  26.9860   116.242
## 763 Jnf-27_2   China           China          Jnf-27 Jnf-27  26.9860   116.242
## 764 Jnf-27_3   China           China          Jnf-27 Jnf-27  26.9860   116.242
## 777 Zla-62_2   China           China          Zla-62 Zla-62  30.0540   119.924
## 778 Zla-62_3   China           China          Zla-62 Zla-62  30.0540   119.924
## 779    991_1  Sweden Northern_Europe Ale-Stenar-41-1    991  55.3833    14.050
##     treatment Replicate tray row column bolting flowering time value
## 762        HL         1    4   1      1      35        38    4 0.420
## 763        HL         2    9   2      3      38        40    4 0.244
## 764        HL         3   21   1      6      35        40    4 0.344
## 777        HL         2   11   3      5      42        47    4 0.331
## 778        HL         3   23   5      2      42        49    4 0.474
## 779        HL         1    2   5      7      NA        NA   11 1.312
```

```
###LL
LLdiam.melted <- melt(LLdiam, id=c("Plant","Country","Region", "Name", "ID", "Latitude", "Longitude","treatment", "Replicate", "tray", "row", "column", "bolting", "flowering"), na.rm=T)
head(LLdiam.melted)
```

```
##   Plant Country          Region            Name  ID Latitude Longitude
## 1 991_1  Sweden Northern_Europe Ale-Stenar-41-1 991  55.3833     14.05
## 2 991_2  Sweden Northern_Europe Ale-Stenar-41-1 991  55.3833     14.05
## 3 991_3  Sweden Northern_Europe Ale-Stenar-41-1 991  55.3833     14.05
## 4 992_1  Sweden Northern_Europe Ale-Stenar-44-4 992  55.3833     14.05
## 5 992_2  Sweden Northern_Europe Ale-Stenar-44-4 992  55.3833     14.05
## 6 992_3  Sweden Northern_Europe Ale-Stenar-44-4 992  55.3833     14.05
##   treatment Replicate tray row column bolting flowering variable value
## 1        LL         1    7   2      1      NA        NA   diam24 4.253
## 2        LL         2   13   4      1      NA        NA   diam24 3.514
## 3        LL         3   23   1      3      NA        NA   diam24 3.453
## 4        LL         1    5   5      4      NA        NA   diam24 2.557
## 5        LL         2   13   2      2      NA        NA   diam24 3.818
## 6        LL         3   19   5      3      NA        NA   diam24 3.141
```

```
str(LLdiam.melted)
```

```
## 'data.frame':    10119 obs. of  16 variables:
##  $ Plant    : chr  "991_1" "991_2" "991_3" "992_1" ...
##  $ Country  : Factor w/ 6 levels "China","France",..: 6 6 6 6 6 6 6 6 6 6 ...
##  $ Region   : Factor w/ 4 levels "China","Northern_Europe",..: 2 2 2 2 2 2 2 2 2 2 ...
##  $ Name     : Factor w/ 284 levels "Agu-1","Ahthx-23",..: 3 3 3 4 4 4 5 5 5 6 ...
##  $ ID       : Factor w/ 280 levels "1002","1061",..: 249 249 249 250 250 250 258 258 258 1 ...
##  $ Latitude : num  55.4 55.4 55.4 55.4 55.4 ...
##  $ Longitude: num  14.1 14.1 14.1 14.1 14.1 ...
##  $ treatment: Factor w/ 2 levels "HL","LL": 2 2 2 2 2 2 2 2 2 2 ...
##  $ Replicate: int  1 2 3 1 2 3 1 2 3 1 ...
##  $ tray     : int  7 13 23 5 13 19 4 16 23 2 ...
##  $ row      : int  2 4 1 5 2 5 2 3 2 2 ...
##  $ column   : int  1 1 3 4 2 3 6 1 6 1 ...
##  $ bolting  : int  NA NA NA NA NA NA NA 52 NA NA ...
##  $ flowering: int  NA NA NA NA NA NA NA 81 NA NA ...
##  $ variable : Factor w/ 13 levels "diam24","diam27",..: 1 1 1 1 1 1 1 1 1 1 ...
##  $ value    : num  4.25 3.51 3.45 2.56 3.82 ...
```

```
#separate phenotype and timepoint by the m in diam -> time axis
LLdiam.melted <- separate(LLdiam.melted, variable, c("phenotype", "time"), sep = "m")
LLdiam.melted$time <- as.numeric(LLdiam.melted$time)
LLdiam.melted$phenotype <- NULL 
head(LLdiam.melted)
```

```
##   Plant Country          Region            Name  ID Latitude Longitude
## 1 991_1  Sweden Northern_Europe Ale-Stenar-41-1 991  55.3833     14.05
## 2 991_2  Sweden Northern_Europe Ale-Stenar-41-1 991  55.3833     14.05
## 3 991_3  Sweden Northern_Europe Ale-Stenar-41-1 991  55.3833     14.05
## 4 992_1  Sweden Northern_Europe Ale-Stenar-44-4 992  55.3833     14.05
## 5 992_2  Sweden Northern_Europe Ale-Stenar-44-4 992  55.3833     14.05
## 6 992_3  Sweden Northern_Europe Ale-Stenar-44-4 992  55.3833     14.05
##   treatment Replicate tray row column bolting flowering time value
## 1        LL         1    7   2      1      NA        NA   24 4.253
## 2        LL         2   13   4      1      NA        NA   24 3.514
## 3        LL         3   23   1      3      NA        NA   24 3.453
## 4        LL         1    5   5      4      NA        NA   24 2.557
## 5        LL         2   13   2      2      NA        NA   24 3.818
## 6        LL         3   19   5      3      NA        NA   24 3.141
```

```
###HL/LLdiam.melted gives info about each individual PLANT
#write tables:
#write.table(HLdiam.melted, file="HLdiam.melted.txt")
#write.table(LLdiam.melted, file="LLdiam.melted.txt")
#####starting from here I only run HL for demonstration
```

### Logistic growth rate --- Fitting of the growth response curves

```
##for every single individual
library(drc)
```

```
## Loading required package: MASS
```

```
## 
## 'drc' has been loaded.
```

```
## Please cite R and 'drc' if used for a publication,
```

```
## for references type 'citation()' and 'citation('drc')'.
```

```
## 
## Attaching package: 'drc'
```

```
## The following objects are masked from 'package:stats':
## 
##     gaussian, getInitial
```

```
##HL
plants <- unique(HLdiam.melted$Plant) #extract indivualnumbers
plants <- sort(plants)
dHLnew <- HLdiam.melted[order(HLdiam.melted$Plant),]
head(dHLnew)
```

```
##       Plant Country          Region             Name   ID Latitude Longitude
## 788  1002_1  Sweden Northern_Europe Ale-Stenar-64-24 1002  55.3833     14.05
## 2344 1002_1  Sweden Northern_Europe Ale-Stenar-64-24 1002  55.3833     14.05
## 3122 1002_1  Sweden Northern_Europe Ale-Stenar-64-24 1002  55.3833     14.05
## 3900 1002_1  Sweden Northern_Europe Ale-Stenar-64-24 1002  55.3833     14.05
## 4678 1002_1  Sweden Northern_Europe Ale-Stenar-64-24 1002  55.3833     14.05
## 5456 1002_1  Sweden Northern_Europe Ale-Stenar-64-24 1002  55.3833     14.05
##      treatment Replicate tray row column bolting flowering time value
## 788         HL         1    6   2      5      NA        NA   11 1.523
## 2344        HL         1    6   2      5      NA        NA   18 2.266
## 3122        HL         1    6   2      5      NA        NA   21 3.853
## 3900        HL         1    6   2      5      NA        NA   25 5.521
## 4678        HL         1    6   2      5      NA        NA   32 6.786
## 5456        HL         1    6   2      5      NA        NA   39 6.760
```

```
grHL <- list()
#########growth-curve creation and extraction of parameters for individual plants
for(j in plants){
  cat("Running HL:",j,"\n")
  df <- dHLnew[dHLnew$Plant%in%j, ]
  dfnew <- df[order(df$Plant),]
  grHL[[j]] <- drm(value~time, fct=LL.4(fixed=c(NA,0,NA,NA), names=c("Slope","0",  "FS", "t50")), data = dfnew, na.action = na.omit)
}
```

```
## Running HL: 1002_1 
## Running HL: 1002_2 
## Running HL: 1002_3 
## Running HL: 1061_1 
## Running HL: 1061_2 
## Running HL: 1061_3 
## Running HL: 1062_1 
## Running HL: 1062_2 
## Running HL: 1062_3 
## Running HL: 1063_1 
## Running HL: 1063_2 
## Running HL: 1063_3 
## Running HL: 1066_1 
## Running HL: 1066_2 
## Running HL: 1066_3 
## Running HL: 2014-242_1 
## Running HL: 2014-242_2 
## Running HL: 2014-242_3 
## Running HL: 2014-245_1 
## Running HL: 2014-245_2 
## Running HL: 2014-245_3 
## Running HL: 2014-249_1 
## Running HL: 2014-249_2 
## Running HL: 2014-249_3 
## Running HL: 2014-250_1 
## Running HL: 2014-250_2 
## Running HL: 2014-250_3 
## Running HL: 2014-251_1 
## Running HL: 2014-251_2 
## Running HL: 2014-252_1 
## Running HL: 2014-252_2 
## Running HL: 2014-252_3 
## Running HL: 2014-253_1 
## Running HL: 2014-253_2 
## Running HL: 2014-253_3 
## Running HL: 2014-255_1 
## Running HL: 2014-255_2 
## Running HL: 2014-255_3 
## Running HL: 2014-256_1 
## Running HL: 2014-256_2 
## Running HL: 2014-256_3 
## Running HL: 2014-258_1 
## Running HL: 2014-258_2 
## Running HL: 2014-258_3 
## Running HL: 2014-260_1 
## Running HL: 2014-260_2 
## Running HL: 2014-260_3 
## Running HL: 2014-261_1 
## Running HL: 2014-261_3 
## Running HL: 2014-262_1 
## Running HL: 2014-262_2 
## Running HL: 2014-262_3 
## Running HL: 2014-264_1 
## Running HL: 2014-264_2 
## Running HL: 2014-264_3 
## Running HL: 2014-265_1 
## Running HL: 2014-265_2 
## Running HL: 2014-265_3 
## Running HL: 2014-266_1 
## Running HL: 2014-266_2 
## Running HL: 2014-266_3 
## Running HL: 2014-268_1 
## Running HL: 2014-268_3 
## Running HL: 2014-269_1 
## Running HL: 2014-269_2 
## Running HL: 2014-269_3 
## Running HL: 2014-270_1 
## Running HL: 2014-270_2 
## Running HL: 2014-270_3 
## Running HL: 2014-272_1 
## Running HL: 2014-272_2 
## Running HL: 2014-272_3 
## Running HL: 2014-274_1 
## Running HL: 2014-274_2 
## Running HL: 2014-274_3 
## Running HL: 2014-275_1 
## Running HL: 2014-275_2 
## Running HL: 2014-275_3 
## Running HL: 2014-276_1 
## Running HL: 2014-276_2 
## Running HL: 2014-276_3 
## Running HL: 2014-277_1 
## Running HL: 2014-277_2 
## Running HL: 2014-277_3 
## Running HL: 2014-278_1 
## Running HL: 2014-278_2 
## Running HL: 2014-278_3 
## Running HL: 2014-281_1 
## Running HL: 2014-281_2 
## Running HL: 2014-281_3 
## Running HL: 2014-282_1 
## Running HL: 2014-282_2 
## Running HL: 2014-284_1 
## Running HL: 2014-284_2 
## Running HL: 2014-284_3 
## Running HL: 2014-285_1 
## Running HL: 2014-285_2 
## Running HL: 2014-285_3 
## Running HL: 2014-286_1 
## Running HL: 2014-286_2 
## Running HL: 2014-286_3 
## Running HL: 2014-293_1 
## Running HL: 2014-293_2 
## Running HL: 2014-293_3 
## Running HL: 2014-296_1 
## Running HL: 2014-296_2 
## Running HL: 2014-296_3 
## Running HL: 2014-298_1 
## Running HL: 2014-298_2 
## Running HL: 2014-298_3 
## Running HL: 2014-299_1 
## Running HL: 2014-299_2 
## Running HL: 2014-299_3 
## Running HL: 2014-300_1 
## Running HL: 2014-300_2 
## Running HL: 2014-300_3 
## Running HL: 2014-306_1 
## Running HL: 2014-306_2 
## Running HL: 2014-306_3 
## Running HL: 2014-307_1 
## Running HL: 2014-307_2 
## Running HL: 2014-307_3 
## Running HL: 2014-309_1 
## Running HL: 2014-309_2 
## Running HL: 2014-309_3 
## Running HL: 2014-311_1 
## Running HL: 2014-311_2 
## Running HL: 2014-311_3 
## Running HL: 2014-316_1 
## Running HL: 2014-316_2 
## Running HL: 2014-317_1 
## Running HL: 2014-317_2 
## Running HL: 2014-321_1 
## Running HL: 2014-321_2 
## Running HL: 2014-321_3 
## Running HL: 2014-324_1 
## Running HL: 2014-324_2 
## Running HL: 2014-324_3 
## Running HL: 2014-325_1 
## Running HL: 2014-325_2 
## Running HL: 2014-325_3 
## Running HL: 2014-328_1 
## Running HL: 2014-328_2 
## Running HL: 2014-328_3 
## Running HL: 2014-332_1 
## Running HL: 2014-332_3 
## Running HL: 2014-333_1 
## Running HL: 2014-333_2 
## Running HL: 2014-333_3 
## Running HL: 2014-336_1 
## Running HL: 2014-336_2 
## Running HL: 2014-336_3 
## Running HL: 5830_1 
## Running HL: 5830_3 
## Running HL: 5831_1 
## Running HL: 5831_2 
## Running HL: 5831_3 
## Running HL: 5832_1 
## Running HL: 6019_1 
## Running HL: 6019_2 
## Running HL: 6019_3 
## Running HL: 6020_1 
## Running HL: 6020_2 
## Running HL: 6020_3 
## Running HL: 6021_1 
## Running HL: 6021_2 
## Running HL: 6021_3 
## Running HL: 6023_1 
## Running HL: 6023_2 
## Running HL: 6023_3 
## Running HL: 6024_1 
## Running HL: 6024_2 
## Running HL: 6024_3 
## Running HL: 6035_1 
## Running HL: 6035_2 
## Running HL: 6035_3 
## Running HL: 6036_1 
## Running HL: 6036_2 
## Running HL: 6036_3 
## Running HL: 6038_1 
## Running HL: 6038_2 
## Running HL: 6038_3 
## Running HL: 6039_1 
## Running HL: 6039_2 
## Running HL: 6039_3 
## Running HL: 6040_1 
## Running HL: 6040_2 
## Running HL: 6040_3 
## Running HL: 6041_1 
## Running HL: 6041_2 
## Running HL: 6041_3 
## Running HL: 6073_1 
## Running HL: 6073_2 
## Running HL: 6073_3 
## Running HL: 6076_1 
## Running HL: 6076_2 
## Running HL: 6076_3 
## Running HL: 6085_1 
## Running HL: 6085_2 
## Running HL: 6085_3 
## Running HL: 6094_1 
## Running HL: 6094_2 
## Running HL: 6094_3 
## Running HL: 6106_1 
## Running HL: 6106_2 
## Running HL: 6106_3 
## Running HL: 6109_1 
## Running HL: 6109_2 
## Running HL: 6109_3 
## Running HL: 6112_1 
## Running HL: 6112_2 
## Running HL: 6112_3 
## Running HL: 6114_1 
## Running HL: 6114_2 
## Running HL: 6114_3 
## Running HL: 6142_1 
## Running HL: 6142_2 
## Running HL: 6142_3 
## Running HL: 6188_1 
## Running HL: 6188_2 
## Running HL: 6188_3 
## Running HL: 6189_1 
## Running HL: 6189_2 
## Running HL: 6189_3 
## Running HL: 6191_1 
## Running HL: 6191_2 
## Running HL: 6191_3 
## Running HL: 6201_1 
## Running HL: 6201_2 
## Running HL: 6201_3 
## Running HL: 6242_1 
## Running HL: 6242_2 
## Running HL: 6242_3 
## Running HL: 6258_1 
## Running HL: 6258_2 
## Running HL: 6258_3 
## Running HL: 6284_1 
## Running HL: 6284_2 
## Running HL: 6284_3 
## Running HL: 6413_1 
## Running HL: 6413_2 
## Running HL: 6413_3 
## Running HL: 6933_1 
## Running HL: 6933_2 
## Running HL: 6933_3 
## Running HL: 6961_1 
## Running HL: 6961_2 
## Running HL: 6961_3 
## Running HL: 6970_1 
## Running HL: 6971_1 
## Running HL: 6971_2 
## Running HL: 6971_3 
## Running HL: 6974_1 
## Running HL: 6974_2 
## Running HL: 6974_3 
## Running HL: 7013_1 
## Running HL: 7013_2 
## Running HL: 7013_3 
## Running HL: 7147_1 
## Running HL: 7147_2 
## Running HL: 7147_3 
## Running HL: 7164_1 
## Running HL: 7164_2 
## Running HL: 7164_3 
## Running HL: 7169_1 
## Running HL: 7169_2 
## Running HL: 7169_3 
## Running HL: 7268_1 
## Running HL: 7268_2 
## Running HL: 7268_3 
## Running HL: 7288_1 
## Running HL: 7288_2 
## Running HL: 7288_3 
## Running HL: 7305_1 
## Running HL: 7305_2 
## Running HL: 7305_3 
## Running HL: 7327_1 
## Running HL: 7327_2 
## Running HL: 7327_3 
## Running HL: 7328_1 
## Running HL: 7328_2 
## Running HL: 7343_1 
## Running HL: 7343_2 
## Running HL: 7343_3 
## Running HL: 7346_1 
## Running HL: 7346_2 
## Running HL: 7346_3 
## Running HL: 7354_1 
## Running HL: 7354_2 
## Running HL: 7354_3 
## Running HL: 8222_1 
## Running HL: 8222_2 
## Running HL: 8222_3 
## Running HL: 8231_1 
## Running HL: 8231_2 
## Running HL: 8231_3 
## Running HL: 8234_1 
## Running HL: 8234_2 
## Running HL: 8234_3 
## Running HL: 8237_1 
## Running HL: 8237_2 
## Running HL: 8237_3 
## Running HL: 8240_1 
## Running HL: 8240_2 
## Running HL: 8240_3 
## Running HL: 8241_1 
## Running HL: 8241_2 
## Running HL: 8241_3 
## Running HL: 8242_1 
## Running HL: 8242_2 
## Running HL: 8242_3 
## Running HL: 8249_1 
## Running HL: 8249_2 
## Running HL: 8249_3 
## Running HL: 8256_1 
## Running HL: 8256_2 
## Running HL: 8256_3 
## Running HL: 8259_2 
## Running HL: 8259_3 
## Running HL: 8264_1 
## Running HL: 8264_2 
## Running HL: 8283_1 
## Running HL: 8283_2 
## Running HL: 8283_3 
## Running HL: 8306_1 
## Running HL: 8306_2 
## Running HL: 8306_3 
## Running HL: 8307_1 
## Running HL: 8307_2 
## Running HL: 8307_3 
## Running HL: 8326_1 
## Running HL: 8326_2 
## Running HL: 8326_3 
## Running HL: 8335_1 
## Running HL: 8335_2 
## Running HL: 8335_3 
## Running HL: 8369_1 
## Running HL: 8369_3 
## Running HL: 8422_1 
## Running HL: 8422_2 
## Running HL: 8422_3 
## Running HL: 8426_1 
## Running HL: 8426_2 
## Running HL: 8426_3 
## Running HL: 9057_1 
## Running HL: 9057_2 
## Running HL: 9057_3 
## Running HL: 9339_1 
## Running HL: 9339_2 
## Running HL: 9339_3 
## Running HL: 9369_1 
## Running HL: 9369_2 
## Running HL: 9369_3 
## Running HL: 9380_1 
## Running HL: 9380_2 
## Running HL: 9380_3 
## Running HL: 9399_1 
## Running HL: 9399_2 
## Running HL: 9404_1 
## Running HL: 9404_2 
## Running HL: 9404_3 
## Running HL: 9405_1 
## Running HL: 9405_2 
## Running HL: 9405_3 
## Running HL: 9407_1 
## Running HL: 9407_2 
## Running HL: 9407_3 
## Running HL: 9408_1 
## Running HL: 9408_2 
## Running HL: 9408_3 
## Running HL: 9409_1 
## Running HL: 9409_2 
## Running HL: 9409_3 
## Running HL: 9421_2 
## Running HL: 9421_3 
## Running HL: 9436_1 
## Running HL: 9436_2 
## Running HL: 9436_3 
## Running HL: 9437_1 
## Running HL: 9437_2 
## Running HL: 9437_3 
## Running HL: 9453_1 
## Running HL: 9453_2 
## Running HL: 9453_3 
## Running HL: 9454_1 
## Running HL: 9454_2 
## Running HL: 9454_3 
## Running HL: 9470_1 
## Running HL: 9470_2 
## Running HL: 9470_3 
## Running HL: 9481_1 
## Running HL: 9481_2 
## Running HL: 9481_3 
## Running HL: 9506_1 
## Running HL: 9506_2 
## Running HL: 9506_3 
## Running HL: 9517_1 
## Running HL: 9517_2 
## Running HL: 9517_3 
## Running HL: 9520_1 
## Running HL: 9520_2 
## Running HL: 9520_3 
## Running HL: 9521_1 
## Running HL: 9521_2 
## Running HL: 9521_3 
## Running HL: 9524_1 
## Running HL: 9524_2 
## Running HL: 9524_3 
## Running HL: 9525_1 
## Running HL: 9525_2 
## Running HL: 9525_3 
## Running HL: 9526_1 
## Running HL: 9526_2 
## Running HL: 9526_3 
## Running HL: 9527_1 
## Running HL: 9527_2 
## Running HL: 9527_3 
## Running HL: 9528_1 
## Running HL: 9528_2 
## Running HL: 9528_3 
## Running HL: 9531_1 
## Running HL: 9531_2 
## Running HL: 9531_3 
## Running HL: 9532_1 
## Running HL: 9532_2 
## Running HL: 9532_3 
## Running HL: 9534_1 
## Running HL: 9534_2 
## Running HL: 9534_3 
## Running HL: 9535_1 
## Running HL: 9535_2 
## Running HL: 9535_3 
## Running HL: 9539_1 
## Running HL: 9539_2 
## Running HL: 9539_3 
## Running HL: 9540_1 
## Running HL: 9540_3 
## Running HL: 9546_1 
## Running HL: 9546_2 
## Running HL: 9546_3 
## Running HL: 9547_1 
## Running HL: 9547_2 
## Running HL: 9547_3 
## Running HL: 9548_1 
## Running HL: 9548_2 
## Running HL: 9548_3 
## Running HL: 9551_1 
## Running HL: 9551_2 
## Running HL: 9551_3 
## Running HL: 9552_1 
## Running HL: 9552_2 
## Running HL: 9552_3 
## Running HL: 9553_1 
## Running HL: 9553_2 
## Running HL: 9553_3 
## Running HL: 9556_1 
## Running HL: 9556_2 
## Running HL: 9556_3 
## Running HL: 9557_1 
## Running HL: 9557_2 
## Running HL: 9557_3 
## Running HL: 9558_1 
## Running HL: 9558_2 
## Running HL: 9558_3 
## Running HL: 9561_1 
## Running HL: 9561_2 
## Running HL: 9561_3 
## Running HL: 9562_1 
## Running HL: 9562_2 
## Running HL: 9562_3 
## Running HL: 9564_1 
## Running HL: 9564_2 
## Running HL: 9564_3 
## Running HL: 9565_1 
## Running HL: 9565_2 
## Running HL: 9565_3 
## Running HL: 9567_1 
## Running HL: 9567_2 
## Running HL: 9567_3 
## Running HL: 9568_1 
## Running HL: 9568_2 
## Running HL: 9568_3 
## Running HL: 9569_1 
## Running HL: 9569_2 
## Running HL: 9569_3 
## Running HL: 9573_1 
## Running HL: 9573_2 
## Running HL: 9573_3 
## Running HL: 9577_1 
## Running HL: 9577_2 
## Running HL: 9577_3 
## Running HL: 9578_1 
## Running HL: 9578_2 
## Running HL: 9578_3 
## Running HL: 9580_2 
## Running HL: 9580_3 
## Running HL: 9581_1 
## Running HL: 9581_2 
## Running HL: 9582_1 
## Running HL: 9582_2 
## Running HL: 9582_3 
## Running HL: 9584_1 
## Running HL: 9584_2 
## Running HL: 9584_3 
## Running HL: 9585_2 
## Running HL: 9585_3 
## Running HL: 9586_1 
## Running HL: 9586_2 
## Running HL: 9586_3 
## Running HL: 9587_1 
## Running HL: 9587_2 
## Running HL: 9587_3 
## Running HL: 9588_1 
## Running HL: 9588_2 
## Running HL: 9588_3 
## Running HL: 9589_1 
## Running HL: 9589_2 
## Running HL: 9589_3 
## Running HL: 9591_2 
## Running HL: 9591_3 
## Running HL: 9592_1 
## Running HL: 9592_2 
## Running HL: 9592_3 
## Running HL: 9593_1 
## Running HL: 9593_2 
## Running HL: 9593_3 
## Running HL: 9594_1 
## Running HL: 9594_2 
## Running HL: 9594_3 
## Running HL: 9595_1 
## Running HL: 9595_2 
## Running HL: 9595_3 
## Running HL: 9596_1 
## Running HL: 9596_2 
## Running HL: 9597_1 
## Running HL: 9597_2 
## Running HL: 9597_3 
## Running HL: 9599_1 
## Running HL: 9599_2 
## Running HL: 9599_3 
## Running HL: 9601_1 
## Running HL: 9601_2 
## Running HL: 9601_3 
## Running HL: 9602_1 
## Running HL: 9602_2 
## Running HL: 9602_3 
## Running HL: 9819_1 
## Running HL: 9819_2 
## Running HL: 9819_3 
## Running HL: 9820_1 
## Running HL: 9820_2 
## Running HL: 9820_3 
## Running HL: 9821_1 
## Running HL: 9821_2 
## Running HL: 9821_3 
## Running HL: 9822_3 
## Running HL: 9824_1 
## Running HL: 9824_2 
## Running HL: 9824_3 
## Running HL: 9825_1 
## Running HL: 9825_2 
## Running HL: 9825_3 
## Running HL: 9826_1 
## Running HL: 9826_2 
## Running HL: 9826_3 
## Running HL: 9829_1 
## Running HL: 9829_2 
## Running HL: 9829_3 
## Running HL: 9833_1 
## Running HL: 9833_2 
## Running HL: 9834_1 
## Running HL: 9834_2 
## Running HL: 9834_3 
## Running HL: 9835_1 
## Running HL: 9835_2 
## Running HL: 9835_3 
## Running HL: 9836_1 
## Running HL: 9836_2 
## Running HL: 9836_3 
## Running HL: 9838_1 
## Running HL: 9838_2 
## Running HL: 9838_3 
## Running HL: 9839_1 
## Running HL: 9839_2 
## Running HL: 9839_3 
## Running HL: 9840_1 
## Running HL: 9840_2 
## Running HL: 9840_3 
## Running HL: 9841_1 
## Running HL: 9841_2 
## Running HL: 9841_3 
## Running HL: 9843_1 
## Running HL: 9843_2 
## Running HL: 9843_3 
## Running HL: 9844_1 
## Running HL: 9844_2 
## Running HL: 9844_3 
## Running HL: 9845_1 
## Running HL: 9845_2 
## Running HL: 9845_3 
## Running HL: 9846_1 
## Running HL: 9846_2 
## Running HL: 9846_3 
## Running HL: 9848_1 
## Running HL: 9848_2 
## Running HL: 9848_3 
## Running HL: 9849_1 
## Running HL: 9849_2 
## Running HL: 9849_3 
## Running HL: 9850_1 
## Running HL: 9850_2 
## Running HL: 9850_3 
## Running HL: 9851_1 
## Running HL: 9851_2 
## Running HL: 9851_3 
## Running HL: 9852_1 
## Running HL: 9852_2 
## Running HL: 9852_3 
## Running HL: 9855_1 
## Running HL: 9855_2 
## Running HL: 9855_3 
## Running HL: 9856_1 
## Running HL: 9856_2 
## Running HL: 9856_3 
## Running HL: 9857_1 
## Running HL: 9857_2 
## Running HL: 9857_3 
## Running HL: 9858_1 
## Running HL: 9858_2 
## Running HL: 9858_3 
## Running HL: 9859_3 
## Running HL: 9860_1 
## Running HL: 9860_2 
## Running HL: 9860_3 
## Running HL: 9861_1 
## Running HL: 9861_2 
## Running HL: 9861_3 
## Running HL: 9864_1 
## Running HL: 9864_2 
## Running HL: 9864_3 
## Running HL: 9867_1 
## Running HL: 9867_2 
## Running HL: 9867_3 
## Running HL: 9868_1 
## Running HL: 9868_2 
## Running HL: 9868_3 
## Running HL: 9870_2 
## Running HL: 9870_3 
## Running HL: 9874_1 
## Running HL: 9874_2 
## Running HL: 9874_3 
## Running HL: 9876_1 
## Running HL: 9876_3 
## Running HL: 9877_1 
## Running HL: 9877_2 
## Running HL: 9877_3 
## Running HL: 9878_1 
## Running HL: 9878_2 
## Running HL: 9878_3 
## Running HL: 9881_1 
## Running HL: 9881_2 
## Running HL: 9881_3 
## Running HL: 9883_1 
## Running HL: 9883_2 
## Running HL: 9883_3 
## Running HL: 9885_1 
## Running HL: 9885_2 
## Running HL: 9885_3 
## Running HL: 9886_2 
## Running HL: 9886_3 
## Running HL: 9888_1 
## Running HL: 9888_2 
## Running HL: 9888_3 
## Running HL: 9890_1 
## Running HL: 9890_2 
## Running HL: 9890_3 
## Running HL: 9891_1 
## Running HL: 9891_2 
## Running HL: 9891_3 
## Running HL: 9892_1 
## Running HL: 9895_1 
## Running HL: 9895_2 
## Running HL: 9895_3 
## Running HL: 9897_1 
## Running HL: 9897_2 
## Running HL: 9897_3 
## Running HL: 9898_1 
## Running HL: 9898_2 
## Running HL: 9899_1 
## Running HL: 9899_2 
## Running HL: 9899_3 
## Running HL: 9901_1 
## Running HL: 9901_2 
## Running HL: 9901_3 
## Running HL: 9902_1 
## Running HL: 9902_2 
## Running HL: 9902_3 
## Running HL: 9903_1 
## Running HL: 9903_2 
## Running HL: 9903_3 
## Running HL: 9904_1 
## Running HL: 9904_2 
## Running HL: 9904_3 
## Running HL: 9906_1 
## Running HL: 9906_2 
## Running HL: 9906_3 
## Running HL: 991_1 
## Running HL: 991_2 
## Running HL: 991_3 
## Running HL: 992_1 
## Running HL: 992_2 
## Running HL: 992_3 
## Running HL: 9927_1 
## Running HL: 9927_2 
## Running HL: 9927_3 
## Running HL: 9931_1 
## Running HL: 9931_2 
## Running HL: 9931_3 
## Running HL: 9938_2 
## Running HL: 9938_3 
## Running HL: 9942_1 
## Running HL: 9942_2 
## Running HL: 9942_3 
## Running HL: 9948_3 
## Running HL: 9949_1 
## Running HL: 9949_2 
## Running HL: 9949_3 
## Running HL: 9950_1 
## Running HL: 9950_2 
## Running HL: 9950_3 
## Running HL: 997_1 
## Running HL: 997_2 
## Running HL: 997_3 
## Running HL: Ayx_3 
## Running HL: Cbb_1 
## Running HL: Cbb_2 
## Running HL: Cbb_3 
## Running HL: Ctl-62_1 
## Running HL: Ctl-62_2 
## Running HL: Ctl-62_3 
## Running HL: Gwx-11_1 
## Running HL: Gwx-11_2 
## Running HL: Gwx-11_3 
## Running HL: Hha-21_1 
## Running HL: Hha-21_2 
## Running HL: Hha-21_3 
## Running HL: Hnxy-46_1 
## Running HL: Hnxy-46_2 
## Running HL: Hnxy-46_3 
## Running HL: HNzjj-75_1 
## Running HL: HNzjj-75_2 
## Running HL: HNzjj-75_3 
## Running HL: Hwc-61_2 
## Running HL: Hwc-61_3 
## Running HL: Hyl-40_1 
## Running HL: Hyl-40_2 
## Running HL: Hyl-40_3 
## Running HL: Jjgs-30_1 
## Running HL: Jjgs-30_2 
## Running HL: Jjgs-30_3 
## Running HL: Jjj-32_3 
## Running HL: Jnf-27_1 
## Running HL: Jnf-27_2 
## Running HL: Jnf-27_3 
## Running HL: Scg-77_1 
## Running HL: Scg-77_2 
## Running HL: Scg-77_3 
## Running HL: Xem_1 
## Running HL: Xem_2 
## Running HL: Xem_3 
## Running HL: Xyl_1 
## Running HL: Xyl_2 
## Running HL: Xyl_3 
## Running HL: Ygs-39_1 
## Running HL: Ygs-39_2 
## Running HL: Ygs-39_3 
## Running HL: Zla-62_2 
## Running HL: Zla-62_3
```

```
########################extract 3 coefficients of the fit
mat <- data.frame(matrix(ncol=7,nrow=0))
for (i in plants) {
  
  coeff.HL <- grHL[[i]]$coefficients
  sl.HL <- coeff.HL[grep("Slope",names(coeff.HL))]
  upp.HL <- coeff.HL[grep("FS",names(coeff.HL))]
  t50.HL <-  coeff.HL[grep("t50",names(coeff.HL))]
  mat <- rbind(mat,
               data.frame(line=rep(i,length(sl.HL))
                          ,Region=rep(HLdiam.melted[which(HLdiam.melted$Plant==i)[1],"Region"],length(sl.HL))
                          ,treatment="HL",Slope=sl.HL,Upper=upp.HL
                          ,t50 =t50.HL)
  )
}
```

#### overview and remove outliers

```
head(mat)
```

```
##                      line          Region treatment     Slope    Upper      t50
## Slope:(Intercept)  1002_1 Northern_Europe        HL -3.929536 7.381513 20.16156
## Slope:(Intercept)1 1002_2 Northern_Europe        HL -4.588898 9.041317 17.27498
## Slope:(Intercept)2 1002_3 Northern_Europe        HL -5.007694 8.119209 19.53363
## Slope:(Intercept)3 1061_1 Northern_Europe        HL -4.704080 8.737315 17.96007
## Slope:(Intercept)4 1061_2 Northern_Europe        HL -5.756125 8.249669 17.50436
## Slope:(Intercept)5 1061_3 Northern_Europe        HL -4.797550 6.836094 28.56952
```

```
plot(mat$Slope)
```

```
plot(mat$Upper)
```

```
plot(mat$t50)
```

```
#mat<-mat[mat$line!="539_1",] #if necessary remove outlier lines with values that are very far off
head(mat)
```

```
##                      line          Region treatment     Slope    Upper      t50
## Slope:(Intercept)  1002_1 Northern_Europe        HL -3.929536 7.381513 20.16156
## Slope:(Intercept)1 1002_2 Northern_Europe        HL -4.588898 9.041317 17.27498
## Slope:(Intercept)2 1002_3 Northern_Europe        HL -5.007694 8.119209 19.53363
## Slope:(Intercept)3 1061_1 Northern_Europe        HL -4.704080 8.737315 17.96007
## Slope:(Intercept)4 1061_2 Northern_Europe        HL -5.756125 8.249669 17.50436
## Slope:(Intercept)5 1061_3 Northern_Europe        HL -4.797550 6.836094 28.56952
```

```
#write.table(mat, file="HLparametersperline.txt")#save table
```

##### Same for LL

```
######Plot Figure 1
#create drm models per region from which I can predict for plotting
WEHL <- HLdiam.melted[HLdiam.melted$Region=="Western_Europe",]
WEHL <- WEHL[which(WEHL$value!= "NA"),] 
drm.WEHL <- drm(value~time, fct=LL.4(fixed=c(NA,0,NA,NA), names=c("Slope","0", "FinalSize", "t50")), data = WEHL, na.action = na.omit)

NEHL <- HLdiam.melted[HLdiam.melted$Region=="Northern_Europe",]
NEHL <- NEHL[which(NEHL$value!= "NA"),] 
drm.NEHL <- drm(value~time, fct=LL.4(fixed=c(NA,0,NA,NA), names=c("Slope","0", "FinalSize", "t50")), data = NEHL, na.action = na.omit)

CHHL <- HLdiam.melted[HLdiam.melted$Region=="China",]
CHHL <- CHHL[which(CHHL$value!= "NA"),] 
drm.CHHL <- drm(value~time, fct=LL.4(fixed=c(NA,0,NA,NA), names=c("Slope","0", "FinalSize", "t50")), data = CHHL, na.action = na.omit)

SPHL <- HLdiam.melted[HLdiam.melted$Region=="Spain",]
SPHL <- SPHL[which(SPHL$value!= "NA"),] 
drm.SPHL <- drm(value~time, fct=LL.4(fixed=c(NA,0,NA,NA), names=c("Slope","0", "FinalSize", "t50")), data = SPHL, na.action = na.omit)
############
#LL
############
WELL <- LLdiam.melted[LLdiam.melted$Region=="Western_Europe",]
WELL <- WELL[which(WELL$value!= "NA"),] 
drm.WELL <- drm(value~time, fct=LL.4(fixed=c(NA,0,NA,NA), names=c("Slope","0", "FinalSize", "t50")), data = WELL, na.action = na.omit)

NELL <- LLdiam.melted[LLdiam.melted$Region=="Northern_Europe",]
NELL <- NELL[which(NELL$value!= "NA"),] 
drm.NELL <- drm(value~time, fct=LL.4(fixed=c(NA,0,NA,NA), names=c("Slope","0", "FinalSize", "t50")), data = NELL, na.action = na.omit)

CHLL <- LLdiam.melted[LLdiam.melted$Region=="China",]
CHLL <- CHLL[which(CHLL$value!= "NA"),] 
drm.CHLL <- drm(value~time, fct=LL.4(fixed=c(NA,0,NA,NA), names=c("Slope","0", "FinalSize", "t50")), data = CHLL, na.action = na.omit)

SPLL <- LLdiam.melted[LLdiam.melted$Region=="Spain",]
SPLL <- SPLL[which(SPLL$value!= "NA"),] 
drm.SPLL <- drm(value~time, fct=LL.4(fixed=c(NA,0,NA,NA), names=c("Slope","0", "FinalSize", "t50")), data = SPLL, na.action = na.omit)


#both
#png("Fig1.png", width = 4000,height = 3000,res=500)
par(lwd=2.5,cex=1)
plot(x=0,y=0, main ="Average regional growth rate",type="n", col="dark green",lwd=1, lty = 1
     , ylim =c(0,16),
     xlim = c(0,100), log="", ylab = "diameter [cm]", xlab ="t [days]", cex.axis=1.5, cex.lab=1.5)
#params[[i]] <- c(slope=drm.mod$fit$par[1],upper=drm.mod$fit$par[2],t50=drm.mod$fit$par[3])
xx<-seq(0,100, length=100) # coordinates of your x axis
#HL
lines(xx, predict(drm.WEHL,data.frame(x=xx)),lwd=2, lty=2,col="red") # plot the predicted fitted model coming from the logistic equation
lines(xx, predict(drm.NEHL,data.frame(x=xx)),lwd=2, lty=2,col="green4")
lines(xx, predict(drm.CHHL,data.frame(x=xx)),lwd=2, lty=2,col="orange")
lines(xx, predict(drm.SPHL,data.frame(x=xx)),lwd=2, lty=2,col="purple")
#LL
lines(xx, predict(drm.WELL,data.frame(x=xx)),lwd=2, col="red") # plot the predicted fitted model coming from the logistic equation
lines(xx, predict(drm.NELL,data.frame(x=xx)),lwd=2, col="green4")
lines(xx, predict(drm.CHLL,data.frame(x=xx)),lwd=2, col="orange")
lines(xx, predict(drm.SPLL,data.frame(x=xx)),lwd=2, col="purple")
legend(60,7, c("China", "Northern Europe", "Spain","Western Europe","LL","HL"), 
       lty=c(1,1,1,1,1,2),bty="n", 
       lwd=c(2.5,2.5),col=c("orange", "green4", "purple","red" ,"black","black"), cex=1.2)
```

```
#dev.off()
```

```
#Plotting

###to have different line colors per region:
ids_in$color <- NA
ids_in[which(ids_in$Region=="China"),"color"] <- "orange" 
ids_in[which(ids_in$Region=="Western_Europe"),"color"] <- "red" 
ids_in[which(ids_in$Region=="Northern_Europe"),"color"] <- "green4" 
ids_in[which(ids_in$Region=="Spain"),"color"] <- "purple" 

#plot them all together:
par(lwd=2.5,cex=1)
plot(x=0,y=0, main ="Growth rate of 20 genotypes",type="n", col="dark green",lwd=1, lty = 1
     , ylim =c(0,25),
     xlim = c(0,100), log="", ylab = "diameter [cm]", xlab ="t [days]", cex.axis=1.5, cex.lab=1.5)
#params[[i]] <- c(slope=drm.mod$fit$par[1],upper=drm.mod$fit$par[2],t50=drm.mod$fit$par[3])
xx<-seq(0,100, length=100) # coordinates of your x axis

for(i in ids_chosen){
  #HL
  samp <- HLdiam.melted[HLdiam.melted$ID==i,]
  drm.mod <- drm(value~time, fct=LL.4(fixed=c(NA,0,NA,NA), names=c("Slope","0", "FS", "t50")), data = samp, na.action = na.omit)
  lines(xx, predict(drm.mod,data.frame(x=xx)), col=ids_in[which(ids_in$ID == i),"color"],lty=20) # plot the predicted fitted model coming from the logistic equation
  #LL
  samp <- LLdiam.melted[LLdiam.melted$ID==i,]
  drm.mod <- drm(value~time, fct=LL.4(fixed=c(NA,0,NA,NA), names=c("Slope","0", "FS", "t50")), data = samp, na.action = na.omit)
  lines(xx, predict(drm.mod,data.frame(x=xx)), col=ids_in[which(ids_in$ID == i),"color"],lty=1) # plot the predicted fitted model coming from the logistic equation
  
  }
```

```
#plot them in separate plots:
xx<-seq(0,100, length=100) # coordinates of x axis

#svg("~/Desktop/Genotype_growthplot.svg",width=20,height=15)
#layout(matrix(1:20,nrow=4,ncol=5,byrow = T))#these commands are for plotting it into a file
for(i in ids_chosen){
  par(lwd=2.5,cex=1)
  plot(x=0,y=0, main =paste(unique(HLdiam.melted[which(HLdiam.melted$ID==i),"Region"]),i,sep=": "),type="n", col="dark green",lwd=1, lty = 1
     , ylim =c(0,25), xlim = c(0,100), log="", ylab = "diameter [cm]", xlab ="t [days]", cex.axis=0.8, cex.lab=0.8)#small axes to make them fit in a single figure
  #HL
  samp <- HLdiam.melted[HLdiam.melted$ID==i,]
  drm.mod <- drm(value~time, fct=LL.4(fixed=c(NA,0,NA,NA), names=c("Slope","0", "FS", "t50")), data = samp, na.action = na.omit)
  lines(xx, predict(drm.mod,data.frame(x=xx)), col=ids_in[which(ids_in$ID == i),"color"],lty=2) 
  #add data_points for the measurements:
  points(HLdiam.melted[which(HLdiam.melted$ID==i),"time"],HLdiam.melted[which(HLdiam.melted$ID==i),"value"],col="red",cex=0.5)
    #LL
  samp <- LLdiam.melted[LLdiam.melted$ID==i,]
  drm.mod <- drm(value~time, fct=LL.4(fixed=c(NA,0,NA,NA), names=c("Slope","0", "FS", "t50")), data = samp, na.action = na.omit)
  lines(xx, predict(drm.mod,data.frame(x=xx)), col=ids_in[which(ids_in$ID == i),"color"],lty=1)
    #add data_points for the measurements:
  points(LLdiam.melted[which(LLdiam.melted$ID==i),"time"],LLdiam.melted[which(LLdiam.melted$ID==i),"value"],col="blue",cex=0.5)
 
 legend(0,26, c("HL","LL","HL","LL"), 
       lty=c(2,1,NA,NA),bty="n", pch=c(NA,NA,1,1),
       lwd=c(2.5,2.5),col=c("black","black","red","blue"), cex=0.8)
  }
```

```
#dev.off()
```

### PART2

###### Genotypic mean estimation

##### Estimate genotype means using glms

```
#FS
mean_FSHL <- glm(Final.Size ~ ID + Replicate+tray/(row+column)-1, data=hl)
plot(mean_FSHL)
```

```
## Warning: not plotting observations with leverage one:
##   18, 264, 494, 535, 553, 573, 734, 761
```

```
## Warning: not plotting observations with leverage one:
##   18, 264, 494, 535, 553, 573, 734, 761
```

```
#summary(mean_FSHL) 
#t50
hist(hl$t50)#~normal with a tail of high t50s
```

```
mean_t50HL <- glm(t50 ~ ID + Replicate+tray/(row+column)-1, data=hl)
plot(mean_t50HL)
```

```
## Warning: not plotting observations with leverage one:
##   18, 264, 494, 535, 553, 573, 734, 761
```

```
## Warning: not plotting observations with leverage one:
##   18, 264, 494, 535, 553, 573, 734, 761
```

```
#summary(mean_t50HL)
#Slope
hist(hl$Slope)#~normal with a tail of high slopes
```

```
mean_slopeHL <- glm(Slope ~ ID + Replicate+tray/(row+column)-1, data=hl)
plot(mean_slopeHL)
```

```
## Warning: not plotting observations with leverage one:
##   18, 264, 494, 535, 553, 573, 734, 761
```

```
## Warning: not plotting observations with leverage one:
##   18, 264, 494, 535, 553, 573, 734, 761
```

```
#summary(mean_slopeHL)

###FT - a lot of NAs
length(hl$flowering)
```

```
## [1] 778
```

```
length(na.omit(hl$flowering))#296 of 778 plants flowered --> I will just average the phenotypes
```

```
## [1] 296
```

```
library(plyr)
#HL
HLFL <- ddply(hl, c("ID"), summarise,
                    FL = mean(flowering, na.rm=T))
LLFL <- ddply(ll, c("ID"), summarise,
              FL = mean(flowering, na.rm=T))
```

### PART3

### Manova, regional differentiation, GXE

```
###MANOVA
corrected_data_both <- read.delim(file="S3.txt")
head(corrected_data_both)
```

```
##               Name   ID          Region Country Final_Size_HL  T50_HL Slope_HL
## 1  Ale-Stenar-41-1  991 Northern Europe  Sweden        8.6731 10.5321   5.7169
## 2  Ale-Stenar-44-4  992 Northern Europe  Sweden        7.2023 11.8398   4.8478
## 3 Ale-Stenar-56-14  997 Northern Europe  Sweden        6.6369 15.5743   4.0614
## 4 Ale-Stenar-64-24 1002 Northern Europe  Sweden        7.6601 13.7759   3.7600
## 5   Brösarp-11-135 1061 Northern Europe  Sweden        7.1276 15.5622   4.9051
## 6   Brösarp-15-138 1062 Northern Europe  Sweden        5.8695 16.4978   3.7495
##   Final_Size_LL  T50_LL Slope_LL GXE_Final_Size GXE_t50 GXE_Slope
## 1       14.5374 34.5172   5.1963        -0.7409  0.4742   -1.6368
## 2       13.7532 33.5521   5.5908        -0.0543 -1.7986   -0.3731
## 3       15.2798 32.4278   5.6417         2.0377 -6.6575    0.4641
## 4       14.2653 37.2868   4.8761        -1.1611 -0.5669    0.9047
## 5       11.4213 35.4196   5.7678        -2.3115 -3.6535   -0.2535
## 6       11.0332 31.6869   5.6354        -1.4415 -8.3219    0.7698
##   Phenotype_analysis GWAS DiamFieldM2 Hypocotyllength Biomass21d  FT_10  FT_16
## 1                  1    1          NA              NA         NA 110.50 106.75
## 2                  0    1          NA              NA         NA 116.00 100.00
## 3                  0    1   14.375000               5     0.0343 101.25  87.50
## 4                  0    1    8.069833              10     0.0239 102.75  86.25
## 5                  1    1    9.499000               7     0.0218  97.75  94.75
## 6                  0    1   10.855333               4     0.0301  94.00  68.25
##   FT_HL FT_LL
## 1    NA    NA
## 2    NA    NA
## 3    NA  81.0
## 4    NA    NA
## 5    NA    NA
## 6    NA  88.5
```

```
###modify to have phenotypes in one column and treatment as info, only keep the non-duplicate location lines
hl <- corrected_data_both[which(corrected_data_both$Phenotype_analysis==1),c("Name","ID","Region","Country","Final_Size_HL","T50_HL","Slope_HL")]
ll <- corrected_data_both[which(corrected_data_both$Phenotype_analysis==1),c("Name","ID","Region","Country","Final_Size_LL","T50_LL","Slope_LL")]

colnames(hl) <- c("Name","ID","Region","Country","FS","T50","SL")
colnames(ll) <- c("Name","ID","Region","Country","FS","T50","SL")

hl$treatment <- "HL"
ll$treatment <- "LL"

corrected_data_both <- rbind(hl,ll)
corrected_data_both$treatment <- as.factor(corrected_data_both$treatment)
head(corrected_data_both)
```

```
##               Name   ID          Region Country     FS     T50     SL treatment
## 1  Ale-Stenar-41-1  991 Northern Europe  Sweden 8.6731 10.5321 5.7169        HL
## 5   Brösarp-11-135 1061 Northern Europe  Sweden 7.1276 15.5622 4.9051        HL
## 9          App1-12 5830 Northern Europe  Sweden 6.0112 12.8023 4.0086        HL
## 15          Fly2-1 6023 Northern Europe  Sweden 6.2978 13.4606 4.7825        HL
## 17         Hov1-10 6035 Northern Europe  Sweden 6.2719 17.7050 5.8510        HL
## 21           Kni-1 6040 Northern Europe  Sweden 6.8210 12.3300 3.9022        HL
```

```
growth <- as.matrix(corrected_data_both[,c("FS","T50","SL")])
fit<-manova(growth~treatment*Region, data=corrected_data_both)
summary(fit)
```

```
##                   Df  Pillai approx F num Df den Df    Pr(>F)    
## treatment          1 0.94886  2665.54      3    431 < 2.2e-16 ***
## Region             4 0.19817     7.66     12   1299 7.334e-14 ***
## treatment:Region   4 0.05169     1.90     12   1299   0.03073 *  
## Residuals        433                                             
## ---
## Signif. codes:  0 '***' 0.001 '**' 0.01 '*' 0.05 '.' 0.1 ' ' 1
```

```
summary.aov(fit)#gives the individual anovas
```

```
##  Response FS :
##                   Df Sum Sq Mean Sq  F value    Pr(>F)    
## treatment          1 3962.7  3962.7 2284.841 < 2.2e-16 ***
## Region             4  121.9    30.5   17.574 2.235e-13 ***
## treatment:Region   4    9.6     2.4    1.383    0.2389    
## Residuals        433  751.0     1.7                       
## ---
## Signif. codes:  0 '***' 0.001 '**' 0.01 '*' 0.05 '.' 0.1 ' ' 1
## 
##  Response T50 :
##                   Df Sum Sq Mean Sq   F value    Pr(>F)    
## treatment          1  47230   47230 6008.1930 < 2.2e-16 ***
## Region             4    231      58    7.3540  9.76e-06 ***
## treatment:Region   4     68      17    2.1697   0.07154 .  
## Residuals        433   3404       8                        
## ---
## Signif. codes:  0 '***' 0.001 '**' 0.01 '*' 0.05 '.' 0.1 ' ' 1
## 
##  Response SL :
##                   Df Sum Sq Mean Sq F value    Pr(>F)    
## treatment          1  51.14  51.137 55.7659 4.536e-13 ***
## Region             4   3.70   0.926  1.0094   0.40219    
## treatment:Region   4  11.79   2.949  3.2155   0.01282 *  
## Residuals        433 397.06   0.917                      
## ---
## Signif. codes:  0 '***' 0.001 '**' 0.01 '*' 0.05 '.' 0.1 ' ' 1
## 
## 9 observations deleted due to missingness
```

### GXE Estimation

```
#####GXE

###I will estimate GxE for all individuals:
#prepare the input data:
data_both <- read.delim(file="S3.txt")
head(data_both)
```

```
##               Name   ID          Region Country Final_Size_HL  T50_HL Slope_HL
## 1  Ale-Stenar-41-1  991 Northern Europe  Sweden        8.6731 10.5321   5.7169
## 2  Ale-Stenar-44-4  992 Northern Europe  Sweden        7.2023 11.8398   4.8478
## 3 Ale-Stenar-56-14  997 Northern Europe  Sweden        6.6369 15.5743   4.0614
## 4 Ale-Stenar-64-24 1002 Northern Europe  Sweden        7.6601 13.7759   3.7600
## 5   Brösarp-11-135 1061 Northern Europe  Sweden        7.1276 15.5622   4.9051
## 6   Brösarp-15-138 1062 Northern Europe  Sweden        5.8695 16.4978   3.7495
##   Final_Size_LL  T50_LL Slope_LL GXE_Final_Size GXE_t50 GXE_Slope
## 1       14.5374 34.5172   5.1963        -0.7409  0.4742   -1.6368
## 2       13.7532 33.5521   5.5908        -0.0543 -1.7986   -0.3731
## 3       15.2798 32.4278   5.6417         2.0377 -6.6575    0.4641
## 4       14.2653 37.2868   4.8761        -1.1611 -0.5669    0.9047
## 5       11.4213 35.4196   5.7678        -2.3115 -3.6535   -0.2535
## 6       11.0332 31.6869   5.6354        -1.4415 -8.3219    0.7698
##   Phenotype_analysis GWAS DiamFieldM2 Hypocotyllength Biomass21d  FT_10  FT_16
## 1                  1    1          NA              NA         NA 110.50 106.75
## 2                  0    1          NA              NA         NA 116.00 100.00
## 3                  0    1   14.375000               5     0.0343 101.25  87.50
## 4                  0    1    8.069833              10     0.0239 102.75  86.25
## 5                  1    1    9.499000               7     0.0218  97.75  94.75
## 6                  0    1   10.855333               4     0.0301  94.00  68.25
##   FT_HL FT_LL
## 1    NA    NA
## 2    NA    NA
## 3    NA  81.0
## 4    NA    NA
## 5    NA    NA
## 6    NA  88.5
```

```
###modify to have phenotypes in one column and treatment as info
hl <- data_both[,c("Name","ID","Region","Country","Final_Size_HL","T50_HL","Slope_HL")]
ll <- data_both[,c("Name","ID","Region","Country","Final_Size_LL","T50_LL","Slope_LL")]

colnames(hl) <- c("Name","ID","Region","Country","FS","T50","SL")
colnames(ll) <- c("Name","ID","Region","Country","FS","T50","SL")

hl$treatment <- "HL"
ll$treatment <- "LL"

geno_means <- rbind(hl,ll)
geno_means$treatment <- as.factor(geno_means$treatment)
head(geno_means)
```

```
##               Name   ID          Region Country     FS     T50     SL treatment
## 1  Ale-Stenar-41-1  991 Northern Europe  Sweden 8.6731 10.5321 5.7169        HL
## 2  Ale-Stenar-44-4  992 Northern Europe  Sweden 7.2023 11.8398 4.8478        HL
## 3 Ale-Stenar-56-14  997 Northern Europe  Sweden 6.6369 15.5743 4.0614        HL
## 4 Ale-Stenar-64-24 1002 Northern Europe  Sweden 7.6601 13.7759 3.7600        HL
## 5   Brösarp-11-135 1061 Northern Europe  Sweden 7.1276 15.5622 4.9051        HL
## 6   Brösarp-15-138 1062 Northern Europe  Sweden 5.8695 16.4978 3.7495        HL
```

```
GxE_FS <- glm(FS~treatment*ID-1,data=geno_means)
#summary(GxE_FS)

#####extract coefficients (I need the interaction term for GxE)
gxeFS_estimate <- GxE_FS$coefficients
head(gxeFS_estimate)
```

```
## treatmentHL treatmentLL      ID1061      ID1062      ID1063      ID1066 
##      7.6601     14.2653     -0.5325     -1.7906     -0.4141      0.2766
```

```
gxeFS_estimate_1 <- gxeFS_estimate[1:281]#individual
tail(gxeFS_estimate_1)
```

```
## IDSmx-34    IDXem    IDXyl IDYgs-39 IDZkh-29 IDZla-62 
##  -3.7081  -0.2194  -0.8206  -0.8156  -1.4582   1.3225
```

```
gxeFS_estimate_2 <- data.frame(value=gxeFS_estimate[282:560])
head(gxeFS_estimate_2)
```

```
##                          value
## treatmentLL:ID1061     -2.3115
## treatmentLL:ID1062     -1.4415
## treatmentLL:ID1063     -0.6944
## treatmentLL:ID1066      0.3013
## treatmentLL:ID2014-241      NA
## treatmentLL:ID2014-242  1.2209
```

```
tail(gxeFS_estimate_2)
```

```
##                          value
## treatmentLL:IDSmx-34        NA
## treatmentLL:IDXem    -2.545818
## treatmentLL:IDXyl           NA
## treatmentLL:IDYgs-39 -1.358935
## treatmentLL:IDZkh-29        NA
## treatmentLL:IDZla-62 -0.900400
```

```
#create data frame
ID <- sapply(strsplit(rownames(gxeFS_estimate_2), "ID"), paste)[2,]
gxeFS <- data.frame(cbind(ID,GxE=gxeFS_estimate_2$value))
head(gxeFS)
```

```
##         ID                GxE
## 1     1061  -2.31150000000003
## 2     1062  -1.44150000000001
## 3     1063 -0.694400000000025
## 4     1066  0.301299999999994
## 5 2014-241               <NA>
## 6 2014-242             1.2209
```

```
str(gxeFS)
```

```
## 'data.frame':    279 obs. of  2 variables:
##  $ ID : Factor w/ 279 levels "1061","1062",..: 1 2 3 4 5 6 7 8 9 10 ...
##  $ GxE: Factor w/ 270 levels "-0.0381999999999829",..: 143 103 55 194 NA 245 106 113 52 108 ...
```

```
gxeFS$GxE <- as.numeric(as.character(gxeFS$GxE))
###this can be added to table S3

#same script for t50 and SL:
GxE_t50 <- glm(T50~treatment*ID-1,data=geno_means)
#summary(GxE_t50)

#####extract coefficients (I need the interaction term for GxE)
gxet50_estimate <- GxE_t50$coefficients
head(gxet50_estimate)
```

```
## treatmentHL treatmentLL      ID1061      ID1062      ID1063      ID1066 
##     13.7759     37.2868      1.7863      2.7219      0.2218     -2.6780
```

```
gxet50_estimate_2 <- data.frame(value=gxet50_estimate[282:560])
head(gxet50_estimate_2)
```

```
##                          value
## treatmentLL:ID1061     -3.6535
## treatmentLL:ID1062     -8.3218
## treatmentLL:ID1063     -7.0445
## treatmentLL:ID1066      0.1089
## treatmentLL:ID2014-241      NA
## treatmentLL:ID2014-242 -1.2582
```

```
tail(gxet50_estimate_2)
```

```
##                           value
## treatmentLL:IDSmx-34         NA
## treatmentLL:IDXem    -13.387999
## treatmentLL:IDXyl            NA
## treatmentLL:IDYgs-39  -9.333376
## treatmentLL:IDZkh-29         NA
## treatmentLL:IDZla-62  -3.235700
```

```
#create data frame
ID <- sapply(strsplit(rownames(gxet50_estimate_2), "ID"), paste)[2,]
gxet50 <- data.frame(cbind(ID,GxE=gxet50_estimate_2$value))
gxet50$GxE <- as.numeric(as.character(gxet50$GxE))
head(gxet50)
```

```
##         ID     GxE
## 1     1061 -3.6535
## 2     1062 -8.3218
## 3     1063 -7.0445
## 4     1066  0.1089
## 5 2014-241      NA
## 6 2014-242 -1.2582
```

```
str(gxet50)
```

```
## 'data.frame':    279 obs. of  2 variables:
##  $ ID : Factor w/ 279 levels "1061","1062",..: 1 2 3 4 5 6 7 8 9 10 ...
##  $ GxE: num  -3.654 -8.322 -7.045 0.109 NA ...
```

```
#Slope:
GxE_SL <- glm(SL~treatment*ID-1,data=geno_means)
#summary(GxE_SL)

#####extract coefficients (I need the interaction term for GxE)
gxeSL_estimate <- GxE_SL$coefficients
head(gxeSL_estimate)
```

```
## treatmentHL treatmentLL      ID1061      ID1062      ID1063      ID1066 
##      3.7600      4.8761      1.1451     -0.0105      3.2135      1.0983
```

```
gxeSL_estimate_2 <- data.frame(value=gxeSL_estimate[282:560])
head(gxeSL_estimate_2)
```

```
##                          value
## treatmentLL:ID1061     -0.2534
## treatmentLL:ID1062      0.7698
## treatmentLL:ID1063     -2.5598
## treatmentLL:ID1066     -1.0479
## treatmentLL:ID2014-241      NA
## treatmentLL:ID2014-242 -1.0144
```

```
tail(gxeSL_estimate_2)
```

```
##                           value
## treatmentLL:IDSmx-34         NA
## treatmentLL:IDXem    -0.2695575
## treatmentLL:IDXyl            NA
## treatmentLL:IDYgs-39  0.3988388
## treatmentLL:IDZkh-29         NA
## treatmentLL:IDZla-62  1.3563000
```

```
#create data frame
ID <- sapply(strsplit(rownames(gxeSL_estimate_2), "ID"), paste)[2,]
gxeSL <- data.frame(cbind(ID,GxE=gxeSL_estimate_2$value))
gxeSL$GxE <- as.numeric(as.character(gxeSL$GxE))
head(gxeSL)
```

```
##         ID     GxE
## 1     1061 -0.2534
## 2     1062  0.7698
## 3     1063 -2.5598
## 4     1066 -1.0479
## 5 2014-241      NA
## 6 2014-242 -1.0144
```

```
str(gxeSL)
```

```
## 'data.frame':    279 obs. of  2 variables:
##  $ ID : Factor w/ 279 levels "1061","1062",..: 1 2 3 4 5 6 7 8 9 10 ...
##  $ GxE: num  -0.253 0.77 -2.56 -1.048 NA ...
```

##### Plot GxE (Fig S2)

```
S3 <- read.delim(file="S3.txt")

x <- S3$Final_Size_HL
y <- S3$Final_Size_LL
z <- S3$GXE_Final_Size

dat <- data.frame(x=x,y=y,z=z)
rbPal <- colorRampPalette(c('blue','red'))
dat$Col <- rbPal(5)[as.numeric(cut(dat$z,breaks = 5))]#10 might be to narrow, just take 5

brks <- round(quantile(z,probs = seq(0,1,0.1),na.rm = T),1)
brks
```

```
##   0%  10%  20%  30%  40%  50%  60%  70%  80%  90% 100% 
## -8.0 -2.3 -1.8 -1.2 -0.9 -0.5 -0.1  0.2  0.6  1.2  3.6
```

```
#plot
plot(dat$x,dat$y,pch = 20,col = dat$Col, cex=2, xlab = "Final Size - HL [cm]", ylab="Final Size - LL [cm]", main="GxE for Final Size", cex.lab=1.5, cex.axis=1.5,cex.main=2)
abline(lm(dat$y~dat$x),lwd=4)
#add legend
legend_image <- as.raster(matrix(rbPal(5), ncol=1))
text(x=6, y=c(15.2,17,18.8),labels = brks[c(1,6,11)])
rasterImage(legend_image, 5.8,19, 5.5,15)
text(5.3,17,"GxE",srt=90, cex=1.5)
##add mean t50 in HL and LL as big black dot
points(mean(x,na.rm = T),mean(y,na.rm=T),pch=20,cex=5)
#error bars
xm <- mean(x,na.rm = T)
ym <- mean(y,na.rm=T)
xsd <- sd(x,na.rm=T)
ysd <- sd(y,na.rm=T)
#segments(mean(x,na.rm = T),mean(y,na.rm=T)-sd(y,na.rm = T),mean(x,na.rm = T),mean(y,na.rm=T)+sd(y,na.rm = T))
epsilon <- 0.4
segments(xm,ym-ysd , xm, ym+ysd, lwd = 4)#error bars
segments(xm-epsilon, ym+ysd , xm+epsilon, ym+ysd,lwd=3)#vertical lines
segments(xm-epsilon, ym-ysd , xm+epsilon, ym-ysd,lwd=3)
#for y:
segments(xm-xsd,ym , xm+xsd, ym,lwd=4)#error bars
segments(xm+xsd,ym-epsilon, xm+xsd,ym+epsilon,lwd=3)#vertical lines
segments(xm-xsd,ym-epsilon, xm-xsd,ym+epsilon,lwd=3)
```

### Regional differentiation

```
#Regional differentiation
library(multcomp)
```

```
## Loading required package: mvtnorm
```

```
## Loading required package: survival
```

```
## Loading required package: TH.data
```

```
## 
## Attaching package: 'TH.data'
```

```
## The following object is masked from 'package:MASS':
## 
##     geyser
```

```
S3 <- read.delim(file="S3.txt")
S3 <- S3[which(S3$Phenotype_analysis==1),]###only keep Individuals with unique sampling location
head(S3)
```

```
##               Name   ID          Region Country Final_Size_HL  T50_HL Slope_HL
## 1  Ale-Stenar-41-1  991 Northern Europe  Sweden        8.6731 10.5321   5.7169
## 5   Brösarp-11-135 1061 Northern Europe  Sweden        7.1276 15.5622   4.9051
## 9          App1-12 5830 Northern Europe  Sweden        6.0112 12.8023   4.0086
## 15          Fly2-1 6023 Northern Europe  Sweden        6.2978 13.4606   4.7825
## 17         Hov1-10 6035 Northern Europe  Sweden        6.2719 17.7050   5.8510
## 21           Kni-1 6040 Northern Europe  Sweden        6.8210 12.3300   3.9022
##    Final_Size_LL  T50_LL Slope_LL GXE_Final_Size GXE_t50 GXE_Slope
## 1        14.5374 34.5172   5.1963        -0.7409  0.4742   -1.6368
## 5        11.4213 35.4196   5.7678        -2.3115 -3.6535   -0.2535
## 9        13.0435 33.7565   5.2702         0.4271 -2.5568    0.1453
## 15       13.1835 38.0945   5.5757         0.2806  1.1229   -0.3229
## 17       12.7316 38.0023   4.3531        -0.1455 -3.2136   -2.6141
## 21       11.4780 34.1555   4.0908        -1.9482 -1.6854   -0.9276
##    Phenotype_analysis GWAS DiamFieldM2 Hypocotyllength Biomass21d  FT_10
## 1                   1    1          NA              NA         NA 110.50
## 5                   1    1    9.499000               7     0.0218  97.75
## 9                   1    1    7.179917               4     0.0180  97.75
## 15                  1    1   11.961833               3     0.0330  93.75
## 17                  1    1    9.728250               4     0.0201  83.00
## 21                  1    1    7.603167               4     0.0385  93.25
##        FT_16 FT_HL FT_LL
## 1  106.75000    NA    NA
## 5   94.75000    NA    NA
## 9   75.33333    NA    90
## 15  89.25000    NA    NA
## 17  73.75000    NA    NA
## 21  82.50000    NA    NA
```

```
#HL
glm_FSHL <- glm(Final_Size_HL ~ Region-1 , data=S3)
summary(glht(glm_FSHL, mcp(Region="Tukey")))
```

```
## 
##   Simultaneous Tests for General Linear Hypotheses
## 
## Multiple Comparisons of Means: Tukey Contrasts
## 
## 
## Fit: glm(formula = Final_Size_HL ~ Region - 1, data = S3)
## 
## Linear Hypotheses:
##                                        Estimate Std. Error z value Pr(>|z|)    
## Northern Europe - China == 0            -0.2379     0.2923  -0.814    0.913    
## Spain - China == 0                       0.6019     0.2760   2.181    0.157    
## Western  Europe - China == 0             0.2215     1.0402   0.213    0.999    
## Western Europe - China == 0             -0.3502     0.3223  -1.087    0.785    
## Spain - Northern Europe == 0             0.8398     0.1622   5.176   <0.001 ***
## Western  Europe - Northern Europe == 0   0.4594     1.0160   0.452    0.990    
## Western Europe - Northern Europe == 0   -0.1123     0.2324  -0.483    0.987    
## Western  Europe - Spain == 0            -0.3804     1.0114  -0.376    0.995    
## Western Europe - Spain == 0             -0.9521     0.2116  -4.500   <0.001 ***
## Western Europe - Western  Europe == 0   -0.5717     1.0250  -0.558    0.977    
## ---
## Signif. codes:  0 '***' 0.001 '**' 0.01 '*' 0.05 '.' 0.1 ' ' 1
## (Adjusted p values reported -- single-step method)
```

```
glm_t50HL <- glm(T50_HL ~ Region-1 , data=S3)
summary(glht(glm_t50HL, mcp(Region="Tukey")))
```

```
## 
##   Simultaneous Tests for General Linear Hypotheses
## 
## Multiple Comparisons of Means: Tukey Contrasts
## 
## 
## Fit: glm(formula = T50_HL ~ Region - 1, data = S3)
## 
## Linear Hypotheses:
##                                        Estimate Std. Error z value Pr(>|z|)  
## Northern Europe - China == 0            -1.1470     0.8169  -1.404   0.5824  
## Spain - China == 0                       0.2002     0.7713   0.260   0.9988  
## Western  Europe - China == 0            -2.9147     2.9072  -1.003   0.8305  
## Western Europe - China == 0             -0.2670     0.9007  -0.296   0.9979  
## Spain - Northern Europe == 0             1.3473     0.4534   2.971   0.0194 *
## Western  Europe - Northern Europe == 0  -1.7677     2.8395  -0.623   0.9657  
## Western Europe - Northern Europe == 0    0.8800     0.6496   1.355   0.6159  
## Western  Europe - Spain == 0            -3.1149     2.8267  -1.102   0.7763  
## Western Europe - Spain == 0             -0.4673     0.5912  -0.790   0.9211  
## Western Europe - Western  Europe == 0    2.6477     2.8647   0.924   0.8683  
## ---
## Signif. codes:  0 '***' 0.001 '**' 0.01 '*' 0.05 '.' 0.1 ' ' 1
## (Adjusted p values reported -- single-step method)
```

```
glm_SLHL <- glm(Slope_HL ~ Region-1 , data=S3)
summary(glht(glm_SLHL, mcp(Region="Tukey")))
```

```
## 
##   Simultaneous Tests for General Linear Hypotheses
## 
## Multiple Comparisons of Means: Tukey Contrasts
## 
## 
## Fit: glm(formula = Slope_HL ~ Region - 1, data = S3)
## 
## Linear Hypotheses:
##                                        Estimate Std. Error z value Pr(>|z|)  
## Northern Europe - China == 0            0.54140    0.33211   1.630   0.4324  
## Spain - China == 0                      0.80160    0.31357   2.556   0.0635 .
## Western  Europe - China == 0            0.31813    1.18198   0.269   0.9986  
## Western Europe - China == 0             0.37147    0.36619   1.014   0.8244  
## Spain - Northern Europe == 0            0.26019    0.18435   1.411   0.5774  
## Western  Europe - Northern Europe == 0 -0.22328    1.15445  -0.193   0.9996  
## Western Europe - Northern Europe == 0  -0.16993    0.26411  -0.643   0.9614  
## Western  Europe - Spain == 0           -0.48347    1.14925  -0.421   0.9920  
## Western Europe - Spain == 0            -0.43013    0.24038  -1.789   0.3363  
## Western Europe - Western  Europe == 0   0.05334    1.16471   0.046   1.0000  
## ---
## Signif. codes:  0 '***' 0.001 '**' 0.01 '*' 0.05 '.' 0.1 ' ' 1
## (Adjusted p values reported -- single-step method)
```

```
#LL
glm_FSLL <- glm(Final_Size_LL ~ Region-1 , data=S3)
summary(glht(glm_FSLL, mcp(Region="Tukey")))
```

```
## 
##   Simultaneous Tests for General Linear Hypotheses
## 
## Multiple Comparisons of Means: Tukey Contrasts
## 
## 
## Fit: glm(formula = Final_Size_LL ~ Region - 1, data = S3)
## 
## Linear Hypotheses:
##                                        Estimate Std. Error z value Pr(>|z|)    
## Northern Europe - China == 0            0.29729    0.41417   0.718   0.9435    
## Spain - China == 0                      1.58276    0.38671   4.093   <0.001 ***
## Western  Europe - China == 0            0.51082    1.60406   0.318   0.9973    
## Western Europe - China == 0             0.57157    0.46145   1.239   0.6934    
## Spain - Northern Europe == 0            1.28546    0.25254   5.090   <0.001 ***
## Western  Europe - Northern Europe == 0  0.21352    1.57710   0.135   0.9999    
## Western Europe - Northern Europe == 0   0.27428    0.35661   0.769   0.9284    
## Western  Europe - Spain == 0           -1.07194    1.57011  -0.683   0.9526    
## Western Europe - Spain == 0            -1.01119    0.32432  -3.118   0.0124 *  
## Western Europe - Western  Europe == 0   0.06076    1.59017   0.038   1.0000    
## ---
## Signif. codes:  0 '***' 0.001 '**' 0.01 '*' 0.05 '.' 0.1 ' ' 1
## (Adjusted p values reported -- single-step method)
```

```
glm_t50LL <- glm(T50_LL ~ Region-1 , data=S3)
summary(glht(glm_t50LL, mcp(Region="Tukey")))
```

```
## 
##   Simultaneous Tests for General Linear Hypotheses
## 
## Multiple Comparisons of Means: Tukey Contrasts
## 
## 
## Fit: glm(formula = T50_LL ~ Region - 1, data = S3)
## 
## Linear Hypotheses:
##                                        Estimate Std. Error z value Pr(>|z|)    
## Northern Europe - China == 0             1.1603     0.7398   1.568  0.47385    
## Spain - China == 0                       2.8663     0.6907   4.150  < 0.001 ***
## Western  Europe - China == 0             3.2611     2.8652   1.138  0.75587    
## Western Europe - China == 0              2.9150     0.8243   3.537  0.00314 ** 
## Spain - Northern Europe == 0             1.7060     0.4511   3.782  0.00119 ** 
## Western  Europe - Northern Europe == 0   2.1009     2.8171   0.746  0.93556    
## Western Europe - Northern Europe == 0    1.7547     0.6370   2.755  0.03723 *  
## Western  Europe - Spain == 0             0.3948     2.8046   0.141  0.99989    
## Western Europe - Spain == 0              0.0487     0.5793   0.084  0.99999    
## Western Europe - Western  Europe == 0   -0.3461     2.8404  -0.122  0.99994    
## ---
## Signif. codes:  0 '***' 0.001 '**' 0.01 '*' 0.05 '.' 0.1 ' ' 1
## (Adjusted p values reported -- single-step method)
```

```
glm_SLLL <- glm(Slope_LL ~ Region-1 , data=S3)
summary(glht(glm_SLLL, mcp(Region="Tukey")))
```

```
## 
##   Simultaneous Tests for General Linear Hypotheses
## 
## Multiple Comparisons of Means: Tukey Contrasts
## 
## 
## Fit: glm(formula = Slope_LL ~ Region - 1, data = S3)
## 
## Linear Hypotheses:
##                                        Estimate Std. Error z value Pr(>|z|)
## Northern Europe - China == 0           -0.08681    0.19279  -0.450    0.990
## Spain - China == 0                     -0.31456    0.18001  -1.748    0.362
## Western  Europe - China == 0           -0.40127    0.74666  -0.537    0.980
## Western Europe - China == 0            -0.37493    0.21480  -1.745    0.363
## Spain - Northern Europe == 0           -0.22775    0.11755  -1.937    0.259
## Western  Europe - Northern Europe == 0 -0.31446    0.73411  -0.428    0.992
## Western Europe - Northern Europe == 0  -0.28811    0.16600  -1.736    0.369
## Western  Europe - Spain == 0           -0.08671    0.73086  -0.119    1.000
## Western Europe - Spain == 0            -0.06036    0.15096  -0.400    0.993
## Western Europe - Western  Europe == 0   0.02635    0.74020   0.036    1.000
## (Adjusted p values reported -- single-step method)
```

```
#GXE
glm_FSGXE <- glm(GXE_Final_Size ~ Region-1 , data=S3)
summary(glht(glm_FSGXE, mcp(Region="Tukey")))
```

```
## 
##   Simultaneous Tests for General Linear Hypotheses
## 
## Multiple Comparisons of Means: Tukey Contrasts
## 
## 
## Fit: glm(formula = GXE_Final_Size ~ Region - 1, data = S3)
## 
## Linear Hypotheses:
##                                        Estimate Std. Error z value Pr(>|z|)
## Northern Europe - China == 0             0.3748     0.4698   0.798    0.919
## Spain - China == 0                       0.8151     0.4455   1.830    0.313
## Western  Europe - China == 0             0.1289     1.6305   0.079    1.000
## Western Europe - China == 0              0.6267     0.5156   1.215    0.707
## Spain - Northern Europe == 0             0.4403     0.2544   1.731    0.370
## Western  Europe - Northern Europe == 0  -0.2459     1.5889  -0.155    1.000
## Western Europe - Northern Europe == 0    0.2519     0.3635   0.693    0.950
## Western  Europe - Spain == 0            -0.6862     1.5819  -0.434    0.991
## Western Europe - Spain == 0             -0.1884     0.3314  -0.569    0.975
## Western Europe - Western  Europe == 0    0.4978     1.6031   0.311    0.998
## (Adjusted p values reported -- single-step method)
```

```
glm_t50GXE <- glm(GXE_t50 ~ Region-1 , data=S3)
summary(glht(glm_t50GXE, mcp(Region="Tukey")))
```

```
## 
##   Simultaneous Tests for General Linear Hypotheses
## 
## Multiple Comparisons of Means: Tukey Contrasts
## 
## 
## Fit: glm(formula = GXE_t50 ~ Region - 1, data = S3)
## 
## Linear Hypotheses:
##                                        Estimate Std. Error z value Pr(>|z|)  
## Northern Europe - China == 0             2.4196     1.0844   2.231   0.1404  
## Spain - China == 0                       2.8257     1.0281   2.748   0.0374 *
## Western  Europe - China == 0             6.2882     3.7632   1.671   0.4065  
## Western Europe - China == 0              3.1594     1.1900   2.655   0.0485 *
## Spain - Northern Europe == 0             0.4061     0.5872   0.691   0.9502  
## Western  Europe - Northern Europe == 0   3.8686     3.6673   1.055   0.8028  
## Western Europe - Northern Europe == 0    0.7398     0.8390   0.882   0.8866  
## Western  Europe - Spain == 0             3.4625     3.6511   0.948   0.8571  
## Western Europe - Spain == 0              0.3337     0.7649   0.436   0.9908  
## Western Europe - Western  Europe == 0   -3.1287     3.6999  -0.846   0.9010  
## ---
## Signif. codes:  0 '***' 0.001 '**' 0.01 '*' 0.05 '.' 0.1 ' ' 1
## (Adjusted p values reported -- single-step method)
```

```
glm_SLGXE <- glm(GXE_Slope ~ Region-1 , data=S3)
summary(glht(glm_SLGXE, mcp(Region="Tukey")))
```

```
## 
##   Simultaneous Tests for General Linear Hypotheses
## 
## Multiple Comparisons of Means: Tukey Contrasts
## 
## 
## Fit: glm(formula = GXE_Slope ~ Region - 1, data = S3)
## 
## Linear Hypotheses:
##                                        Estimate Std. Error z value Pr(>|z|)   
## Northern Europe - China == 0           -0.71025    0.39534  -1.797  0.33212   
## Spain - China == 0                     -1.20577    0.37482  -3.217  0.00884 **
## Western  Europe - China == 0           -0.80145    1.37192  -0.584  0.97277   
## Western Europe - China == 0            -0.82951    0.43384  -1.912  0.27024   
## Spain - Northern Europe == 0           -0.49552    0.21409  -2.315  0.11595   
## Western  Europe - Northern Europe == 0 -0.09120    1.33698  -0.068  0.99999   
## Western Europe - Northern Europe == 0  -0.11927    0.30587  -0.390  0.99404   
## Western  Europe - Spain == 0            0.40432    1.33105   0.304  0.99773   
## Western Europe - Spain == 0             0.37625    0.27884   1.349  0.61896   
## Western Europe - Western  Europe == 0  -0.02806    1.34886  -0.021  1.00000   
## ---
## Signif. codes:  0 '***' 0.001 '**' 0.01 '*' 0.05 '.' 0.1 ' ' 1
## (Adjusted p values reported -- single-step method)
```

```
#Diameter Field:
glm_Diam <- glm(DiamFieldM2 ~ Region-1 , data=S3)
summary(glht(glm_Diam, mcp(Region="Tukey")))
```

```
## 
##   Simultaneous Tests for General Linear Hypotheses
## 
## Multiple Comparisons of Means: Tukey Contrasts
## 
## 
## Fit: glm(formula = DiamFieldM2 ~ Region - 1, data = S3)
## 
## Linear Hypotheses:
##                                        Estimate Std. Error z value Pr(>|z|)    
## Northern Europe - China == 0            -1.3779     0.7756  -1.777  0.34419    
## Spain - China == 0                      -0.1960     0.7447  -0.263  0.99871    
## Western  Europe - China == 0             0.1735     2.3443   0.074  0.99999    
## Western Europe - China == 0             -3.2843     0.8274  -3.969  < 0.001 ***
## Spain - Northern Europe == 0             1.1820     0.3961   2.984  0.01843 *  
## Western  Europe - Northern Europe == 0   1.5514     2.2579   0.687  0.95148    
## Western Europe - Northern Europe == 0   -1.9063     0.5357  -3.558  0.00244 ** 
## Western  Europe - Spain == 0             0.3695     2.2474   0.164  0.99980    
## Western Europe - Spain == 0             -3.0883     0.4898  -6.305  < 0.001 ***
## Western Europe - Western  Europe == 0   -3.4578     2.2762  -1.519  0.50572    
## ---
## Signif. codes:  0 '***' 0.001 '**' 0.01 '*' 0.05 '.' 0.1 ' ' 1
## (Adjusted p values reported -- single-step method)
```

### Plot Figures 2a-d (does not work in the html, but in the Rscript)

### Plot Fig. S8

###### Plot Fig. S11

'

###### Heritability

```
library(nlme)

data <- read.delim(file = "~/Desktop/Bene/Documents/Protocols_publications/2020_PlosGenetics/uptodate326/S2.txt")
data$Replicate <- as.factor(data$Replicate)
S3 <- read.delim(file="/home/bene/Desktop/Bene/Documents/Protocols_publications/2020_PlosGenetics/uptodate326/S3.txt")
S3 <- S3[which(S3$Phenotype_analysis==1),]###only keep Individuals with unique sampling location

#separate treatments and remove duplicate origins
hl <- data[which(data$treatment=="HL" & data$ID %in% S3$ID),]
ll <- data[which(data$treatment=="LL"),]


#Broad-Sense Heritability (H²) of FSHL
h2.model_FSHL <- lme(fixed = Final.Size ~ Replicate, random = ~ 1 | ID, data= hl)
VarCorr(h2.model_FSHL)
```

```
## ID = pdLogChol(1) 
##             Variance  StdDev   
## (Intercept) 0.7542641 0.8684838
## Residual    0.5711426 0.7557398
```

```
plot(h2.model_FSHL)
```

```
hist(hl$Final.Size)
```

```
sigma2.g_FSHL <- as.numeric(VarCorr(h2.model_FSHL)[1])^2###Intercept variance
sigma2.e_FSHL <- as.numeric(VarCorr(h2.model_FSHL)[2])^2###Residual Variance
h2_FSHL <- sigma2.g_FSHL / (sigma2.g_FSHL + sigma2.e_FSHL)
h2_FSHL #0.6355745
```

```
## [1] 0.6355745
```

```
#####same script for all heritabilities
```

##### Correlation among experiments

```
#estimate the correlation matrix with corrected p-values
require(nlme)
library(coxme)
```

```
## Loading required package: bdsmatrix
```

```
## 
## Attaching package: 'bdsmatrix'
```

```
## The following object is masked from 'package:base':
## 
##     backsolve
```

```
data <- read.delim(file = "S2.txt")
data$Replicate <- as.factor(data$Replicate)


S3 <- read.delim(file="S3.txt")
head(S3)
```

```
##               Name   ID          Region Country Final_Size_HL  T50_HL Slope_HL
## 1  Ale-Stenar-41-1  991 Northern Europe  Sweden        8.6731 10.5321   5.7169
## 2  Ale-Stenar-44-4  992 Northern Europe  Sweden        7.2023 11.8398   4.8478
## 3 Ale-Stenar-56-14  997 Northern Europe  Sweden        6.6369 15.5743   4.0614
## 4 Ale-Stenar-64-24 1002 Northern Europe  Sweden        7.6601 13.7759   3.7600
## 5   Brösarp-11-135 1061 Northern Europe  Sweden        7.1276 15.5622   4.9051
## 6   Brösarp-15-138 1062 Northern Europe  Sweden        5.8695 16.4978   3.7495
##   Final_Size_LL  T50_LL Slope_LL GXE_Final_Size GXE_t50 GXE_Slope
## 1       14.5374 34.5172   5.1963        -0.7409  0.4742   -1.6368
## 2       13.7532 33.5521   5.5908        -0.0543 -1.7986   -0.3731
## 3       15.2798 32.4278   5.6417         2.0377 -6.6575    0.4641
## 4       14.2653 37.2868   4.8761        -1.1611 -0.5669    0.9047
## 5       11.4213 35.4196   5.7678        -2.3115 -3.6535   -0.2535
## 6       11.0332 31.6869   5.6354        -1.4415 -8.3219    0.7698
##   Phenotype_analysis GWAS DiamFieldM2 Hypocotyllength Biomass21d  FT_10  FT_16
## 1                  1    1          NA              NA         NA 110.50 106.75
## 2                  0    1          NA              NA         NA 116.00 100.00
## 3                  0    1   14.375000               5     0.0343 101.25  87.50
## 4                  0    1    8.069833              10     0.0239 102.75  86.25
## 5                  1    1    9.499000               7     0.0218  97.75  94.75
## 6                  0    1   10.855333               4     0.0301  94.00  68.25
##   FT_HL FT_LL
## 1    NA    NA
## 2    NA    NA
## 3    NA  81.0
## 4    NA    NA
## 5    NA    NA
## 6    NA  88.5
```

```
S3 <- S3[which(S3$Phenotype_analysis==1),]###only keep Individuals with unique sampling location
S3 <- S3[-which(S3$ID %in% c("9580","9829","Ahthx-23","Xyl")),]#no genomic data
corrset_14 <- S3[-c(which(S3$Region=="Western Europe")), c("ID","Final_Size_HL","T50_HL","Slope_HL","FT_HL","Final_Size_LL","T50_LL","Slope_LL","FT_LL","GXE_Final_Size","GXE_t50","GXE_Slope","DiamFieldM2","Hypocotyllength","Biomass21d")]#traits to be correlated without the french accessions (no genomic data)
colnames(corrset_14) <- c("ID","FS_HL","T50_HL","SL_HL","FT_HL","FS_LL","T50_LL","SL_LL","FT_LL","GXE_FS","GXE_T50","GXE_SL","DiamFieldM2","Hypocotyllength","Biomass21d")
##replace chinese names:
corrset_14$ID <- as.numeric(as.character(corrset_14$ID))
```

```
## Warning: NAs introduced by coercion
```

```
corrset_14[176:193,"ID"] <- c("2189710","2189718","2204175","2204176","2204177","6414609","2204317","2204239","2204174",
"2204313","2204161","2204162","6414604","6414610","2204626","2204179","2204168","2204170")

#Kinship matrix
load("K_227_filtered.R")


#estimate pearson correlation
correlation_lmekin14 <- cor(corrset_14[,2:15],use = "pairwise.complete.obs",method="pearson")


pvalues_lmekin14 <- matrix(data = NA,nrow = 14,ncol=14)

####lmekin for p-values
for (i in 2:ncol(corrset_14)){
  for (j in i:ncol(corrset_14)){
    traitcorr <- lmekin(corrset_14[,i]~corrset_14[,j]+ (1|corrset_14$ID) ,method="ML",varlist=K_227_filtered,na.action = na.omit)
    #extract the correlation and pvalue:
    tabli<-capture.output(print(traitcorr)[0])#extract model output
    g<-strsplit (tabli[10], "\\s+")
    pval<-g[[1]][length(g[[1]])]
    pvalues_lmekin14[i-1,j-1]<-pval
    pvalues_lmekin14[i-1,i-1]<-0
  }
}

rownames(correlation_lmekin14) <- colnames(corrset_14)[2:15]
colnames(correlation_lmekin14) <- colnames(corrset_14)[2:15]
correlation_lmekin14
```

```
##                        FS_HL      T50_HL        SL_HL       FT_HL       FS_LL
## FS_HL            1.000000000 -0.24209833 -0.008092318 -0.24059653  0.38932980
## T50_HL          -0.242098328  1.00000000  0.050643698  0.10599831 -0.10807403
## SL_HL           -0.008092318  0.05064370  1.000000000  0.33710461  0.18383677
## FT_HL           -0.240596529  0.10599831  0.337104611  1.00000000  0.26051421
## FS_LL            0.389329796 -0.10807403  0.183836770  0.26051421  1.00000000
## T50_LL           0.005883554  0.18476805  0.157651117  0.20708348  0.43028008
## SL_LL           -0.106957146 -0.04322218  0.064815303  0.06382357 -0.29646732
## FT_LL           -0.143772080  0.05736238  0.365044334  0.63131820  0.32395613
## GXE_FS          -0.273552600  0.05335189  0.195474917  0.42172222  0.77946283
## GXE_T50          0.193932150 -0.62821153  0.082091543  0.10527363  0.41392426
## GXE_SL          -0.054208822 -0.07183249 -0.836696362 -0.24974865 -0.31663736
## DiamFieldM2      0.262926201 -0.01961356  0.072612423 -0.32196202  0.09765287
## Hypocotyllength  0.372065615  0.05828829  0.127093986  0.35695874  0.31625183
## Biomass21d       0.267081752 -0.17280739  0.049888330 -0.32692546  0.02861364
##                       T50_LL       SL_LL       FT_LL      GXE_FS      GXE_T50
## FS_HL            0.005883554 -0.10695715 -0.14377208 -0.27355260  0.193932150
## T50_HL           0.184768050 -0.04322218  0.05736238  0.05335189 -0.628211526
## SL_HL            0.157651117  0.06481530  0.36504433  0.19547492  0.082091543
## FT_HL            0.207083478  0.06382357  0.63131820  0.42172222  0.105273634
## FS_LL            0.430280077 -0.29646732  0.32395613  0.77946283  0.413924259
## T50_LL           1.000000000 -0.36462554  0.31451874  0.43157881  0.648572961
## SL_LL           -0.364625536  1.00000000 -0.05482461 -0.22461604 -0.249454427
## FT_LL            0.314518739 -0.05482461  1.00000000  0.40942246  0.219796638
## GXE_FS           0.431578806 -0.22461604  0.40942246  1.00000000  0.300348699
## GXE_T50          0.648572961 -0.24945443  0.21979664  0.30034870  1.000000000
## GXE_SL          -0.333652084  0.49228473 -0.34724325 -0.29378309 -0.208510282
## DiamFieldM2     -0.023039321  0.02252715 -0.32168176 -0.07706916  0.008289135
## Hypocotyllength  0.247096480 -0.22025945  0.24407457  0.06687632  0.141296054
## Biomass21d      -0.076387754  0.02534068 -0.21191143 -0.17362617  0.076510579
##                      GXE_SL  DiamFieldM2 Hypocotyllength  Biomass21d
## FS_HL           -0.05420882  0.262926201      0.37206561  0.26708175
## T50_HL          -0.07183249 -0.019613557      0.05828829 -0.17280739
## SL_HL           -0.83669636  0.072612423      0.12709399  0.04988833
## FT_HL           -0.24974865 -0.321962021      0.35695874 -0.32692546
## FS_LL           -0.31663736  0.097652869      0.31625183  0.02861364
## T50_LL          -0.33365208 -0.023039321      0.24709648 -0.07638775
## SL_LL            0.49228473  0.022527154     -0.22025945  0.02534068
## FT_LL           -0.34724325 -0.321681763      0.24407457 -0.21191143
## GXE_FS          -0.29378309 -0.077069162      0.06687632 -0.17362617
## GXE_T50         -0.20851028  0.008289135      0.14129605  0.07651058
## GXE_SL           1.00000000 -0.059569637     -0.23439691 -0.02964972
## DiamFieldM2     -0.05956964  1.000000000      0.13276868  0.20862884
## Hypocotyllength -0.23439691  0.132768676      1.00000000  0.02283247
## Biomass21d      -0.02964972  0.208628843      0.02283247  1.00000000
```

```
pvalues_lmekin14
```

```
##       [,1] [,2]    [,3]   [,4]      [,5]      [,6]      [,7]      [,8]     
##  [1,] "0"  "6e-04" "0.91" "2.8e-02" "7.5e-09" "9.4e-01" "0.14"    "0.24"   
##  [2,] NA   "0"     "0.49" "3.9e-01" "0.14"    "0.01000" "0.55"    "6.4e-01"
##  [3,] NA   NA      "0"    "0.0042"  "1.1e-02" "0.029"   "3.7e-01" "1.4e-03"
##  [4,] NA   NA      NA     "0"       "3.2e-02" "0.0930"  "0.61"    "1.1e-09"
##  [5,] NA   NA      NA     NA        "0"       "4.5e-11" "1.8e-05" "0.0042" 
##  [6,] NA   NA      NA     NA        NA        "0"       "6.2e-08" "0.0056" 
##  [7,] NA   NA      NA     NA        NA        NA        "0"       "0.65"   
##  [8,] NA   NA      NA     NA        NA        NA        NA        "0"      
##  [9,] NA   NA      NA     NA        NA        NA        NA        NA       
## [10,] NA   NA      NA     NA        NA        NA        NA        NA       
## [11,] NA   NA      NA     NA        NA        NA        NA        NA       
## [12,] NA   NA      NA     NA        NA        NA        NA        NA       
## [13,] NA   NA      NA     NA        NA        NA        NA        NA       
## [14,] NA   NA      NA     NA        NA        NA        NA        NA       
##       [,9]      [,10]     [,11]     [,12]     [,13]     [,14]    
##  [1,] "1e-04"   "0.0069"  "0.46"    "0.00088" "9.1e-07" "4.6e-05"
##  [2,] "0.47"    "0"       "0.32"    "0.81"    "0.47"    "0.032"  
##  [3,] "0.0064"  "0.26"    "0"       "0.37"    "0.12"    "0.54"   
##  [4,] "0.00022" "0.4"     "0.041"   "1.5e-04" "0.0042"  "0.01"   
##  [5,] "0"       "5e-10"   "5e-06"   "0.23"    "3.7e-05" "0.72"   
##  [6,] "6.1e-11" "0"       "1.3e-06" "0.78"    "0.0016"  "0.34"   
##  [7,] "0.0016"  "0.00043" "1e-14"   "0.78"    "0.0052"  "0.75"   
##  [8,] "0.00027" "0.067"   "0.0026"  "4.9e-03" "0.049"   "0.093"  
##  [9,] "0"       "1.7e-05" "2.6e-05" "0.35"    "0.4100"  "0.032"  
## [10,] NA        "0"       "0.0036"  "0.920"   "0.043"   "0.35000"
## [11,] NA        NA        "0"       "0.47"    "0.0032"  "0.72"   
## [12,] NA        NA        NA        "0"       "2.0e-01" "9.7e-03"
## [13,] NA        NA        NA        NA        "0"       "0.87000"
## [14,] NA        NA        NA        NA        NA        "0"
```

```
corr <- apply(correlation_lmekin14,FUN=as.numeric,MARGIN = 2)
rownames(corr) <- colnames(corrset_14)[2:15]
pval <- apply(pvalues_lmekin14,FUN=as.numeric,MARGIN = 2)
pval.adj <- matrix(p.adjust(t(pval),method = "fdr"),nrow=14,ncol=14,byrow=T)#correct for multiple testing
```

###### Fig S18

```
###Plot
library(corrplot)
```

```
## corrplot 0.84 loaded
```

```
col1 <- colorRampPalette(c("blue4", "white", "red2"))#blue to red, but pretty versions of it

corrplot(corr=t(corr),method="square", type="lower", p.mat =t(pval.adj),sig.level=0.05, diag=F,
         insig = "blank",tl.col="black", mar = c(1,1,1,1),col=col1(200),tl.pos="ld")
```

###### Prepare data for Fig S23

```
#climate data from table S1 has to be merged with genotypic means from table S3:
S1 <- read.delim("S1.txt")
head(S1)
```

```
##   Country Region Genotype X1001GenomesID                 Location Latitude
## 1   China  China Ahthx-23             NA         Anhui, Taihuxian 30.45222
## 2   China  China   Aqs-11             NA      Anhui, Qianshanxian 30.74600
## 3   China  China      Ayx             NA         Anhui, Yuexixian 30.42890
## 4   China  China      Cbb             NA      Chongqing, Beibeiqu 29.79400
## 5   China  China   Ctl-62             NA Chongqing, Tongliangxian 29.82300
## 6   China  China   Gwx-11             NA           Gansu, Wenxian 32.72500
##   Longitude          Collector  X Nr_Growing.months   T_GS  SWC_GS
## 1  117.2856 Fei He, Hongya  Gu NA                11 17.597  91.818
## 2  117.9730 Fei He, Hongya  Gu NA                11 17.449  92.091
## 3  116.1533 Fei He, Hongya  Gu NA                11 16.668 100.000
## 4  106.4840 Fei He, Hongya  Gu NA                12 17.473  83.917
## 5  106.0560 Fei He, Hongya  Gu NA                12 17.993  85.000
## 6  105.1220 Fei He, Hongya  Gu NA                10 15.926  54.300
##   Water.vapor_GS Wind_GS Radiation_GS Rain_GS Bioclim1_T_Annual
## 1          1.687   2.373     15323.73 134.909            16.527
## 2          1.719   2.465     15474.27 120.182            16.340
## 3          1.527   2.381     14992.73 136.273            15.583
## 4          1.668   1.339     12056.67  94.583            17.472
## 5          1.728   1.310     11824.75  92.500            17.993
## 6          1.329   1.339     13477.30  72.900            13.942
##   Bioclim2_Diurnal_Range Bioclim3_Isothermality Bioclim4_T_Seasonality
## 1                  8.887                 27.273                886.441
## 2                  8.977                 26.919                901.863
## 3                  7.790                 24.690                864.066
## 4                  6.545                 24.014                744.532
## 5                  6.443                 23.809                738.784
## 6                  8.807                 30.458                725.205
##   Bioclim5_MaxT_Wrmst_Month Bioclim_MinT_Cldst_Month Bioclim7_T_Ann_Range
## 1                    33.332                    0.748               32.584
## 2                    33.552                    0.204               33.348
## 3                    31.536                   -0.016               31.552
## 4                    30.280                    3.024               27.256
## 5                    30.732                    3.672               27.060
## 6                    27.624                   -1.292               28.916
##   Bioclim8_T_Wettest_Qtr Bioclim9_T_Driest_Qtr Bioclim10_T_Wrmst_Qtr
## 1                 20.335                 7.347                27.296
## 2                 24.735                 6.918                27.303
## 3                 23.857                 6.713                26.031
## 4                 26.378                 7.925                26.378
## 5                 26.843                 8.499                26.843
## 6                 21.773                 4.587                22.594
##   Bioclim11_T_Cldst_Qtr Bioclim12_Ann_Precip Bioclim13_P_Wettest_Month
## 1                 5.604                 1521                       287
## 2                 5.131                 1358                       238
## 3                 4.766                 1538                       273
## 4                 7.925                 1135                       192
## 5                 8.499                 1110                       203
## 6                 4.587                  734                       167
##   Bioclim14_P_Driest_Month Bioclim15_P_Seasonality Bioclim16_P_Wettest_Qtr
## 1                       33                  55.281                     637
## 2                       32                  51.795                     555
## 3                       36                  55.307                     649
## 4                       17                  69.233                     522
## 5                       17                  73.259                     539
## 6                        2                  92.797                     426
##   Bioclim17_P_Driest_Qtr Bioclim18_P_Wrmst_Qtr Bioclim19_P_Cldst_Qtr Radiation
## 1                    145                   597                   148  15323.73
## 2                    135                   523                   141  15474.27
## 3                    145                   632                   150  14992.73
## 4                     57                   522                    57  12056.67
## 5                     56                   539                    56  11824.75
## 6                     10                   403                    10  13477.30
##   PC1_growS PC2_growS   PC1_T  PC2_T   PC1_P    PC2_P
## 1    76.637   -10.831 252.659 -2.721 934.429 -283.015
## 2    63.727    -3.774 268.271 -2.563 736.026 -242.715
## 3    81.620    -4.205 230.459  0.127 960.713 -304.008
## 4    37.293     0.734 111.269 -1.608 504.260 -327.447
## 5    35.966     2.640 105.582 -2.603 494.650 -346.619
## 6     4.270   -15.280  91.504  2.529  78.061 -318.699
```

```
S3 <- read.delim("S3.txt")
head(S3)
```

```
##               Name   ID          Region Country Final_Size_HL  T50_HL Slope_HL
## 1  Ale-Stenar-41-1  991 Northern Europe  Sweden        8.6731 10.5321   5.7169
## 2  Ale-Stenar-44-4  992 Northern Europe  Sweden        7.2023 11.8398   4.8478
## 3 Ale-Stenar-56-14  997 Northern Europe  Sweden        6.6369 15.5743   4.0614
## 4 Ale-Stenar-64-24 1002 Northern Europe  Sweden        7.6601 13.7759   3.7600
## 5   Brösarp-11-135 1061 Northern Europe  Sweden        7.1276 15.5622   4.9051
## 6   Brösarp-15-138 1062 Northern Europe  Sweden        5.8695 16.4978   3.7495
##   Final_Size_LL  T50_LL Slope_LL GXE_Final_Size GXE_t50 GXE_Slope
## 1       14.5374 34.5172   5.1963        -0.7409  0.4742   -1.6368
## 2       13.7532 33.5521   5.5908        -0.0543 -1.7986   -0.3731
## 3       15.2798 32.4278   5.6417         2.0377 -6.6575    0.4641
## 4       14.2653 37.2868   4.8761        -1.1611 -0.5669    0.9047
## 5       11.4213 35.4196   5.7678        -2.3115 -3.6535   -0.2535
## 6       11.0332 31.6869   5.6354        -1.4415 -8.3219    0.7698
##   Phenotype_analysis GWAS DiamFieldM2 Hypocotyllength Biomass21d  FT_10  FT_16
## 1                  1    1          NA              NA         NA 110.50 106.75
## 2                  0    1          NA              NA         NA 116.00 100.00
## 3                  0    1   14.375000               5     0.0343 101.25  87.50
## 4                  0    1    8.069833              10     0.0239 102.75  86.25
## 5                  1    1    9.499000               7     0.0218  97.75  94.75
## 6                  0    1   10.855333               4     0.0301  94.00  68.25
##   FT_HL FT_LL
## 1    NA    NA
## 2    NA    NA
## 3    NA  81.0
## 4    NA    NA
## 5    NA    NA
## 6    NA  88.5
```

```
#only keep necessary columns:
S1a <- S1[,c("Region","Genotype","X1001GenomesID","Latitude","Radiation","PC1_growS","PC2_growS","PC1_T","PC2_T","PC1_P","PC2_P")]

#chinese IDs have to be added for merging:
colnames(S1a)[3] <- "ID"
S1a$ID <- as.character(S1a$ID)
S1a[which(is.na(S1a$ID==T)),"ID"] <- as.character(S1a[which(is.na(S1a$ID==T)),"Genotype"])
##replace chinese names with numbers:

S1a[which(S1a$Region=="China"),"ID"] <- c("6414606","2189710","2189718","2204175","2204176","2204177","6414609","2204317","2204239","2204174",
"2204313","2204161","2204162","6414604","6414610",NA,"2204626","2204179",NA,"2204168","2204170")

S1a$ID <- as.numeric(as.character(S1a$ID))
```

```
## Warning: NAs introduced by coercion
```

```
S1a <- S1a[-which(is.na(S1a$ID)==T),]
S1a$ID <- as.factor(S1a$ID)

#merge datasets
S19 <- merge(S1a,S3,by="ID",all = T)
head(S19)
```

```
##     ID        Region.x         Genotype Latitude Radiation PC1_growS PC2_growS
## 1  991 Northern Europe  Ale-Stenar-41-1  55.3833  14894.71    -5.100    15.183
## 2  992 Northern Europe  Ale-Stenar-44-4  55.3833  14894.71    -5.100    15.183
## 3  997 Northern Europe Ale-Stenar-56-14  55.3833  14894.71    -5.100    15.183
## 4 1002 Northern Europe Ale-Stenar-64-24  55.3833  14894.71    -5.100    15.183
## 5 1061 Northern Europe   Brösarp-11-135  55.7167  14646.71    -0.415    13.372
## 6 1062 Northern Europe   Brösarp-15-138  55.7167  14646.71    -0.415    13.372
##    PC1_T  PC2_T    PC1_P  PC2_P             Name        Region.y Country
## 1 -1.345  8.939 -140.029 -3.076  Ale-Stenar-41-1 Northern Europe  Sweden
## 2 -1.345  8.939 -140.029 -3.076  Ale-Stenar-44-4 Northern Europe  Sweden
## 3 -1.345  8.939 -140.029 -3.076 Ale-Stenar-56-14 Northern Europe  Sweden
## 4 -1.345  8.939 -140.029 -3.076 Ale-Stenar-64-24 Northern Europe  Sweden
## 5 -9.538 14.299  -97.426 -8.890   Brösarp-11-135 Northern Europe  Sweden
## 6 -9.538 14.299  -97.426 -8.890   Brösarp-15-138 Northern Europe  Sweden
##   Final_Size_HL  T50_HL Slope_HL Final_Size_LL  T50_LL Slope_LL GXE_Final_Size
## 1        8.6731 10.5321   5.7169       14.5374 34.5172   5.1963        -0.7409
## 2        7.2023 11.8398   4.8478       13.7532 33.5521   5.5908        -0.0543
## 3        6.6369 15.5743   4.0614       15.2798 32.4278   5.6417         2.0377
## 4        7.6601 13.7759   3.7600       14.2653 37.2868   4.8761        -1.1611
## 5        7.1276 15.5622   4.9051       11.4213 35.4196   5.7678        -2.3115
## 6        5.8695 16.4978   3.7495       11.0332 31.6869   5.6354        -1.4415
##   GXE_t50 GXE_Slope Phenotype_analysis GWAS DiamFieldM2 Hypocotyllength
## 1  0.4742   -1.6368                  1    1          NA              NA
## 2 -1.7986   -0.3731                  0    1          NA              NA
## 3 -6.6575    0.4641                  0    1   14.375000               5
## 4 -0.5669    0.9047                  0    1    8.069833              10
## 5 -3.6535   -0.2535                  1    1    9.499000               7
## 6 -8.3219    0.7698                  0    1   10.855333               4
##   Biomass21d  FT_10  FT_16 FT_HL FT_LL
## 1         NA 110.50 106.75    NA    NA
## 2         NA 116.00 100.00    NA    NA
## 3     0.0343 101.25  87.50    NA  81.0
## 4     0.0239 102.75  86.25    NA    NA
## 5     0.0218  97.75  94.75    NA    NA
## 6     0.0301  94.00  68.25    NA  88.5
```

```
##replace chinese names with numbers:

S19[which(S19$Region.y=="China"),"ID"] <- c("6414606","2189710","2189718","2204175","2204176","2204177","6414609","2204317","2204239","2204174",
"2204313","2204161","2204162","6414604","6414610","2204626","2204179",NA,NA,NA,"2204168","2204170")

S19$ID <- as.numeric(as.character(S19$ID))
```

```
## Warning: NAs introduced by coercion
```

```
S19 <- S19[-which(is.na(S19$ID)==T),]
S19$ID <- as.factor(S19$ID)


#western european IDs do not merge, but they will be removed now anyway:
S19 <- S19[-which(S19$Region.x=="Western Europe"),]
S19 <- S19[-which(S19$Region.y=="Western Europe"),]
#only phenotype_analysis data:
S19 <- S19[which(S19$Phenotype_analysis==1),]
#6414606,9580 and 9829 have no genetic data for the kinship matrix:
S19 <- S19[-which(S19$ID %in% c("6414606","9580","9829")),]
#reorder the table:
S19 <- S19[,c("ID","Final_Size_HL","T50_HL","Slope_HL","FT_HL",
              "Final_Size_LL","T50_LL","Slope_LL","FT_LL",
              "GXE_Final_Size","GXE_t50","GXE_Slope","Latitude","FT_10","FT_16",
              "Radiation","PC1_growS","PC2_growS","PC1_T","PC2_T","PC1_P","PC2_P")]
#rename variables to have shorter names for the plot:
colnames(S19) <- c("ID","FS_HL","T50_HL","SL_HL","FT_HL",
              "FS_LL","T50_LL","SL_LL","FT_LL",
              "GXE_FS","GXE_T50","GXE_SL","Latitude","FT_10","FT_16",
              "Radiation","PC1_growS","PC2_growS","PC1_T","PC2_T","PC1_P","PC2_P")

####run the correlation
#estimate pearson correlation
correlation_lmekin <- cor(S19[,2:22],use = "pairwise.complete.obs",method="pearson")


pvalues_lmekin <- matrix(data = NA,nrow = 21,ncol=21)

####lmekin for p-values
require(nlme)
library(coxme)

#Kinship matrix
load("K_227_filtered.R")

for (i in 2:ncol(S19)){
  for (j in i:ncol(S19)){
    traitcorr <- lmekin(S19[,i]~S19[,j]+ (1|S19$ID) ,method="ML",varlist=K_227_filtered,na.action = na.omit)
    #extract the correlation and pvalue:
    tabli<-capture.output(print(traitcorr)[0])#extract model output
    g<-strsplit (tabli[10], "\\s+")
    pval<-g[[1]][length(g[[1]])]
    pvalues_lmekin[i-1,j-1]<-pval
    pvalues_lmekin[i-1,i-1]<-0
  }
}

rownames(correlation_lmekin) <- colnames(S19)[2:22]
colnames(correlation_lmekin) <- colnames(S19)[2:22]
correlation_lmekin
```

```
##                  FS_HL      T50_HL         SL_HL       FT_HL         FS_LL
## FS_HL      1.000000000 -0.24305247 -0.0082247514 -0.24141157  0.3893033123
## T50_HL    -0.243052465  1.00000000  0.0493839791  0.10351933 -0.1100214915
## SL_HL     -0.008224751  0.04938398  1.0000000000  0.33627007  0.1834012683
## FT_HL     -0.241411573  0.10351933  0.3362700730  1.00000000  0.2600792069
## FS_LL      0.389303312 -0.11002149  0.1834012683  0.26007921  1.0000000000
## T50_LL     0.006059101  0.18705538  0.1583256250  0.20930054  0.4312357239
## SL_LL     -0.107083766 -0.04446081  0.0644758310  0.06307283 -0.2970090597
## FT_LL     -0.143984302  0.05655674  0.3651119868  0.63145181  0.3239595376
## GXE_FS    -0.273765985  0.05201202  0.1950975921  0.42168791  0.7793418488
## GXE_T50    0.194910628 -0.62643725  0.0838651819  0.11116155  0.4170136237
## GXE_SL    -0.054155991 -0.07140392 -0.8367275967 -0.24921419 -0.3165258033
## Latitude  -0.353444982 -0.22129204 -0.1124763982 -0.29416508 -0.3806196673
## FT_10     -0.189547969 -0.24252050 -0.2268272007  0.42686384 -0.0946786619
## FT_16     -0.176148412 -0.21505018 -0.1712515036  0.69664426 -0.0558340692
## Radiation  0.237027610 -0.01932937 -0.0153353780  0.21020490  0.2356549684
## PC1_growS  0.054734028 -0.04877752 -0.0004586347 -0.27686883 -0.2079739430
## PC2_growS -0.179302016 -0.15501801  0.0626666593  0.09217922 -0.1660360190
## PC1_T     -0.325601821 -0.02645998 -0.1118007682  0.27403819 -0.0004489083
## PC2_T     -0.264138855 -0.28522561 -0.0899335205 -0.32747215 -0.3791622707
## PC1_P      0.113021839 -0.03425568 -0.0119917435 -0.26611192 -0.1569839684
## PC2_P      0.217035926  0.07415934 -0.0197036564 -0.17614400  0.0222869601
##                 T50_LL        SL_LL        FT_LL      GXE_FS     GXE_T50
## FS_HL      0.006059101 -0.107083766 -0.143984302 -0.27376598  0.19491063
## T50_HL     0.187055381 -0.044460812  0.056556744  0.05201202 -0.62643725
## SL_HL      0.158325625  0.064475831  0.365111987  0.19509759  0.08386518
## FT_HL      0.209300542  0.063072827  0.631451814  0.42168791  0.11116155
## FS_LL      0.431235724 -0.297009060  0.323959538  0.77934185  0.41701362
## T50_LL     1.000000000 -0.364328605  0.315547822  0.43246203  0.64853523
## SL_LL     -0.364328605  1.000000000 -0.054945091 -0.22508933 -0.24889596
## FT_LL      0.315547822 -0.054945091  1.000000000  0.40932708  0.22489890
## GXE_FS     0.432462025 -0.225089334  0.409327080  1.00000000  0.30284625
## GXE_T50    0.648535230 -0.248895960  0.224898897  0.30284625  1.00000000
## GXE_SL    -0.334036732  0.492531172 -0.347128140 -0.29366924 -0.20972254
## Latitude  -0.298784695  0.154136560 -0.203255318 -0.15511315 -0.06941177
## FT_10     -0.118726443  0.064847935  0.462819740  0.03051476  0.09440333
## FT_16     -0.031618282  0.026724530  0.797423950  0.05997001  0.14899402
## Radiation  0.112275414 -0.002655905  0.100562667  0.08126655  0.10682409
## PC1_growS -0.267827567 -0.012540028 -0.312100236 -0.25176946 -0.16864281
## PC2_growS -0.211137651  0.167650553  0.006942101 -0.04516417 -0.05332668
## PC1_T      0.125317483 -0.012168273  0.371717955  0.22736447  0.10943873
## PC2_T     -0.330949495  0.176696210 -0.249949604 -0.20958880 -0.04342804
## PC1_P     -0.191222087 -0.037162574 -0.269166870 -0.23971082 -0.11794863
## PC2_P     -0.032758723 -0.174501887 -0.169115934 -0.12926210 -0.06674902
##                GXE_SL    Latitude       FT_10       FT_16    Radiation
## FS_HL     -0.05415599 -0.35344498 -0.18954797 -0.17614841  0.237027610
## T50_HL    -0.07140392 -0.22129204 -0.24252050 -0.21505018 -0.019329367
## SL_HL     -0.83672760 -0.11247640 -0.22682720 -0.17125150 -0.015335378
## FT_HL     -0.24921419 -0.29416508  0.42686384  0.69664426  0.210204898
## FS_LL     -0.31652580 -0.38061967 -0.09467866 -0.05583407  0.235654968
## T50_LL    -0.33403673 -0.29878470 -0.11872644 -0.03161828  0.112275414
## SL_LL      0.49253117  0.15413656  0.06484793  0.02672453 -0.002655905
## FT_LL     -0.34712814 -0.20325532  0.46281974  0.79742395  0.100562667
## GXE_FS    -0.29366924 -0.15511315  0.03051476  0.05997001  0.081266545
## GXE_T50   -0.20972254 -0.06941177  0.09440333  0.14899402  0.106824092
## GXE_SL     1.00000000  0.18690292  0.24139713  0.17297969  0.011030133
## Latitude   0.18690292  1.00000000  0.15508822  0.08985714 -0.582680027
## FT_10      0.24139713  0.15508822  1.00000000  0.89531244  0.319160194
## FT_16      0.17297969  0.08985714  0.89531244  1.00000000  0.266596467
## Radiation  0.01103013 -0.58268003  0.31916019  0.26659647  1.000000000
## PC1_growS -0.01490273  0.24874029 -0.11424635 -0.14717494 -0.100481941
## PC2_growS  0.03392057  0.65747895  0.10776665  0.08744603 -0.337610900
## PC1_T      0.10214629  0.25975952  0.09176120  0.07357720 -0.217359511
## PC2_T      0.17967034  0.86733434  0.29820614  0.22610430 -0.209778476
## PC1_P     -0.01767987  0.06075398 -0.09227627 -0.12289726  0.058247998
## PC2_P     -0.08975534 -0.32808690 -0.12523651 -0.12297609  0.204252920
##               PC1_growS    PC2_growS         PC1_T       PC2_T       PC1_P
## FS_HL      0.0547340280 -0.179302016 -0.3256018208 -0.26413885  0.11302184
## T50_HL    -0.0487775236 -0.155018015 -0.0264599805 -0.28522561 -0.03425568
## SL_HL     -0.0004586347  0.062666659 -0.1118007682 -0.08993352 -0.01199174
## FT_HL     -0.2768688277  0.092179224  0.2740381857 -0.32747215 -0.26611192
## FS_LL     -0.2079739430 -0.166036019 -0.0004489083 -0.37916227 -0.15698397
## T50_LL    -0.2678275674 -0.211137651  0.1253174830 -0.33094949 -0.19122209
## SL_LL     -0.0125400282  0.167650553 -0.0121682730  0.17669621 -0.03716257
## FT_LL     -0.3121002358  0.006942101  0.3717179546 -0.24994960 -0.26916687
## GXE_FS    -0.2517694615 -0.045164172  0.2273644672 -0.20958880 -0.23971082
## GXE_T50   -0.1686428057 -0.053326683  0.1094387281 -0.04342804 -0.11794863
## GXE_SL    -0.0149027267  0.033920571  0.1021462891  0.17967034 -0.01767987
## Latitude   0.2487402905  0.657478948  0.2597595244  0.86733434  0.06075398
## FT_10     -0.1142463507  0.107766652  0.0917612039  0.29820614 -0.09227627
## FT_16     -0.1471749414  0.087446030  0.0735771990  0.22610430 -0.12289726
## Radiation -0.1004819409 -0.337610900 -0.2173595111 -0.20977848  0.05824800
## PC1_growS  1.0000000000  0.191552353 -0.6398079517  0.38805682  0.94477835
## PC2_growS  0.1915523534  1.000000000  0.0385398867  0.65728593 -0.07742936
## PC1_T     -0.6398079517  0.038539887  1.0000000000  0.05131258 -0.70350060
## PC2_T      0.3880568218  0.657285925  0.0513125801  1.00000000  0.22370379
## PC1_P      0.9447783526 -0.077429361 -0.7035006034  0.22370379  1.00000000
## PC2_P      0.6004615038 -0.544276267 -0.6409223969 -0.23491685  0.79347138
##                 PC2_P
## FS_HL      0.21703593
## T50_HL     0.07415934
## SL_HL     -0.01970366
## FT_HL     -0.17614400
## FS_LL      0.02228696
## T50_LL    -0.03275872
## SL_LL     -0.17450189
## FT_LL     -0.16911593
## GXE_FS    -0.12926210
## GXE_T50   -0.06674902
## GXE_SL    -0.08975534
## Latitude  -0.32808690
## FT_10     -0.12523651
## FT_16     -0.12297609
## Radiation  0.20425292
## PC1_growS  0.60046150
## PC2_growS -0.54427627
## PC1_T     -0.64092240
## PC2_T     -0.23491685
## PC1_P      0.79347138
## PC2_P      1.00000000
```

```
pvalues_lmekin
```

```
##       [,1] [,2]      [,3]      [,4]      [,5]      [,6]      [,7]     
##  [1,] "0"  "4.6e-06" "3.6e-01" "0.048"   "8.9e-06" "3.0e-01" "2.6e-01"
##  [2,] NA   "0"       "5.6e-01" "0.44000" "1.3e-02" "0.00940" "9.1e-01"
##  [3,] NA   NA        "0"       "0.039"   "0.072"   "0.13"    "0.320"  
##  [4,] NA   NA        NA        "0"       "0.0460"  "0.85000" "1.2e-01"
##  [5,] NA   NA        NA        NA        "0"       "1.0e-07" "4.2e-05"
##  [6,] NA   NA        NA        NA        NA        "0"       "2.1e-06"
##  [7,] NA   NA        NA        NA        NA        NA        "0"      
##  [8,] NA   NA        NA        NA        NA        NA        NA       
##  [9,] NA   NA        NA        NA        NA        NA        NA       
## [10,] NA   NA        NA        NA        NA        NA        NA       
## [11,] NA   NA        NA        NA        NA        NA        NA       
## [12,] NA   NA        NA        NA        NA        NA        NA       
## [13,] NA   NA        NA        NA        NA        NA        NA       
## [14,] NA   NA        NA        NA        NA        NA        NA       
## [15,] NA   NA        NA        NA        NA        NA        NA       
## [16,] NA   NA        NA        NA        NA        NA        NA       
## [17,] NA   NA        NA        NA        NA        NA        NA       
## [18,] NA   NA        NA        NA        NA        NA        NA       
## [19,] NA   NA        NA        NA        NA        NA        NA       
## [20,] NA   NA        NA        NA        NA        NA        NA       
## [21,] NA   NA        NA        NA        NA        NA        NA       
##       [,8]      [,9]      [,10]     [,11]     [,12]     [,13]     [,14]    
##  [1,] "1.4e-02" "2.2e-07" "1.1e-02" "8.9e-01" "6.2e-07" "1.0e-01" "9.5e-02"
##  [2,] "6.1e-01" "6.9e-01" "0.0e+00" "5.9e-01" "0.0028"  "0.00098" "0.0046" 
##  [3,] "0.850"   "0.0180"  "0.450"   "0"       "0.14"    "0.0021"  "0.025"  
##  [4,] "7.3e-09" "1.2e-03" "5.9e-01" "4.9e-01" "0.028"   "7.8e-09" "8.2e-13"
##  [5,] "0.0045"  "0"       "4.0e-09" "1.4e-04" "6.7e-08" "8.5e-01" "8.9e-01"
##  [6,] "7.4e-01" "5.4e-09" "0"       "6.2e-05" "4e-05"   "0.26"    "0.8"    
##  [7,] "0.65"    "0.0016"  "0.00046" "1.2e-14" "0.041"   "0.39"    "0.73"   
##  [8,] "0"       "9.2e-04" "4.0e-01" "6.9e-01" "0.14"    "2.6e-05" "0.0e+00"
##  [9,] NA        "0"       "1.5e-05" "2.8e-05" "0.039"   "0.69"    "0.44"   
## [10,] NA        NA        "0"       "0.0034"  "0.36"    "0.2100"  "5.4e-02"
## [11,] NA        NA        NA        "0"       "0.0130"  "1.1e-03" "0.02500"
## [12,] NA        NA        NA        NA        "0"       "0.33"    "0.36"   
## [13,] NA        NA        NA        NA        NA        "0"       "0.0e+00"
## [14,] NA        NA        NA        NA        NA        NA        "0"      
## [15,] NA        NA        NA        NA        NA        NA        NA       
## [16,] NA        NA        NA        NA        NA        NA        NA       
## [17,] NA        NA        NA        NA        NA        NA        NA       
## [18,] NA        NA        NA        NA        NA        NA        NA       
## [19,] NA        NA        NA        NA        NA        NA        NA       
## [20,] NA        NA        NA        NA        NA        NA        NA       
## [21,] NA        NA        NA        NA        NA        NA        NA       
##       [,15]     [,16]     [,17]     [,18]     [,19]     [,20]     [,21]    
##  [1,] "0.09900" "4.6e-01" "9.4e-01" "9.0e-04" "4.7e-01" "4.3e-01" "2.1e-01"
##  [2,] "2.7e-01" "7.1e-01" "3.1e-01" "5.5e-01" "0.00012" "8.5e-01" "4.6e-01"
##  [3,] "8.4e-01" "1"       "0.42"    "0.15"    "0.24"    "0.88"    "0.8"    
##  [4,] "0.3800"  "2.5e-01" "4.1e-01" "1.2e-01" "0.014"   "3.3e-01" "5.9e-01"
##  [5,] "9.1e-02" "5.1e-02" "8.6e-01" "3e-01"   "1.4e-07" "7.5e-02" "6.0e-01"
##  [6,] "6.7e-01" "0.00036" "0.093"   "0.074"   "6.6e-06" "0.081"   "0.58"   
##  [7,] "0.97"    "0.87"    "0.029"   "0.88"    "0.021"   "0.63"    "0.023"  
##  [8,] "4.8e-01" "0.021"   "0.96"    "0.0051"  "0.071"   "0.05"    "0.23"   
##  [9,] "0.29"    "8.3e-04" "5.6e-01" "2.7e-03" "5.9e-03" "1.5e-03" "9.4e-02"
## [10,] "0.170"   "0.028"   "0.49"    "0.16"    "0.58"    "0.13"    "0.39"   
## [11,] "0.89"    "8.5e-01" "6.6e-01" "1.9e-01" "1.9e-02" "8.2e-01" "2.5e-01"
## [12,] "7.8e-05" "0.022"   "7.2e-07" "0.035"   "0"       "0.27"    "0.043"  
## [13,] "9.6e-05" "0.10000" "0.27000" "0.50000" "0.01500" "0.28000" "0.34000"
## [14,] "0.019"   "0.041"   "0.510"   "0.530"   "0.022"   "0.170"   "0.34"   
## [15,] "0"       "8.8e-01" "7.8e-01" "1.2e-01" "6.0e-06" "1.9e-01" "2.0e-01"
## [16,] NA        "0"       "0.47"    "0.00"    "0.00001" "0.00"    "0.00"   
## [17,] NA        NA        "0"       "0.23"    "3.4e-06" "0.0037"  "1.3e-09"
## [18,] NA        NA        NA        "0"       "0.044"   "0.00"    "0.00"   
## [19,] NA        NA        NA        NA        "0"       "0.00019" "0.93"   
## [20,] NA        NA        NA        NA        NA        "0"       "0.00"   
## [21,] NA        NA        NA        NA        NA        NA        "0"
```

```
corr <- apply(correlation_lmekin,FUN=as.numeric,MARGIN = 2)
rownames(corr) <- colnames(S19)[2:22]
pval <- apply(pvalues_lmekin,FUN=as.numeric,MARGIN = 2)
pval.adj <- matrix(p.adjust(t(pval),method = "fdr"),nrow=21,ncol=21,byrow=T)#correct for multiple testing
```

###### Fig S19

```
#Plot
library(corrplot)
col1 <- colorRampPalette(c("blue4", "white", "red2"))#blue to red, but pretty versions of it

corrplot(corr=t(corr),method="square", type="lower", p.mat =t(pval.adj),sig.level=0.05, diag=F,
         insig = "blank",tl.col="black", mar = c(1,1,1,1),col=col1(200),tl.pos="ld",tl.srt=360)
```
