## Supplementary Figures for "Polygenic adaptation of rosette growth in *Arabidopsis thaliana*"

### Slide 1
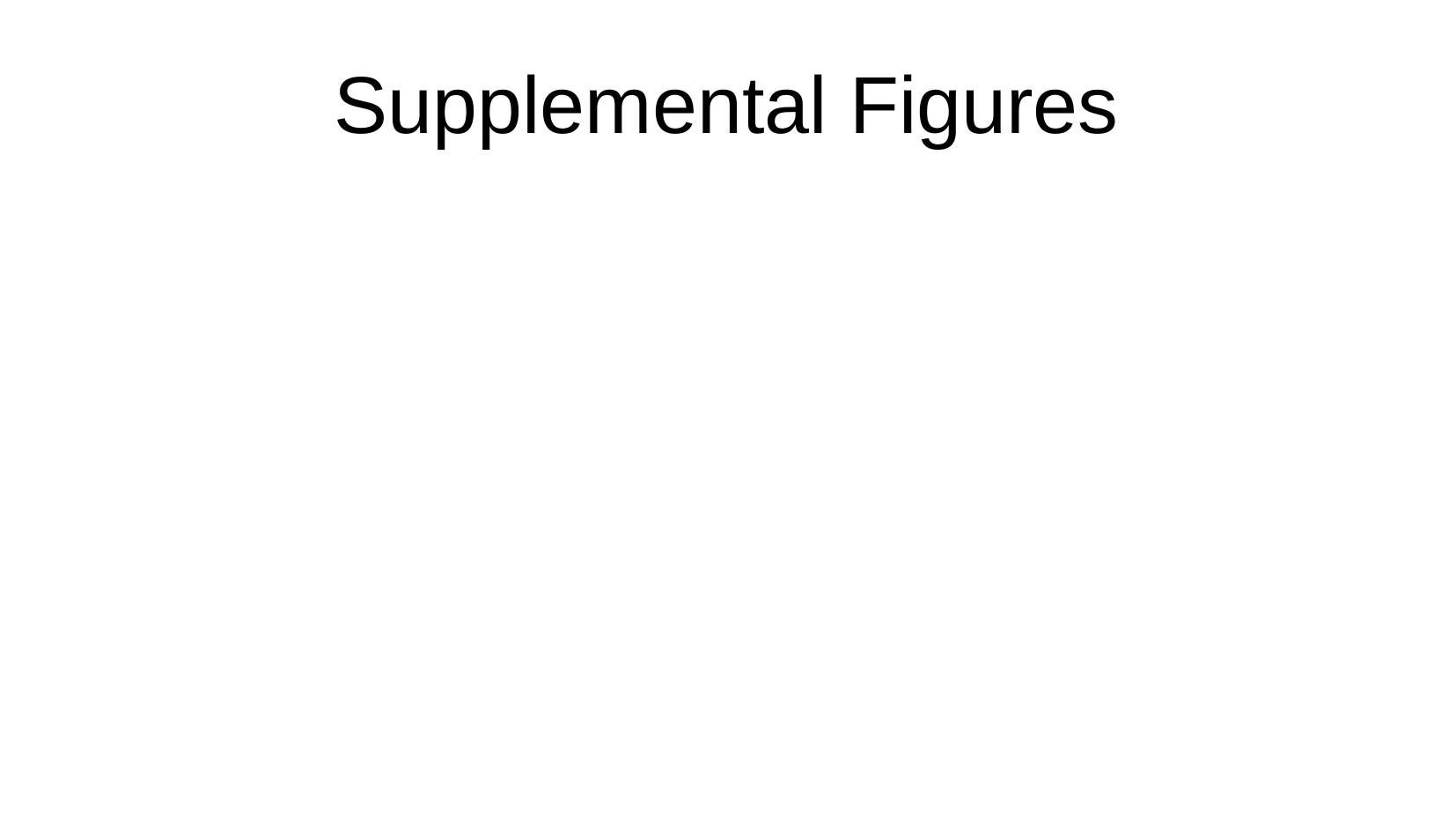

Supplemental Figures

### Slide 2
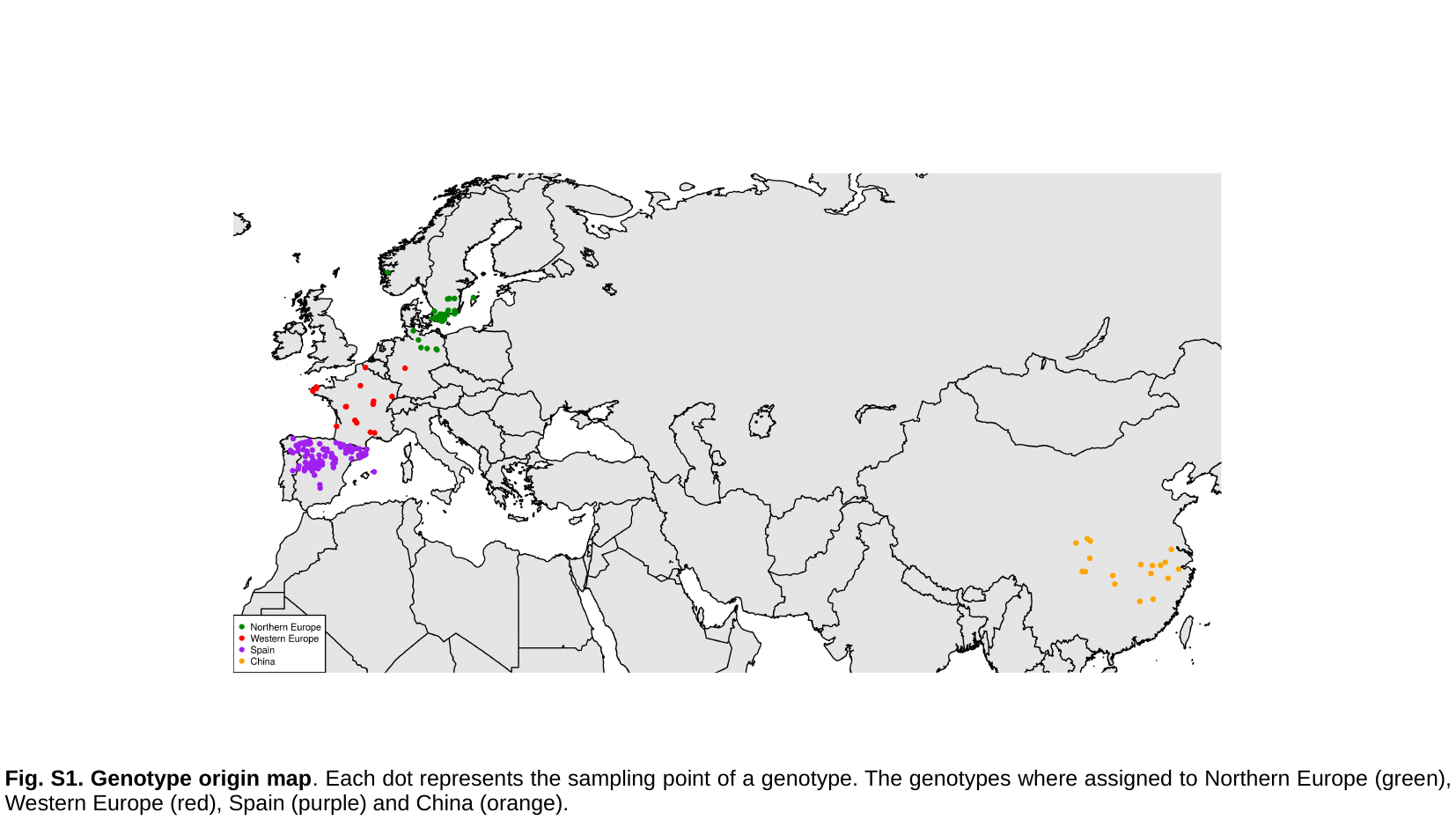

Fig. S1. Genotype origin map. Each dot represents the sampling point of a genotype. The genotypes where assigned to Northern Europe (green), Western Europe (red), Spain (purple) and China (orange).

### Slide 3
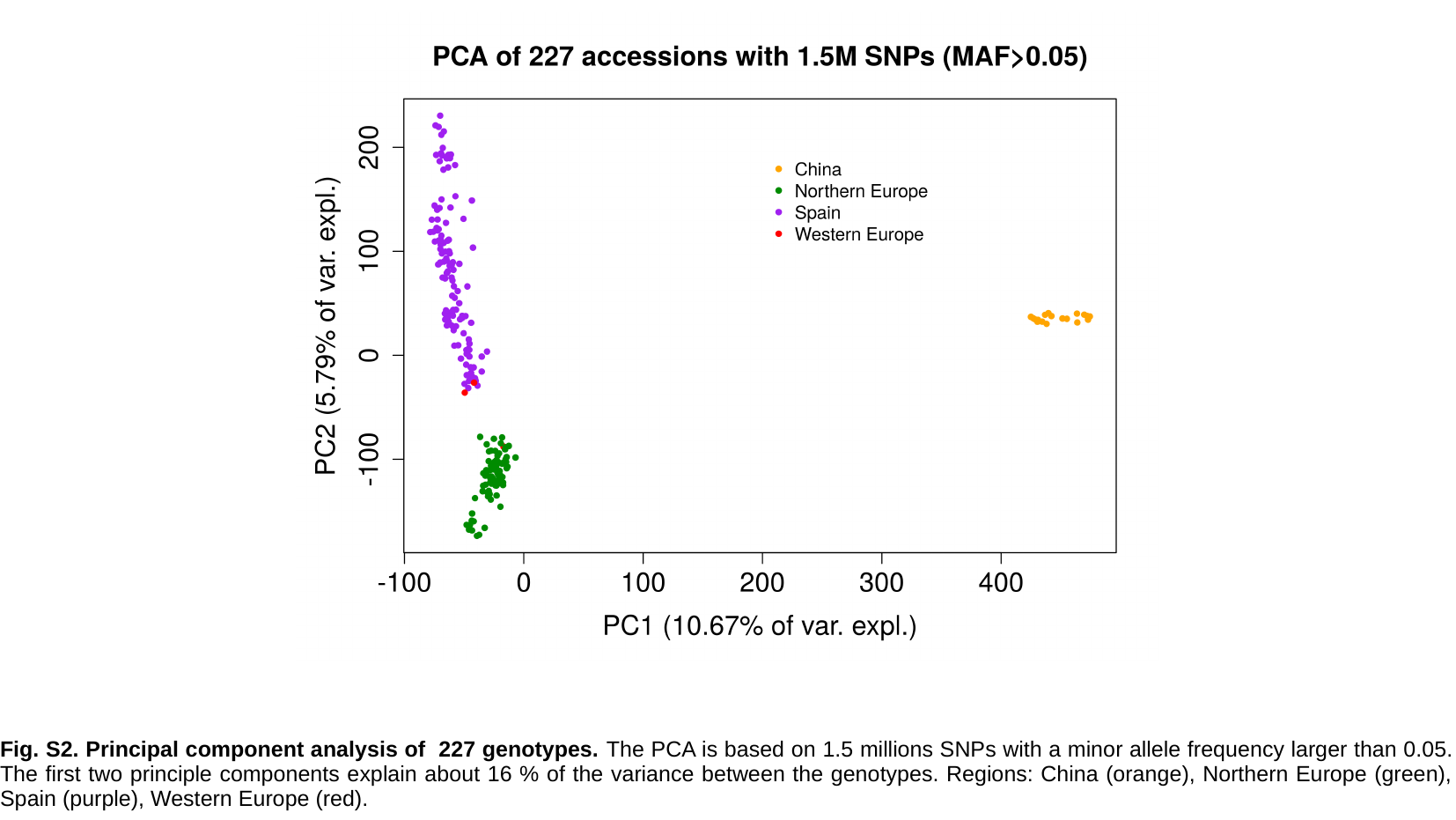

Fig. S2. Principal component analysis of 227 genotypes. The PCA is based on 1.5 millions SNPs with a minor allele frequency larger than 0.05. The first two principle components explain about 16 % of the variance between the genotypes. Regions: China (orange), Northern Europe (green), Spain (purple), Western Europe (red).

### Slide 4
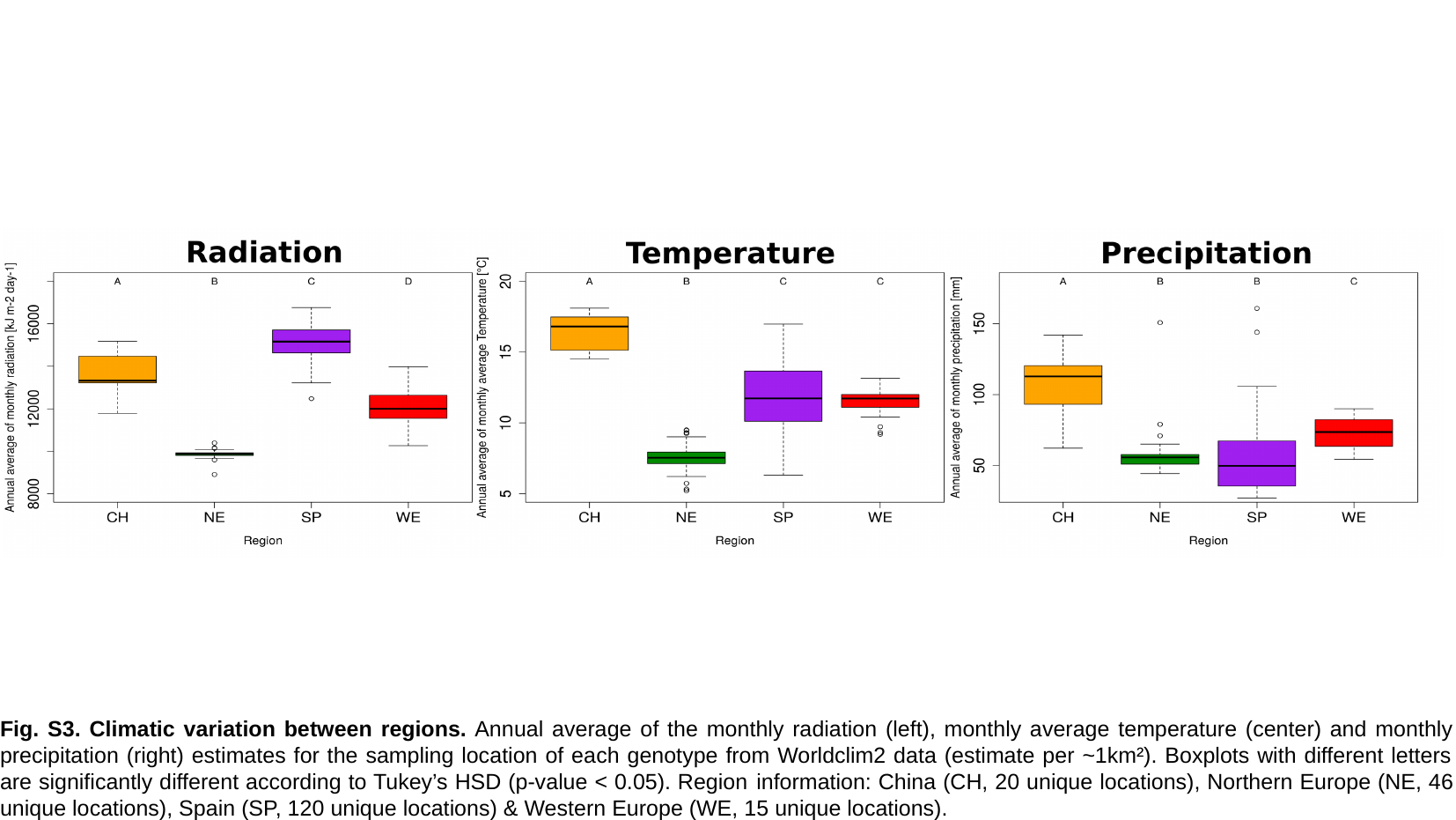

Fig. S3. Climatic variation between regions. Annual average of the monthly radiation (left), monthly average temperature (center) and monthly precipitation (right) estimates for the sampling location of each genotype from Worldclim2 data (estimate per ~1km²). Boxplots with different letters are significantly different according to Tukey’s HSD (p-value < 0.05). Region information: China (CH, 20 unique locations), Northern Europe (NE, 46 unique locations), Spain (SP, 120 unique locations) & Western Europe (WE, 15 unique locations).

### Slide 5
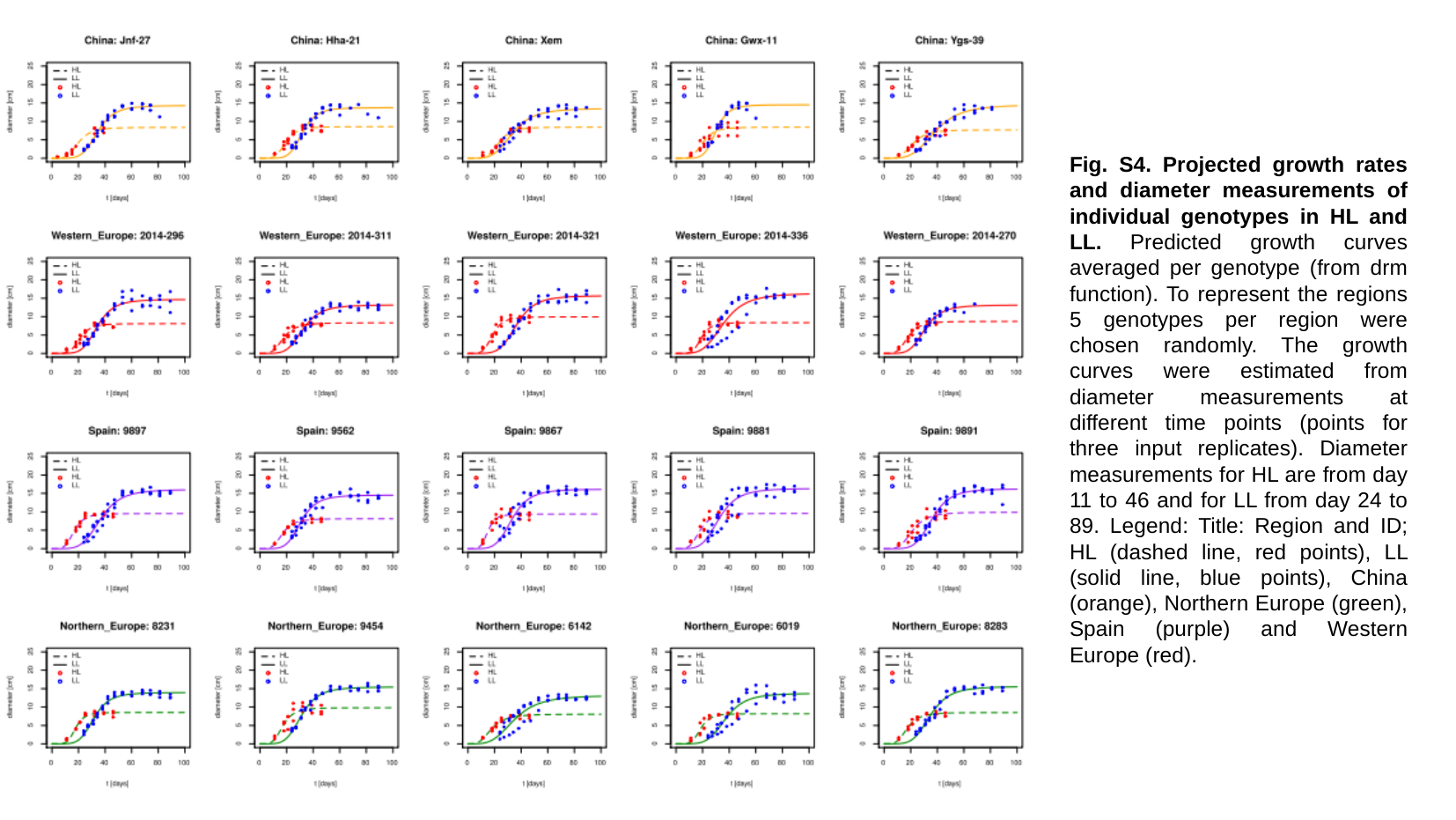

Fig. S4. Projected growth rates and diameter measurements of individual genotypes in HL and LL. Predicted growth curves averaged per genotype (from drm function). To represent the regions 5 genotypes per region were chosen randomly. The growth curves were estimated from diameter measurements at different time points (points for three input replicates). Diameter measurements for HL are from day 11 to 46 and for LL from day 24 to 89. Legend: Title: Region and ID; HL (dashed line, red points), LL (solid line, blue points), China (orange), Northern Europe (green), Spain (purple) and Western Europe (red).

### Slide 6
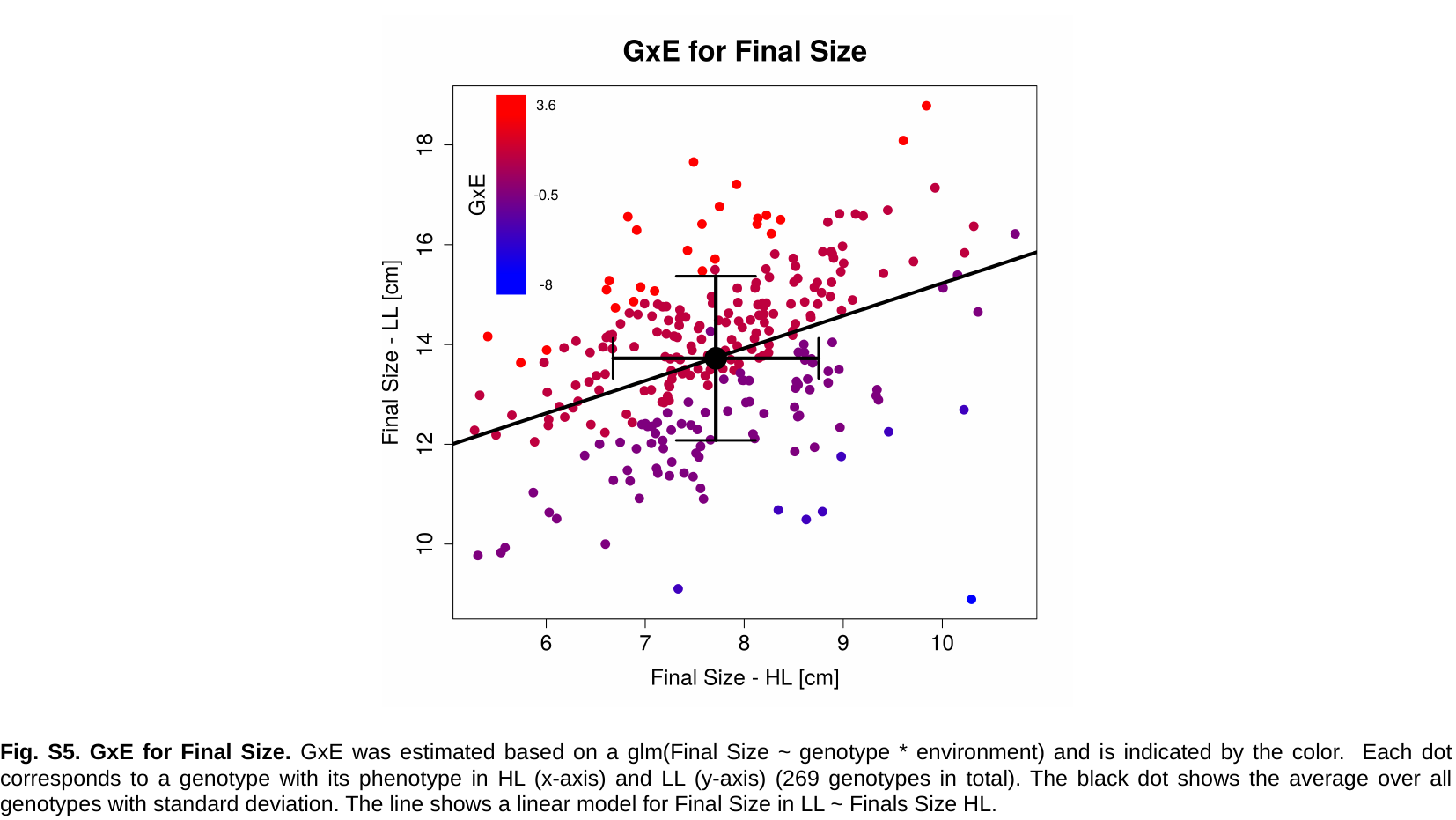

Fig. S5. GxE for Final Size. GxE was estimated based on a glm(Final Size ~ genotype * environment) and is indicated by the color. Each dot corresponds to a genotype with its phenotype in HL (x-axis) and LL (y-axis) (269 genotypes in total). The black dot shows the average over all genotypes with standard deviation. The line shows a linear model for Final Size in LL ~ Finals Size HL.

### Slide 7
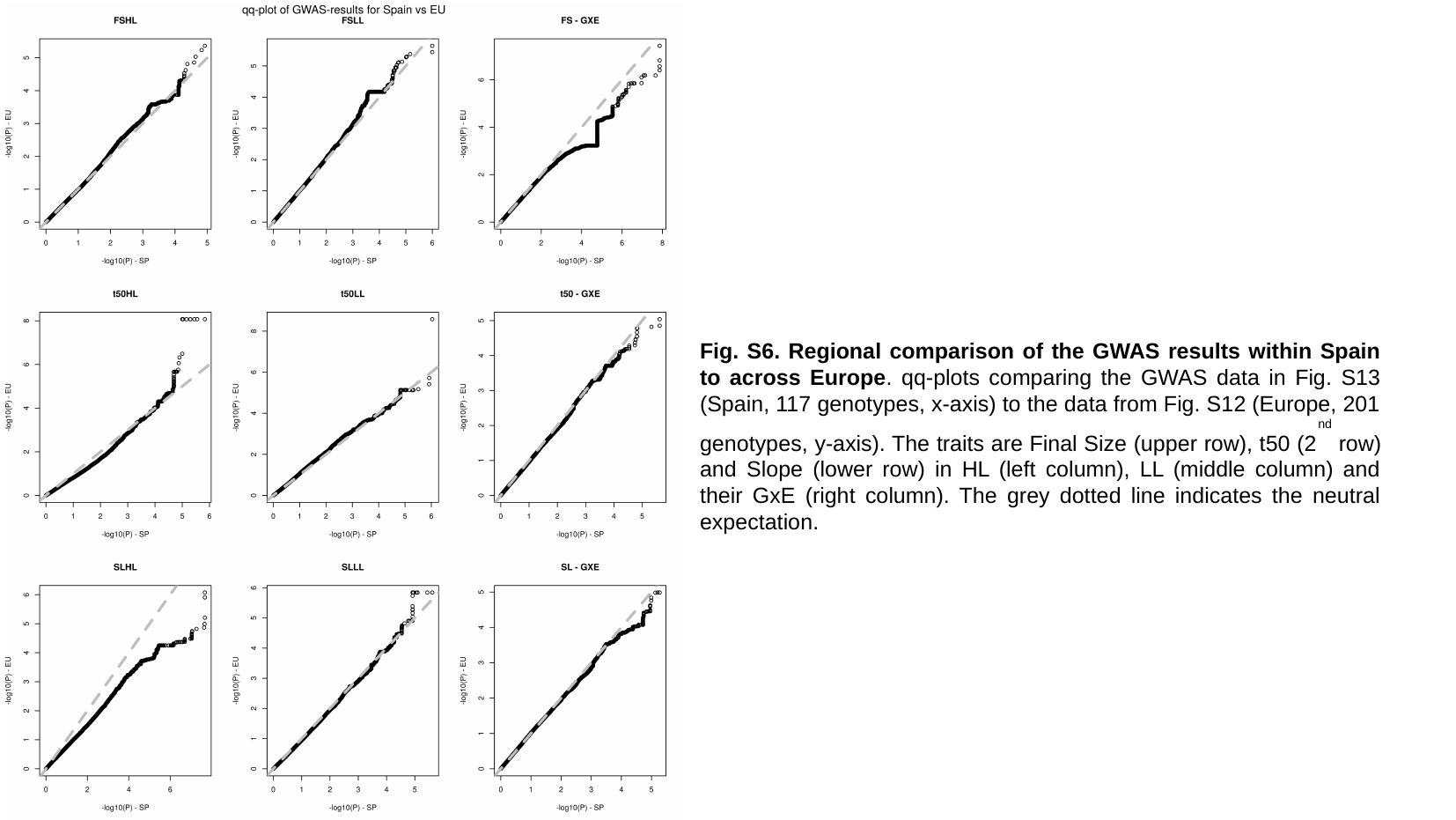

Fig. S6. Regional comparison of the GWAS results within Spain to across Europe. qq-plots comparing the GWAS data in Fig. S13 (Spain, 117 genotypes, x-axis) to the data from Fig. S12 (Europe, 201 genotypes, y-axis). The traits are Final Size (upper row), t50 (2^nd^ row) and Slope (lower row) in HL (left column), LL (middle column) and their GxE (right column). The grey dotted line indicates the neutral expectation.

### Slide 8
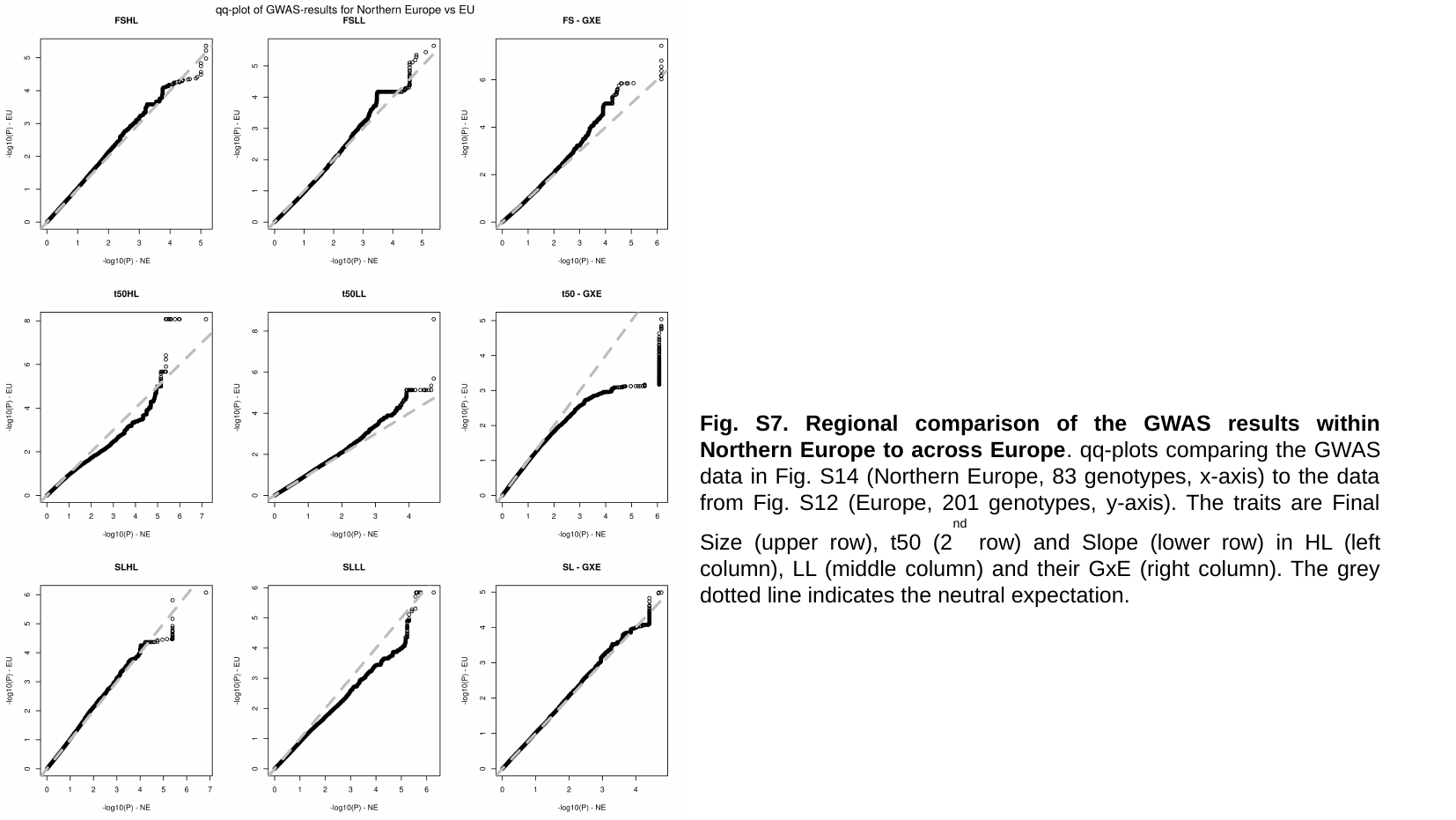

Fig. S7. Regional comparison of the GWAS results within Northern Europe to across Europe. qq-plots comparing the GWAS data in Fig. S14 (Northern Europe, 83 genotypes, x-axis) to the data from Fig. S12 (Europe, 201 genotypes, y-axis). The traits are Final Size (upper row), t50 (2^nd^ row) and Slope (lower row) in HL (left column), LL (middle column) and their GxE (right column). The grey dotted line indicates the neutral expectation.

### Slide 9
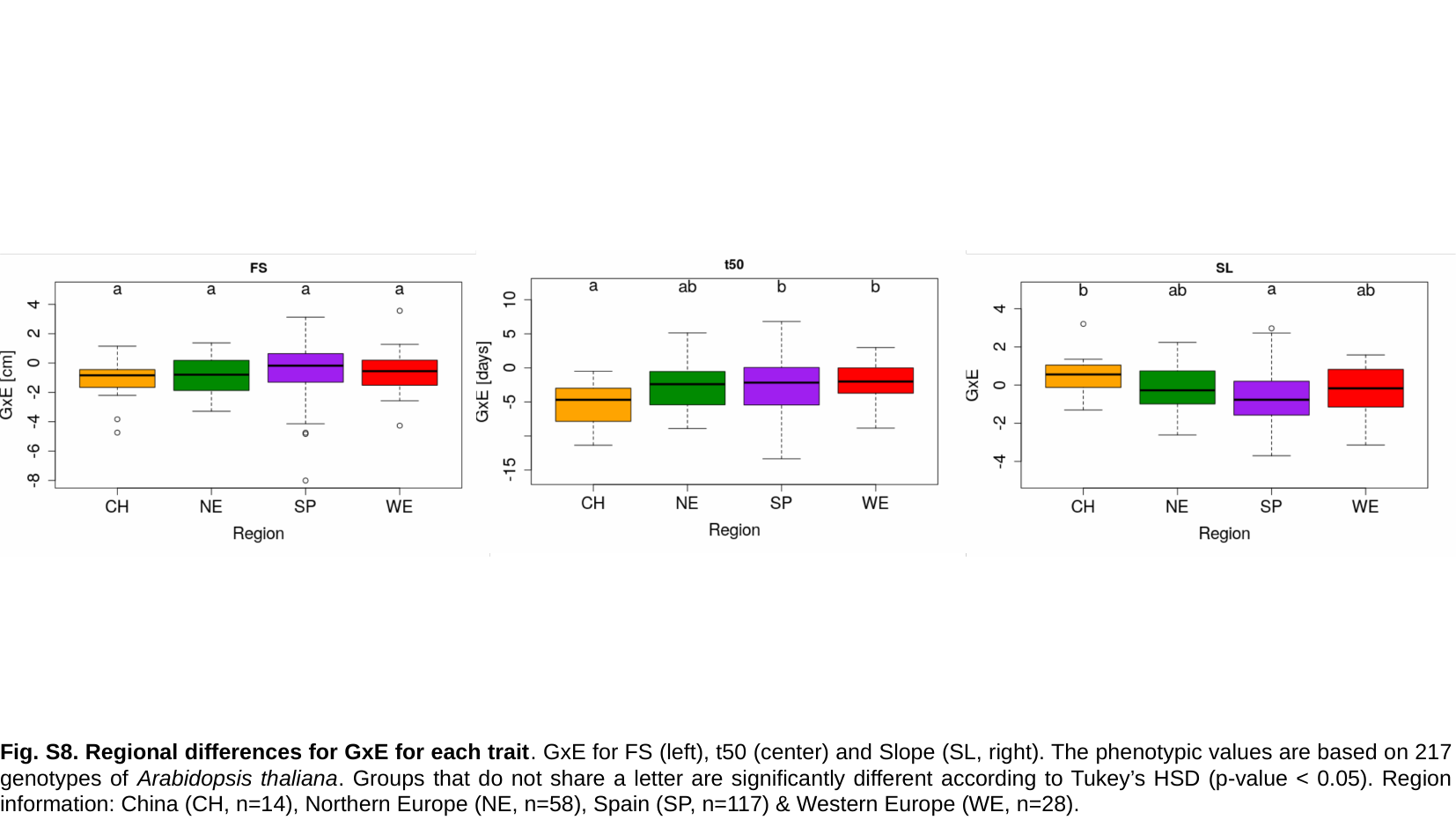

Fig. S8. Regional differences for GxE for each trait. GxE for FS (left), t50 (center) and Slope (SL, right). The phenotypic values are based on 217 genotypes of Arabidopsis thaliana. Groups that do not share a letter are significantly different according to Tukey’s HSD (p-value < 0.05). Region information: China (CH, n=14), Northern Europe (NE, n=58), Spain (SP, n=117) & Western Europe (WE, n=28).

### Slide 10
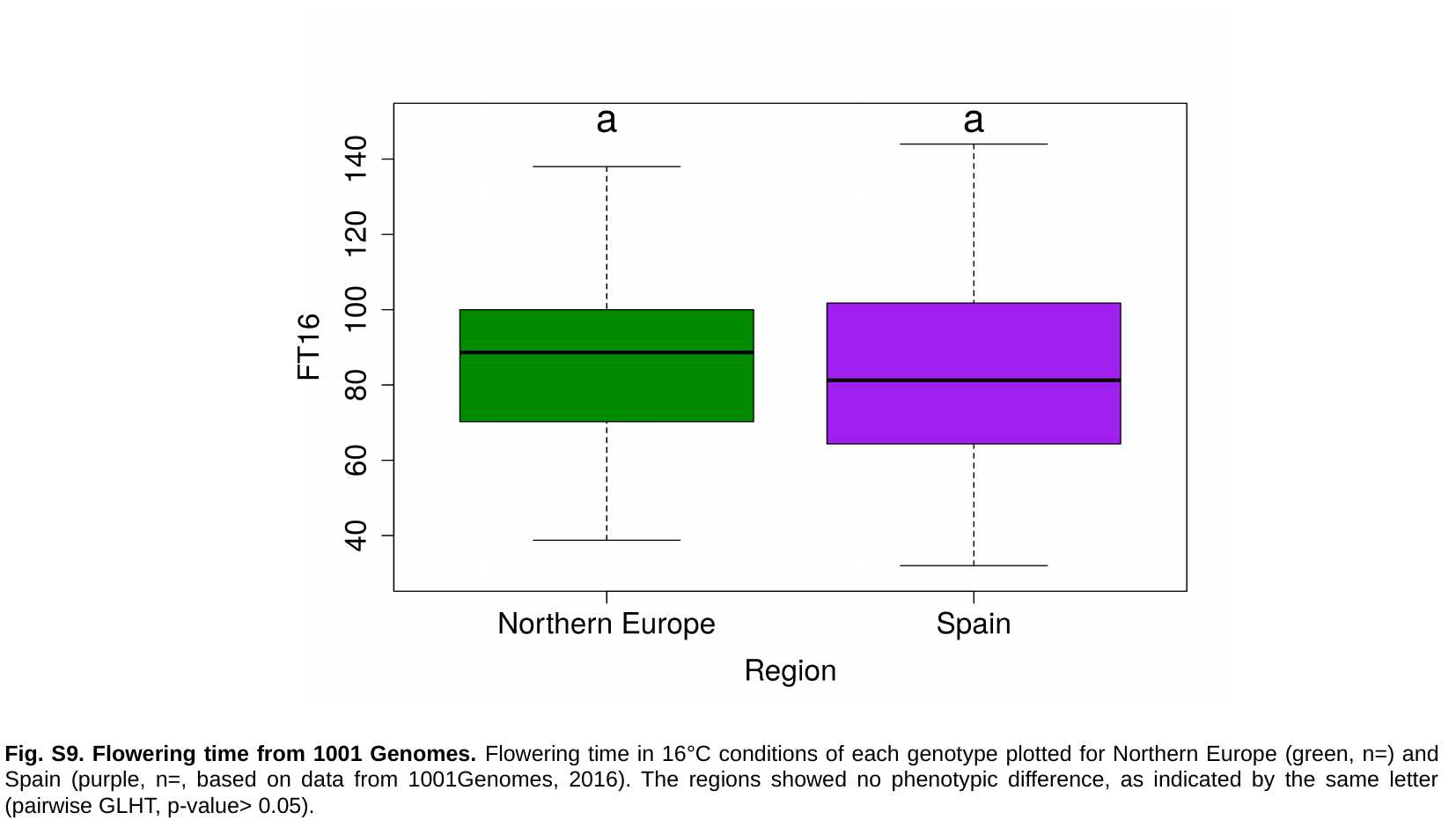

Fig. S9. Flowering time from 1001 Genomes. Flowering time in 16°C conditions of each genotype plotted for Northern Europe (green, n=) and Spain (purple, n=, based on data from 1001Genomes, 2016). The regions showed no phenotypic difference, as indicated by the same letter (pairwise GLHT, p-value> 0.05).

### Slide 11
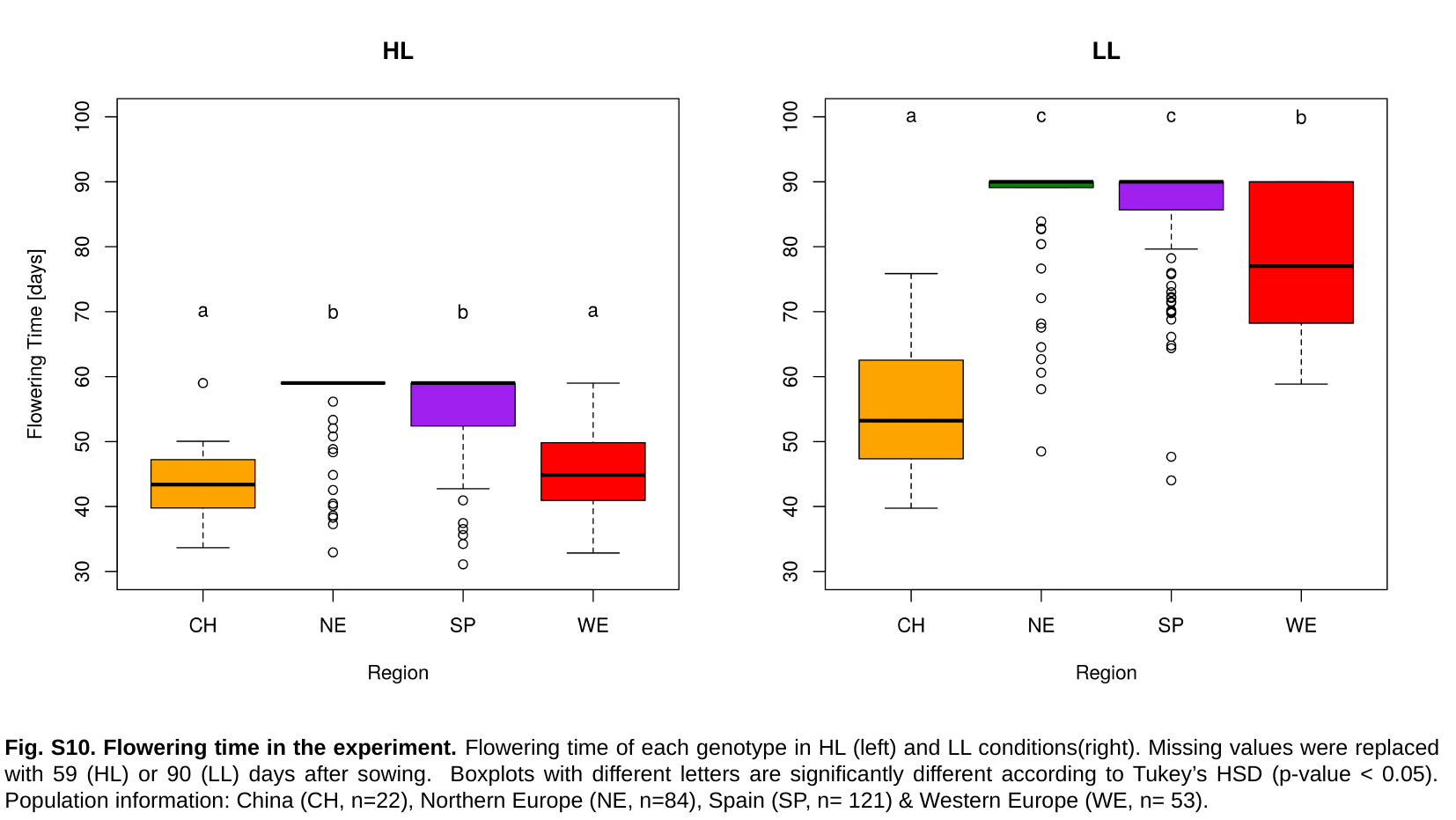

Fig. S10. Flowering time in the experiment. Flowering time of each genotype in HL (left) and LL conditions(right). Missing values were replaced with 59 (HL) or 90 (LL) days after sowing. Boxplots with different letters are significantly different according to Tukey’s HSD (p-value < 0.05). Population information: China (CH, n=22), Northern Europe (NE, n=84), Spain (SP, n= 121) & Western Europe (WE, n= 53).

### Slide 12
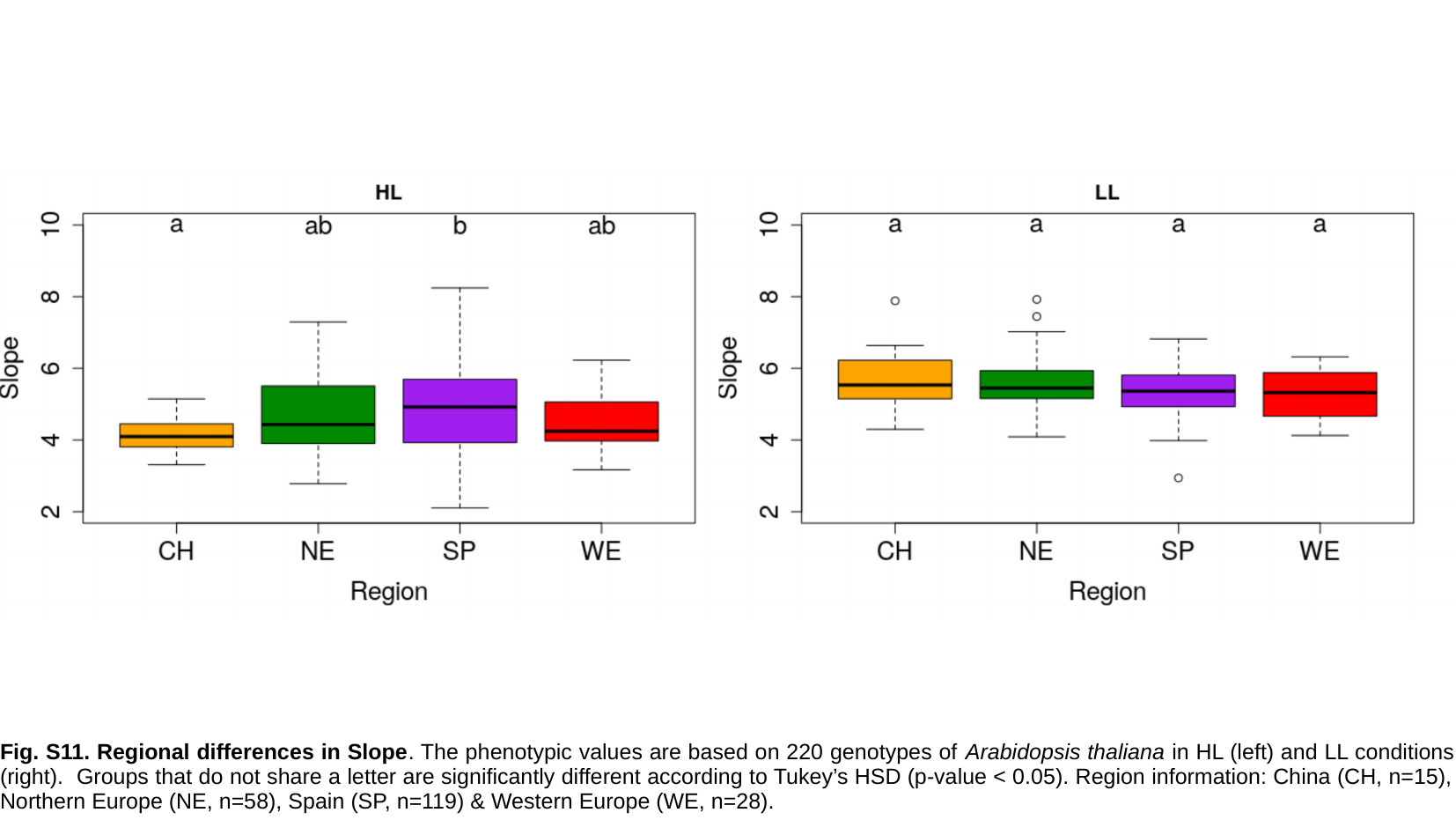

Fig. S11. Regional differences in Slope. The phenotypic values are based on 220 genotypes of Arabidopsis thaliana in HL (left) and LL conditions (right). Groups that do not share a letter are significantly different according to Tukey’s HSD (p-value < 0.05). Region information: China (CH, n=15), Northern Europe (NE, n=58), Spain (SP, n=119) & Western Europe (WE, n=28).

### Slide 13
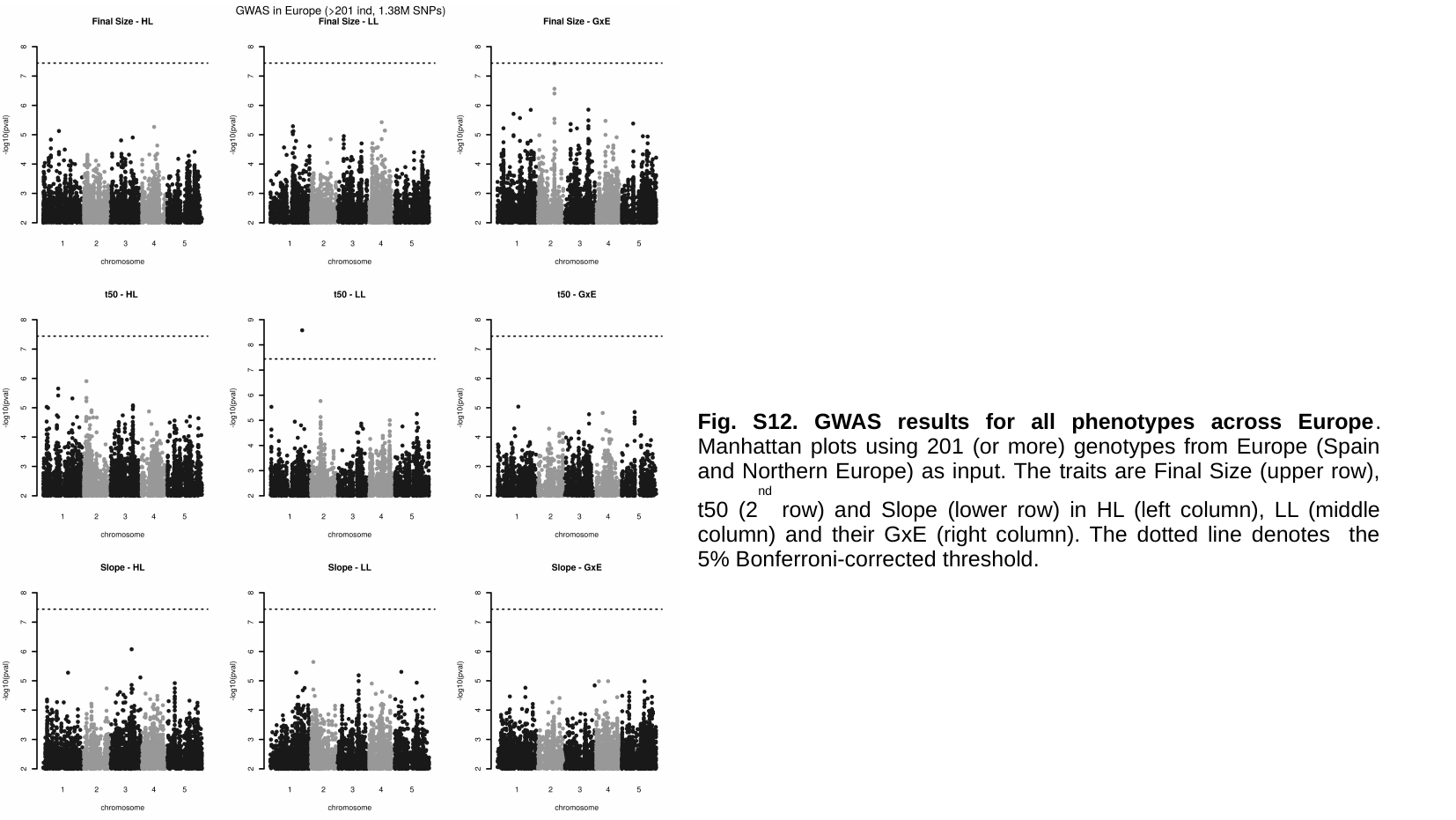

Fig. S12. GWAS results for all phenotypes across Europe. Manhattan plots using 201 (or more) genotypes from Europe (Spain and Northern Europe) as input. The traits are Final Size (upper row), t50 (2^nd^ row) and Slope (lower row) in HL (left column), LL (middle column) and their GxE (right column). The dotted line denotes the 5% Bonferroni-corrected threshold.

### Slide 14
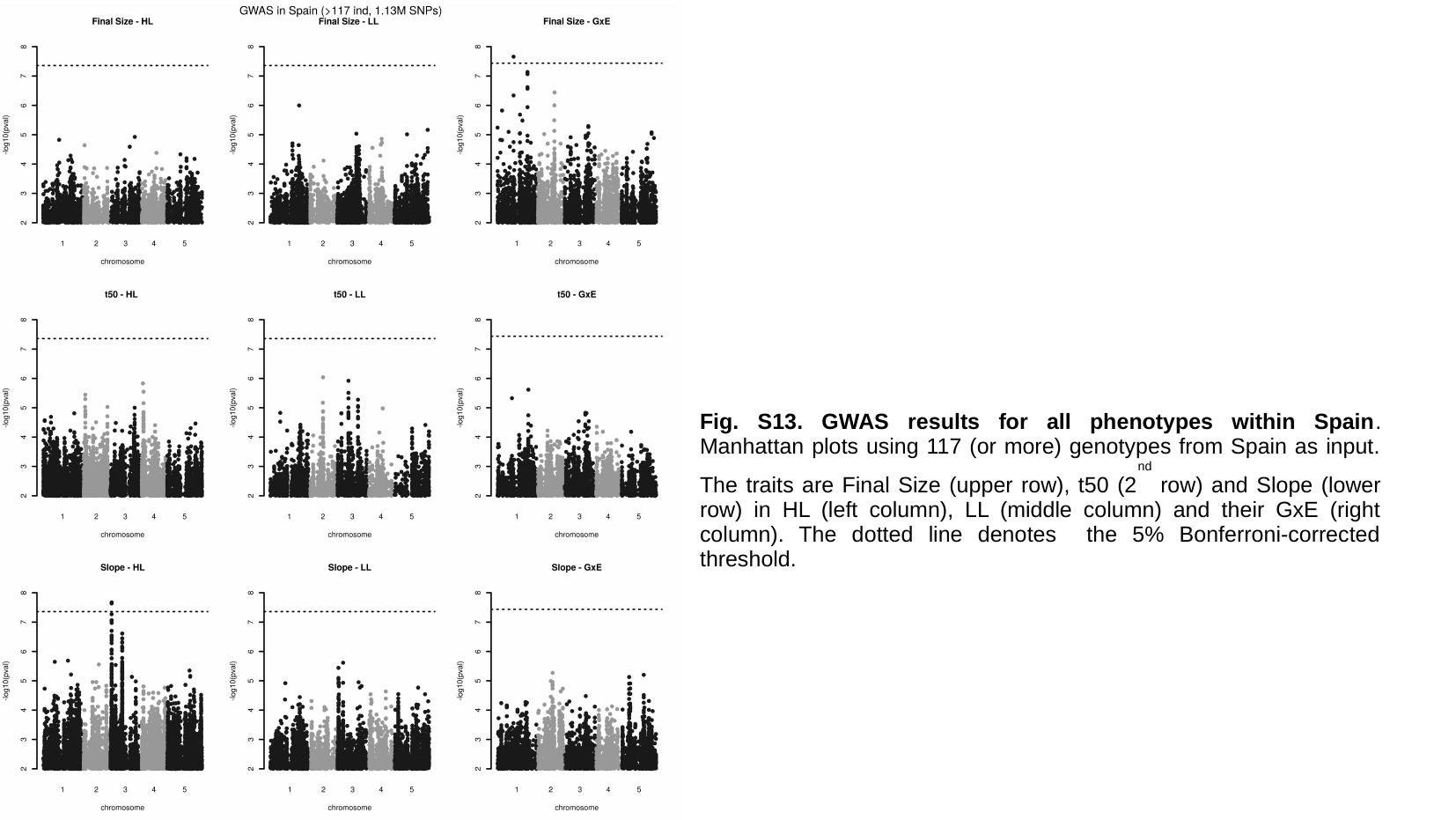

Fig. S13. GWAS results for all phenotypes within Spain. Manhattan plots using 117 (or more) genotypes from Spain as input. The traits are Final Size (upper row), t50 (2^nd^ row) and Slope (lower row) in HL (left column), LL (middle column) and their GxE (right column). The dotted line denotes the 5% Bonferroni-corrected threshold.

### Slide 15
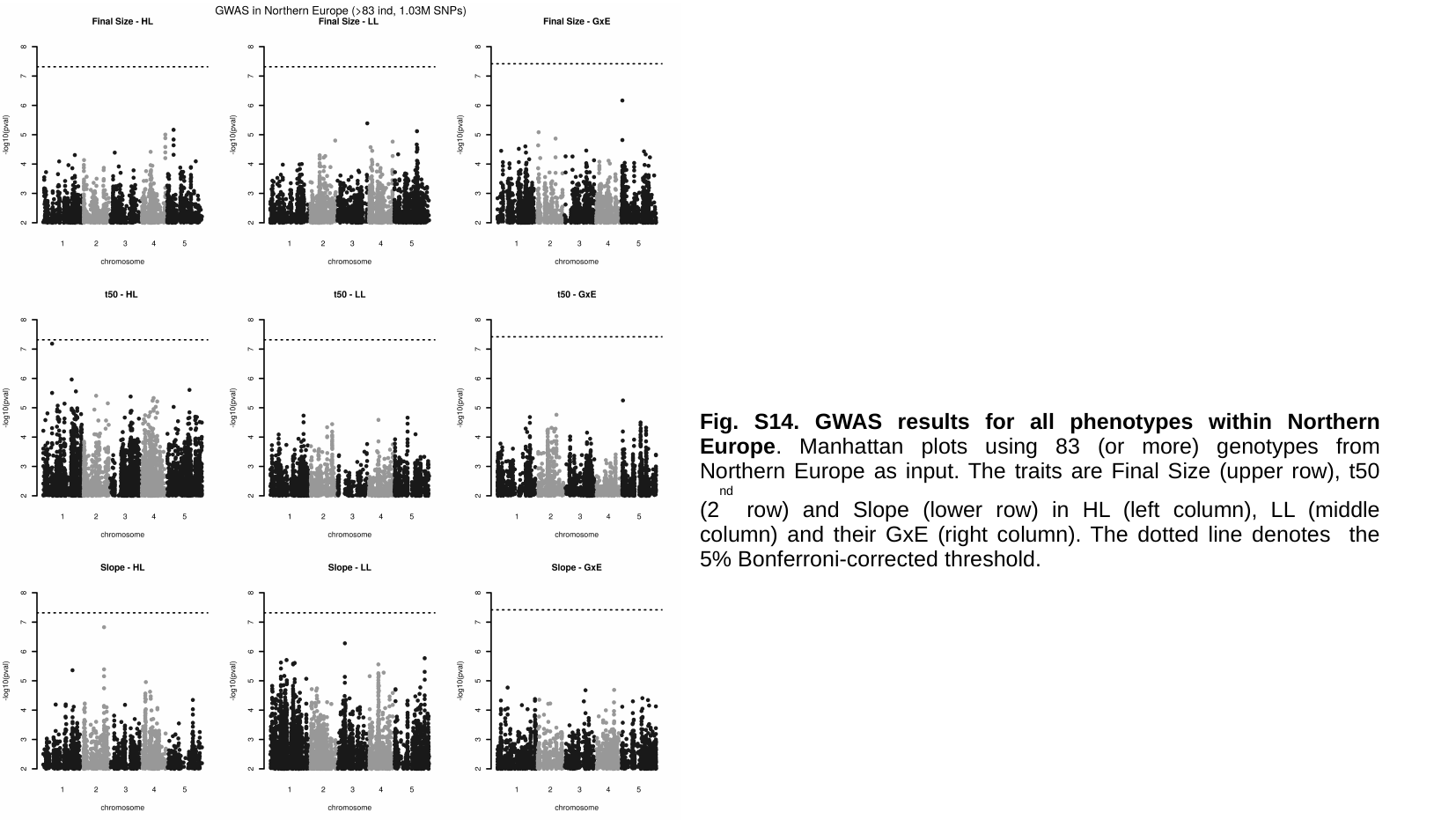

Fig. S14. GWAS results for all phenotypes within Northern Europe. Manhattan plots using 83 (or more) genotypes from Northern Europe as input. The traits are Final Size (upper row), t50 (2^nd^ row) and Slope (lower row) in HL (left column), LL (middle column) and their GxE (right column). The dotted line denotes the 5% Bonferroni-corrected threshold.

### Slide 16
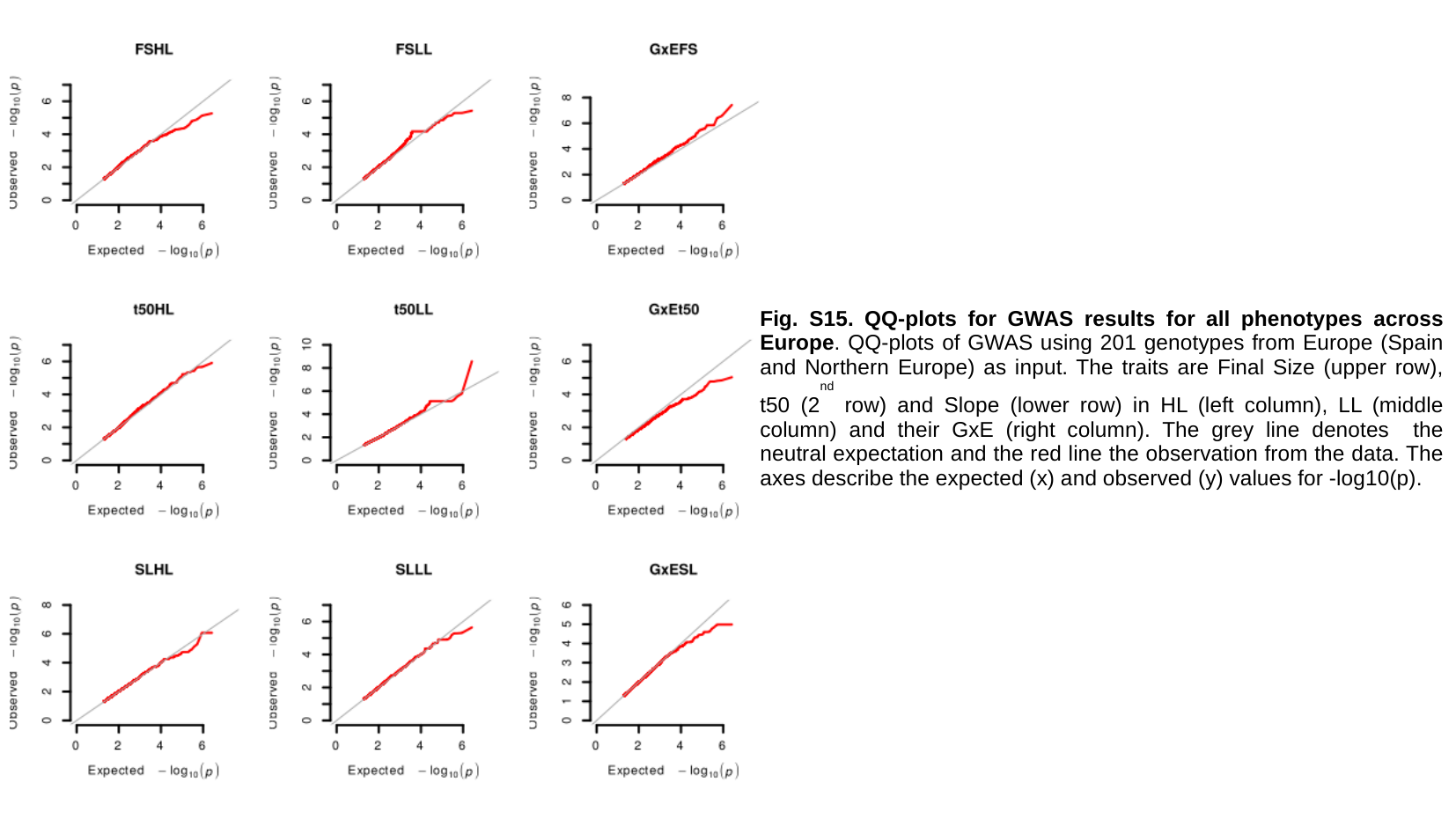

Fig. S15. QQ-plots for GWAS results for all phenotypes across Europe. QQ-plots of GWAS using 201 genotypes from Europe (Spain and Northern Europe) as input. The traits are Final Size (upper row), t50 (2^nd^ row) and Slope (lower row) in HL (left column), LL (middle column) and their GxE (right column). The grey line denotes the neutral expectation and the red line the observation from the data. The axes describe the expected (x) and observed (y) values for -log10(p).

### Slide 17
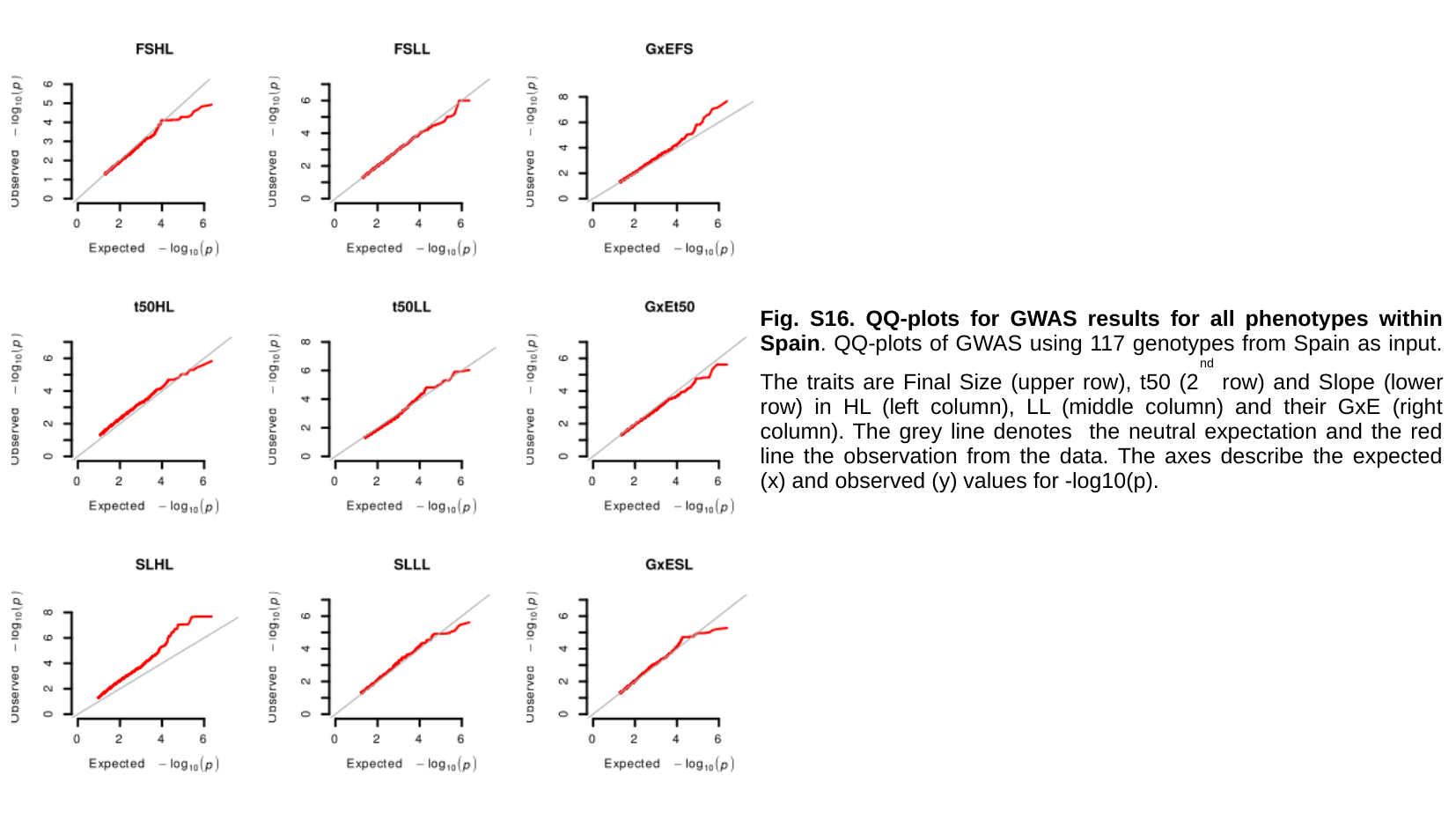

Fig. S16. QQ-plots for GWAS results for all phenotypes within Spain. QQ-plots of GWAS using 117 genotypes from Spain as input. The traits are Final Size (upper row), t50 (2^nd^ row) and Slope (lower row) in HL (left column), LL (middle column) and their GxE (right column). The grey line denotes the neutral expectation and the red line the observation from the data. The axes describe the expected (x) and observed (y) values for -log10(p).

### Slide 18
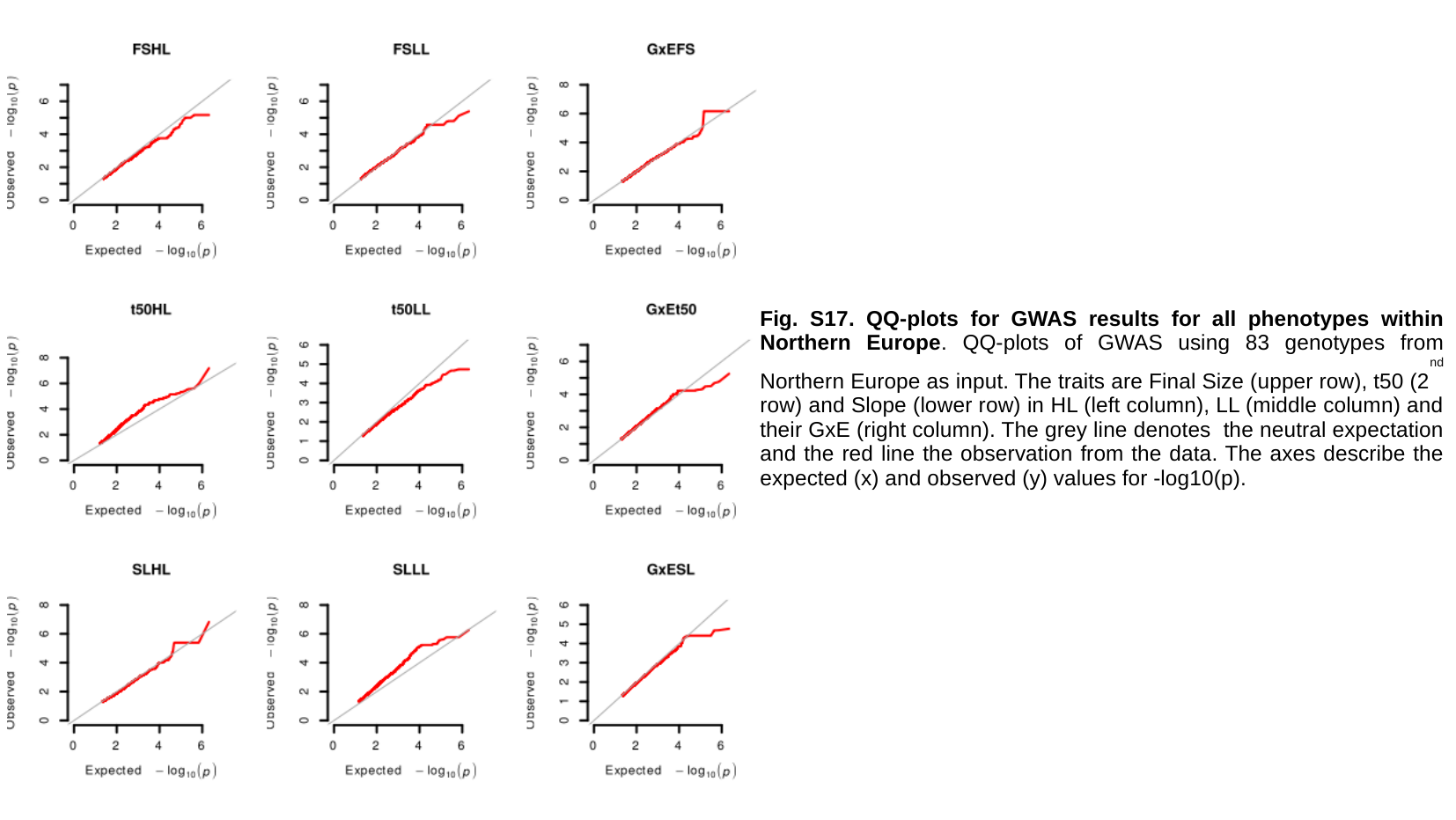

Fig. S17. QQ-plots for GWAS results for all phenotypes within Northern Europe. QQ-plots of GWAS using 83 genotypes from Northern Europe as input. The traits are Final Size (upper row), t50 (2^nd^ row) and Slope (lower row) in HL (left column), LL (middle column) and their GxE (right column). The grey line denotes the neutral expectation and the red line the observation from the data. The axes describe the expected (x) and observed (y) values for -log10(p).

### Slide 19
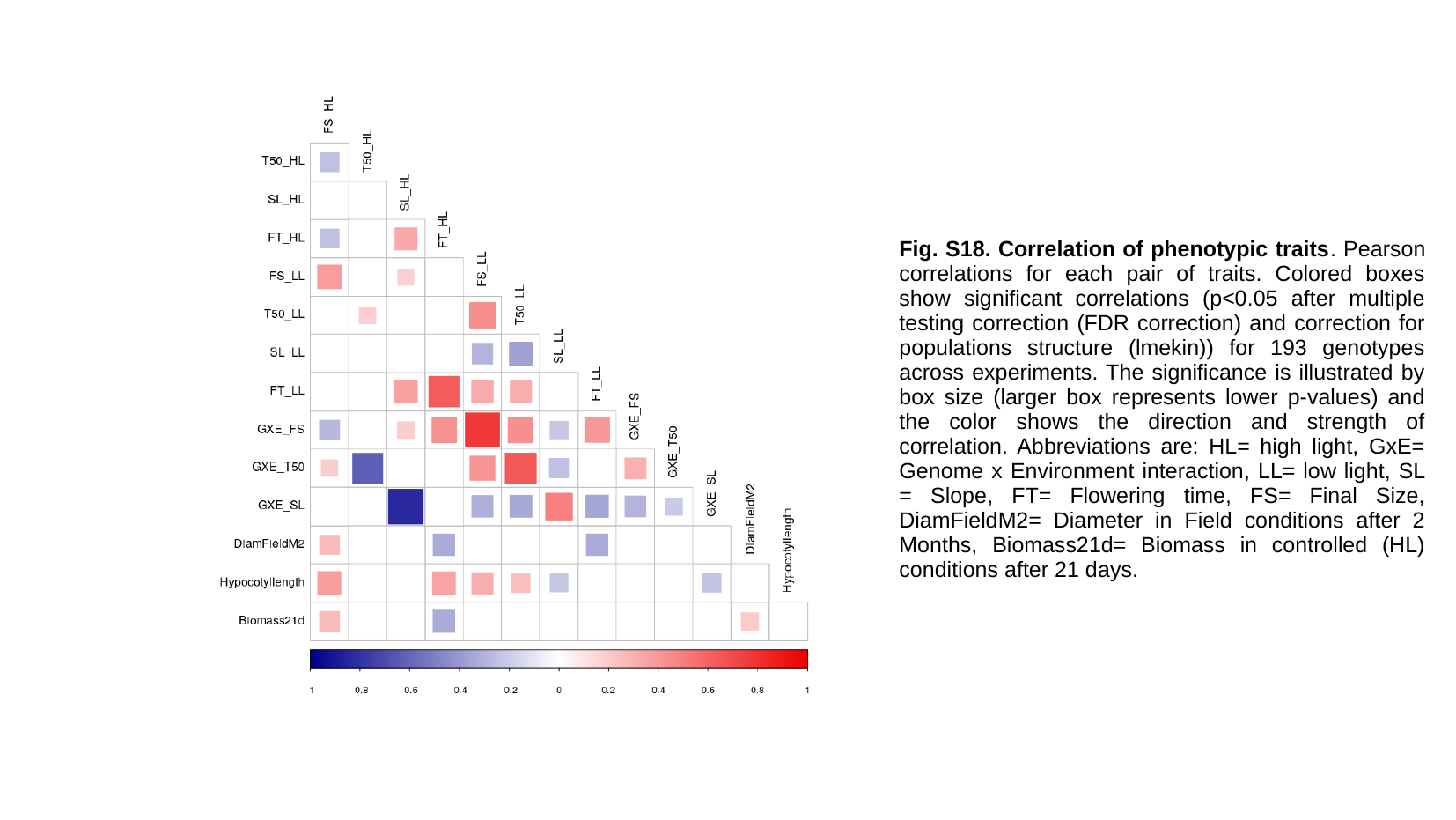

Fig. S18. Correlation of phenotypic traits. Pearson correlations for each pair of traits. Colored boxes show significant correlations (p<0.05 after multiple testing correction (FDR correction) and correction for populations structure (lmekin)) for 193 genotypes across experiments. The significance is illustrated by box size (larger box represents lower p-values) and the color shows the direction and strength of correlation. Abbreviations are: HL= high light, GxE= Genome x Environment interaction, LL= low light, SL = Slope, FT= Flowering time, FS= Final Size, DiamFieldM2= Diameter in Field conditions after 2 Months, Biomass21d= Biomass in controlled (HL) conditions after 21 days.

### Slide 20
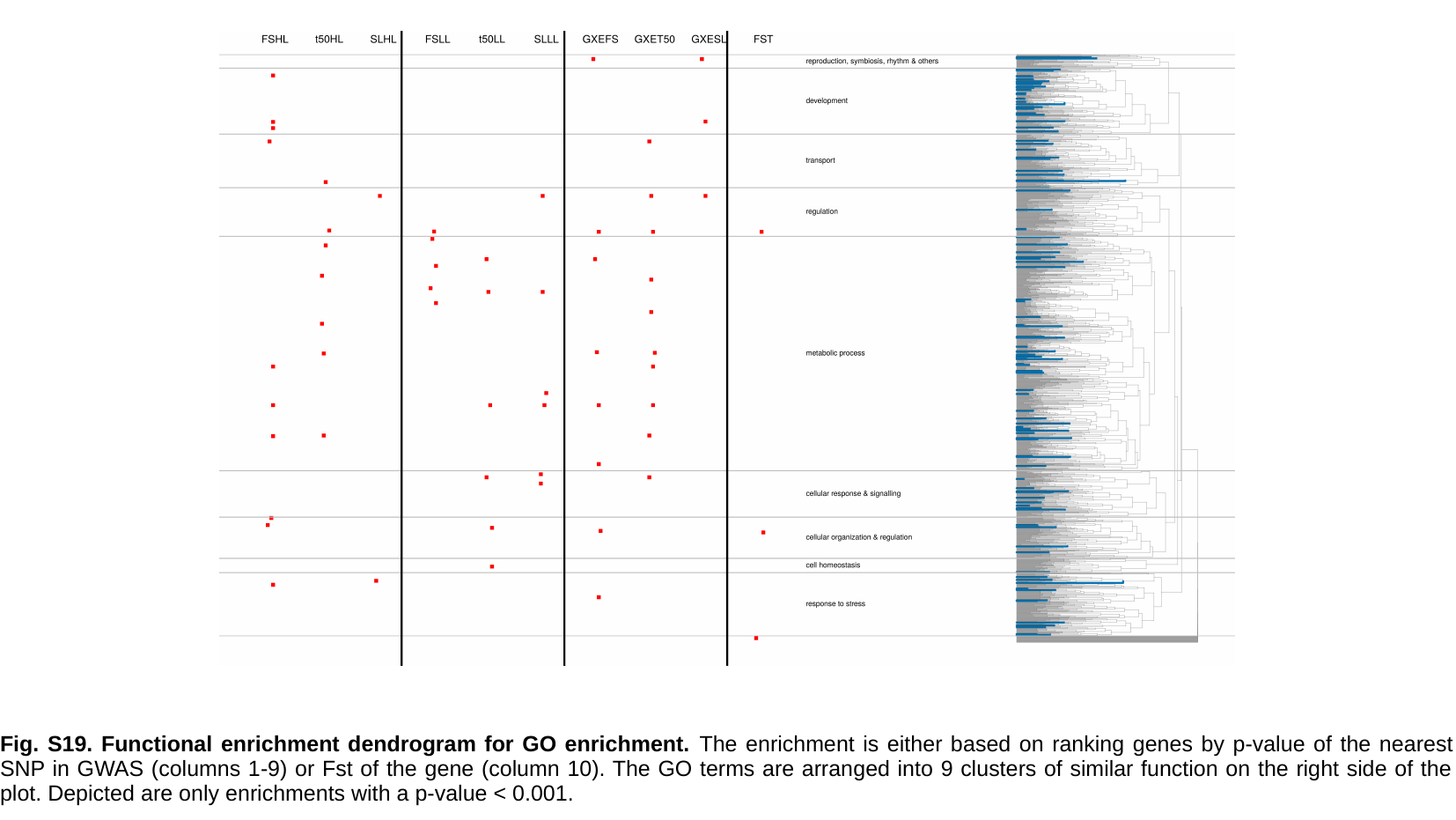

Fig. S19. Functional enrichment dendrogram for GO enrichment. The enrichment is either based on ranking genes by p-value of the nearest SNP in GWAS (columns 1-9) or Fst of the gene (column 10). The GO terms are arranged into 9 clusters of similar function on the right side of the plot. Depicted are only enrichments with a p-value < 0.001.

### Slide 21
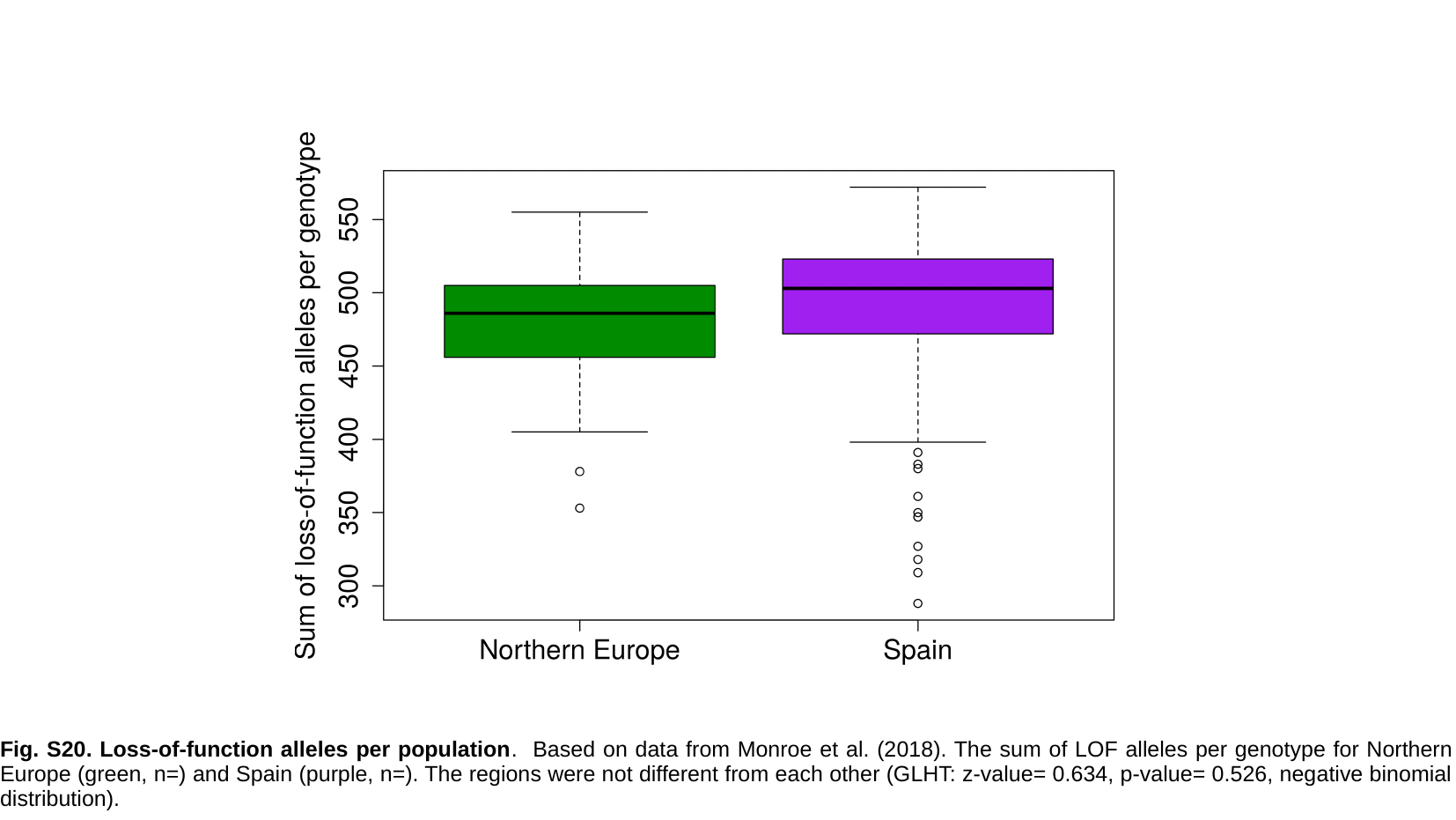

Fig. S20. Loss-of-function alleles per population. Based on data from Monroe et al. (2018). The sum of LOF alleles per genotype for Northern Europe (green, n=) and Spain (purple, n=). The regions were not different from each other (GLHT: z-value= 0.634, p-value= 0.526, negative binomial distribution).

### Slide 22
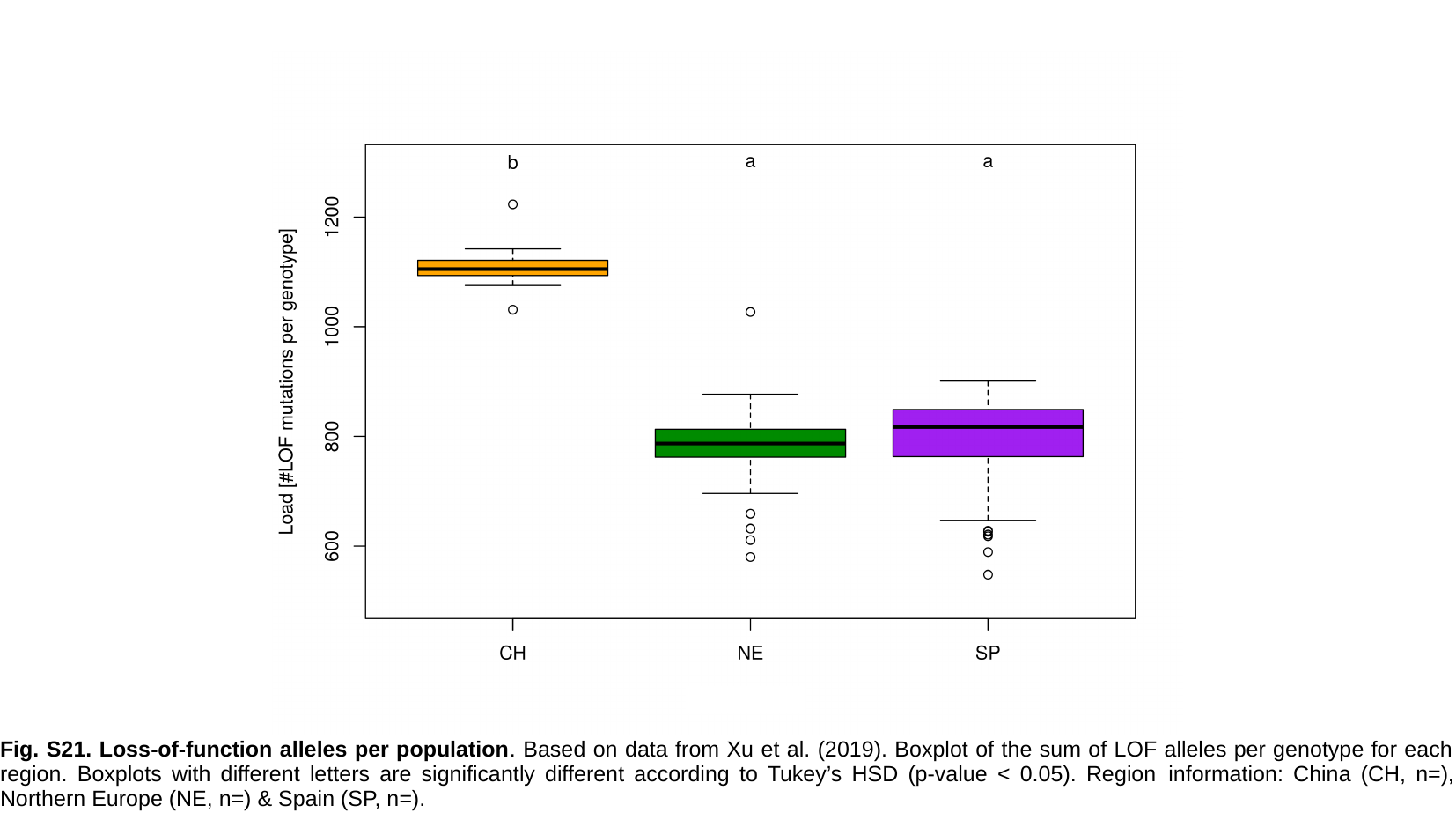

Fig. S21. Loss-of-function alleles per population. Based on data from Xu et al. (2019). Boxplot of the sum of LOF alleles per genotype for each region. Boxplots with different letters are significantly different according to Tukey’s HSD (p-value < 0.05). Region information: China (CH, n=), Northern Europe (NE, n=) & Spain (SP, n=).

### Slide 23
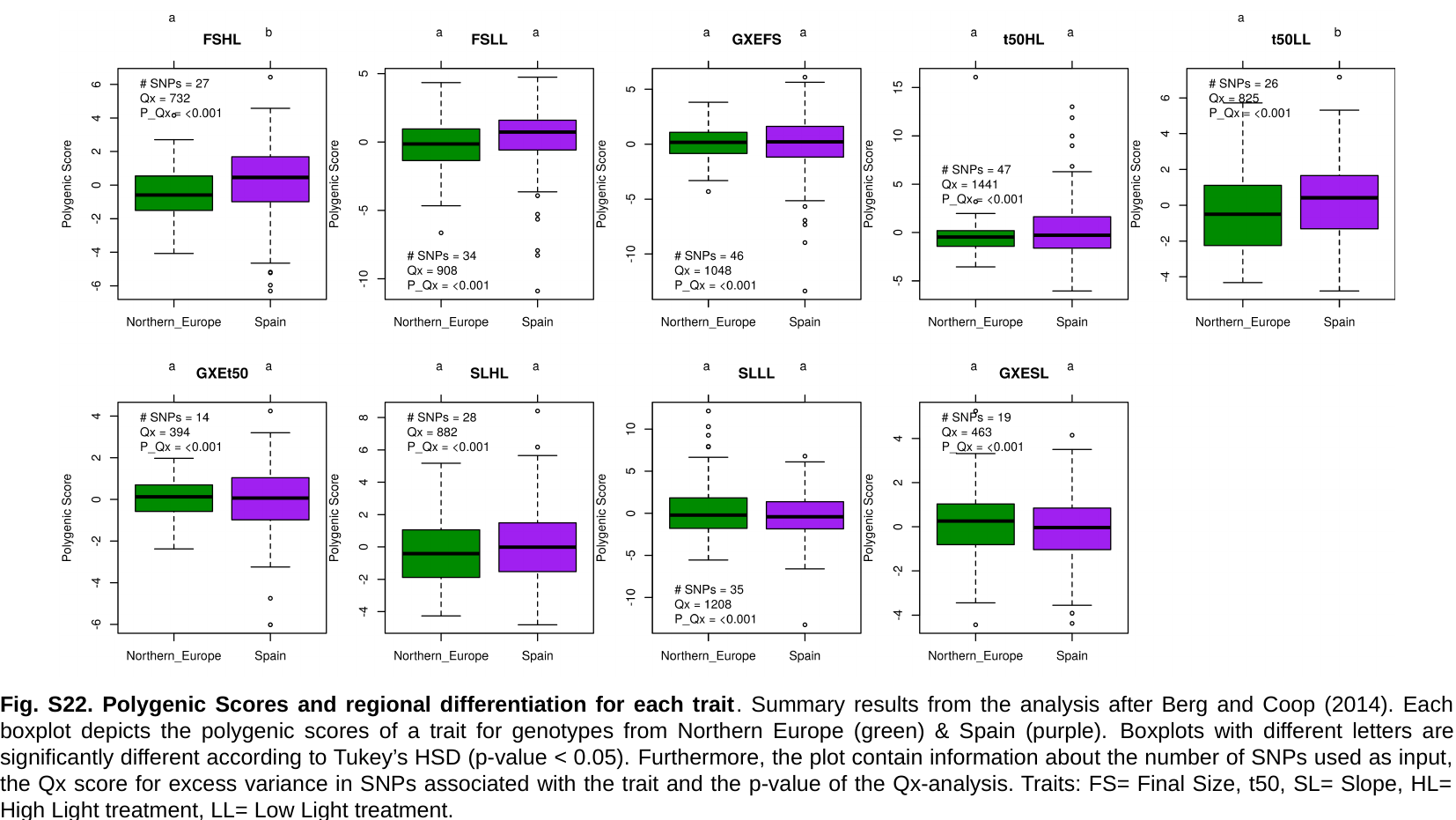

Fig. S22. Polygenic Scores and regional differentiation for each trait. Summary results from the analysis after Berg and Coop (2014). Each boxplot depicts the polygenic scores of a trait for genotypes from Northern Europe (green) & Spain (purple). Boxplots with different letters are significantly different according to Tukey’s HSD (p-value < 0.05). Furthermore, the plot contain information about the number of SNPs used as input, the Qx score for excess variance in SNPs associated with the trait and the p-value of the Qx-analysis. Traits: FS= Final Size, t50, SL= Slope, HL= High Light treatment, LL= Low Light treatment.

### Slide 24
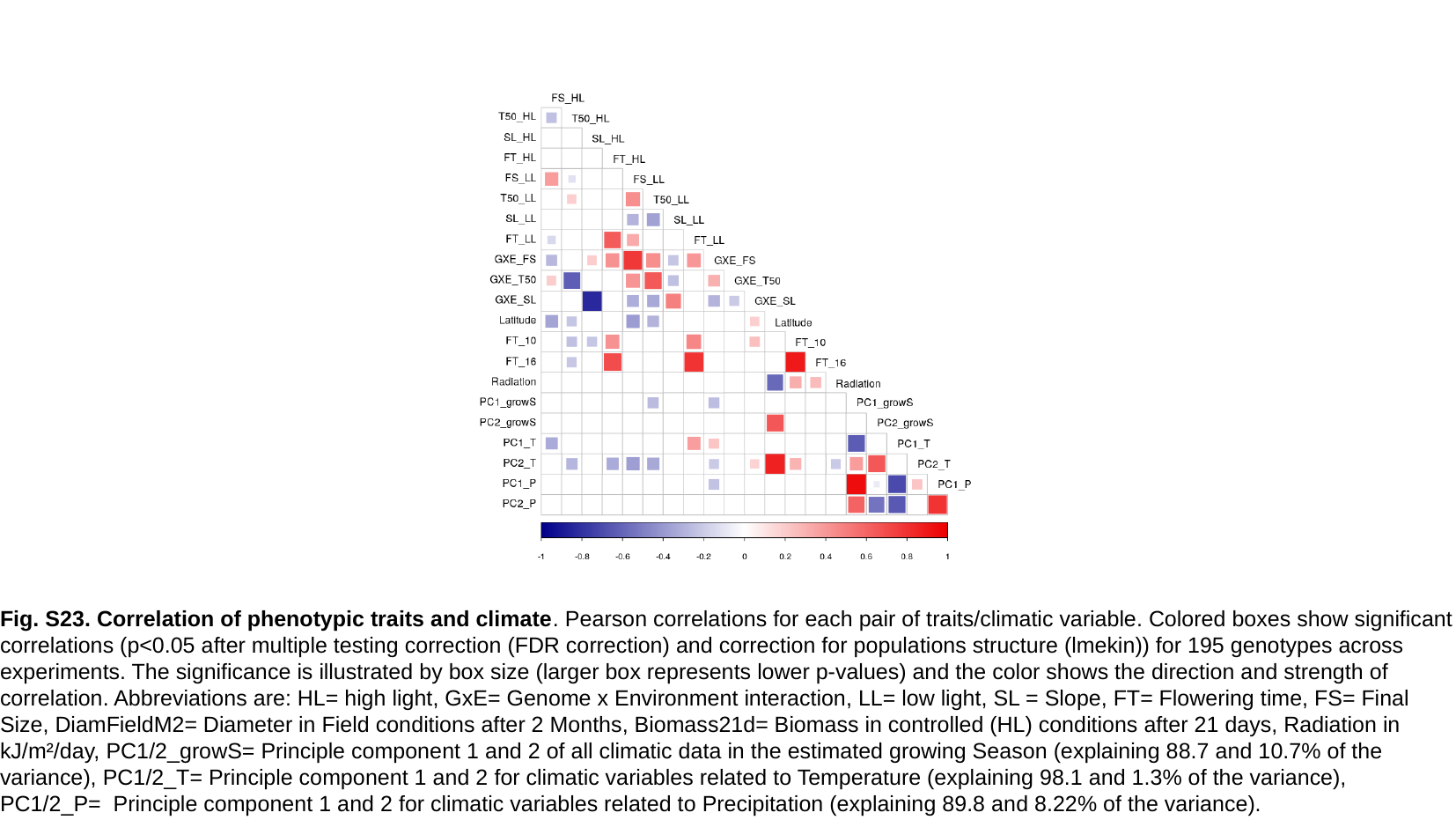

Fig. S23. Correlation of phenotypic traits and climate. Pearson correlations for each pair of traits/climatic variable. Colored boxes show significant correlations (p<0.05 after multiple testing correction (FDR correction) and correction for populations structure (lmekin)) for 195 genotypes across experiments. The significance is illustrated by box size (larger box represents lower p-values) and the color shows the direction and strength of correlation. Abbreviations are: HL= high light, GxE= Genome x Environment interaction, LL= low light, SL = Slope, FT= Flowering time, FS= Final Size, DiamFieldM2= Diameter in Field conditions after 2 Months, Biomass21d= Biomass in controlled (HL) conditions after 21 days, Radiation in kJ/m²/day, PC1/2_growS= Principle component 1 and 2 of all climatic data in the estimated growing Season (explaining 88.7 and 10.7% of the variance), PC1/2_T= Principle component 1 and 2 for climatic variables related to Temperature (explaining 98.1 and 1.3% of the variance), PC1/2_P= Principle component 1 and 2 for climatic variables related to Precipitation (explaining 89.8 and 8.22% of the variance).
